# A hypothalamus-liver-skeletal muscle axis controlled by JNK1 and FGF21 mediates olanzapine-induced insulin resistance in an intraperitoneal treatment in male mice

**DOI:** 10.64898/2026.01.12.698937

**Authors:** Vítor Ferreira, Cintia Folgueira, Ana B. Hitos, Ángela Montes San-Lorenzo, Ánxela Estévez-Salguero, Roger J Davis, Miguel López, Guadalupe Sabio, Patricia Rada, Ángela M. Valverde

## Abstract

**Background:** Olanzapine (OLA), a widely prescribed second-generation antipsychotic, is associated with adverse metabolic effects. We recently showed that oral OLA treatment in male mice induces weight gain and hepatic steatosis, whereas intraperitoneal (i.p.) administration leads to weight loss due to higher hypothalamic OLA levels and activation of brown adipose tissue. Since clinical studies report insulin resistance in individuals treated with OLA, here we investigated the impact of OLA i.p. treatment on insulin sensitivity, focusing on the liver– skeletal muscle axis.

**Material and Methods:** Wild-type male mice were treated with OLA (10 mg/kg, i.p.) for 8 weeks or received a single intrahypothalamic injection (15 nmol). Glucose homeostasis parameters were assessed. Mechanistic studies were performed in vagotomized mice, mice lacking JNK in either the hypothalamus or liver, mice overexpressing hepatic FGF21, and PTP1B-deficient mice (PTP1B-KO).

**Results:** OLA i.p. treatment induced systemic insulin resistance, pyruvate intolerance, and reduced insulin signaling in both liver and skeletal muscle. These effects were accompanied by increased hepatic JNK phosphorylation and IRS1 serine phosphorylation. A single intrahypothalamic OLA injection similarly impaired peripheral insulin action and activated hepatic JNK. Deletion of hypothalamic or hepatic JNK1, as well as vagotomy, prevented these defects. OLA reduced hepatic *Fgf21* expression, an effect reversed by hypothalamic JNK1 deletion or vagotomy. Hepatic FGF21 overexpression prevented OLA-induced insulin resistance in skeletal muscle but not in liver. PTP1B-KO mice were protected from all metabolic impairments.

**Conclusion:** Although OLA i.p. treatment prevents weight gain, it decreases peripheral insulin sensitivity through a hypothalamus–liver axis driven by hypothalamic JNK1, which activates hepatic JNK via the vagus nerve, suppresses hepatic FGF21 and ultimately impairs insulin signaling in skeletal muscle. Importantly, the protection conferred by PTP1B deficiency against OLA-induced insulin resistance strongly suggests that targeting PTP1B might prevent metabolic comorbidities in patients under OLA treatment in a personalized manner.

## 1. Introduction

Second generation antipsychotics (SGA) are the mainstay therapy for several psychiatric disorders including schizophrenia, bipolar disease, acute mania and borderline personality disorder [1–3]. A large proportion of patients treated with SGA develop metabolic abnormalities, including weight gain, insulin resistance, hyperglycemia and dyslipidemia [4–6], being olanzapine (OLA) among the prescribed SGA with higher risk of weight gain [7]. A recent large population-based trial reported that the use of antipsychotics increases the risk for type 2 diabetes (T2D) development by 3-4-fold [8], with young patients being particularly vulnerable to antipsychotic-induced glucose intolerance [9] and T2D [10]. Furthermore, a prospective, controlled, open study comparing OLA-treated patients (7.5-20 mg/day) with healthy volunteers revealed that disturbances in glucose homeostasis induced by OLA are mainly due to insulin resistance, since beta cell function was unaltered [11]. Another study with healthy volunteers receiving OLA (10 mg/day) or placebo for 21 days showed hyperinsulinemia and reduced insulin sensitivity during an oral glucose tolerance test (GTT) at day 19, as well as diminished insulin sensitivity during a hyperinsulinemic-euglycemic clamp, supporting the impact of OLA treatment in glucose homeostasis, specifically in insulin sensitivity [12]. While olanzapine is widely prescribed, the molecular basis of its adverse metabolic effects remains unclear. In fact, very little is known about its metabolic effects, particularly those elicited through neuroendocrine axes.

Many studies using preclinical models have investigated the molecular basis of the adverse metabolic effects of SGAs as reviewed by us and others [13–15], with rodents commonly being used to this end. The most robust and reproducible results of these studies are increases in body weight and metabolic comorbidities induced by SGAs [14, 16–22]. However, rodents do not currently fully recapitulate the human phenotypes [23–26]. Despite these limitations, rat and mouse models remain the best option for the in-depth study of the molecular mechanisms behind the adverse metabolic effects induced by these drugs, including those related to glucose metabolism. Wang *et al.* reported insulin resistance in female rats receiving an oral treatment with OLA (1.5 mg/kg/day) for 8 weeks [27]. This effect was attributed to a defective brown adipose tissue (BAT) thermogenic function evidenced by reductions in both uncoupling protein 1 (UCP-1) and glucose transporter 4 (GLUT4) expression [27] in this fat depot. White adipose tissue (WAT) has also been implicated in the pathophysiology of OLA-induced insulin resistance in female rats following intraperitoneal (i.p.) injection of OLA (10 mg/kg/day) for 8 weeks, specifically by increasing expression levels of proinflammatory cytokines including IL-1β, IL-6, IL-8 and TNFα in this fat depot and also in plasma which, in turn, activated the nuclear factor kappa B (NFκB) through degradation of its inhibitor IĸBα [28]. This hypothesis was supported by the work of Guo *et al.* showing that OLA (8 mg/kg/day via i.p.) administered to female rats for 8 weeks induces inflammation in plasma and epididymal WAT and, as a consequence, impairs insulin responses [29]. Importantly, metformin successfully reversed OLA-induced macrophage infiltration and polarization in WAT, an effect accompanied by attenuation of insulin resistance [29]. Additionally, OLA (5 µM) induced insulin resistance in human preadipocytes co-cultured with macrophages [30]. Likewise, intragastric OLA administration in male rats by continuous infusion (3 mg/kg/h for 160 min) revealed hepatic and extra-hepatic insulin resistance [31]. Other researchers have shown that the oral treatment of male rats with OLA (5 mg/kg/day) for 8 weeks increased the phosphorylation of insulin receptor substrate-1 (IRS1) at serine residues, a molecular hallmark of impaired insulin signaling, in the liver [32]. Moreover, direct treatment of L6 myotubes with OLA (10-100 µM) reduced glycogen content, IRS1 tyrosine phosphorylation and its association with phosphatidylinositol 3-kinase (PI3K), as well as downstream AKT and Glycogen Synthase Kinase-3 (GSK-3) phosphorylation, supporting diminished insulin signaling by OLA in muscle cells [33]. These studies highlight that the impact of OLA in insulin sensitivity must be investigated both as a direct and tissue-specific response, as well as systemically as a result of an interactome between relevant metabolic organs.

Protein tyrosine phosphatase 1B (PTP1B), a negative regulator of leptin and insulin signaling, is currently considered a therapeutic target for obesity and T2D due to the results of many preclinical studies conducted in global or tissue-specific mice deficient in *Ptpn1,* gene encoding PTP1B [34–37]. Regarding insulin sensitivity, the inhibition or deletion of PTP1B protected against insulin resistance in several contexts such as aging [35] or high fat diet (HFD) [38]. Recently, our group has reported a potential therapeutic advantage of the administration of OLA via i.p. when compared to an oral treatment in preventing weight gain and hepatic steatosis in male mice. These beneficial effects were associated with higher OLA levels in plasma and hypothalamus that, on the one hand, prevented weight gain by activating BAT thermogenesis and inguinal WAT (iWAT) browning via inhibition of hypothalamic AMP-activated protein kinase (AMPK) in a PTP1B-independent manner [16] and, on the other, prevented steatosis through a metabolic rewiring in liver metabolism in a c-jun n-terminal kinase 1 (JNK1) and PTP1B-dependent manner [39]. In the latter study, low-grade inflammatory features were found in the hypothalamus of male mice receiving an i.p. treatment with OLA that were attenuated either by deleting JNK1 in this brain region or by using PTP1B-deficient mice. As neuroinflammation is closely linked with insulin resistance and T2D [40], we investigated whether the i.p. administration of OLA could affect insulin sensitivity, particularly in the liver and skeletal muscle and the potential benefits of PTP1B inhibition.

## 2. Methods

Antibodies used for Western blot, primers used for qRT-PCR and antibodies used for immunohistochemistry/immunofluorescence and are listed in Supplementary Tables 1, 2 and 3, respectively. Additional details regarding methods can be found in Supplementary information.

### 2.1. Animals and treatments

Three-months-old WT and PTP1B-knockout (KO) male mice on the C57BL/6J x 129Sv/J genetic background [41], age-matched C57BL/6J, JNK2-KO/JNK1^flox-flox^, liver-specific JNK1/2-KO (L-JNK1/2-KO) and Albumin-Cre male mice were used. Animal studies were approved by the Ethics Committee of Consejo Superior de Investigaciones Científicas (CSIC, Spain) and conducted in accordance with the guidelines for animal care of Comunidad de Madrid and Directive 2010/63/EU (PROEX 007/19, PROEX 215/18 and CNIC-07/18 (PROEX 215/18). Animals were maintained at 22-24 °C and 55 % humidity on 12 h light/dark cycles (starting at 8 am) and fed a regular rodent chow diet (A04, Panlab, Barcelona, Spain) and tap water *ad libitum*.

#### Animal models

Three-months-old WT and PTP1B-knockout (KO) male mice on the C57BL/6J x 129Sv/J genetic background [41] and age-matched C57BL/6J mice were used. In order to generate mice lacking JNK1 and JNK2 in the hypothalamus, JNK2-KO/JNK1^flox-flox^ mice on the C57BL/6J background were injected adeno-associated viruses encoding Cre recombinase (AAV-Cre, SigmaGen Laboratories, Rockville, MD, USA) in the ventromedial nucleus of the hypothalamus (VMH) using the coordinates: 1.46 mm posterior, ± 0.5 mm lateral and 5.5 mm depth to Bregma [42] (1 µl/injection site), as previously reported [16, 43–45]. These mice were referred as AAV-Cre mice. In parallel, JNK2-KO/JNK1^flox-flox^ mice were injected AAV-GFP (Viraquest Inc., North Liberty, IA, USA) as controls (referred as AAV-GFP mice). In order to generate mice with specific deletion of JNK1/2 in the liver (L-JNK1/2-KO), JNK2-KO/JNK1*^flox/flox^* animals were crossed with mice expressing Cre recombinase under the control of the mouse albumin promote (Albumin Cre (B6.Cg Tg(Alb cre)21Mgn/J) mice; Jackson Laboratory). Alb-Cre mice were used as controls. To overexpress FGF21 in the liver, C57BL/6J mice were injected 1 × 10^11^ viral particles (per mouse) of AAV-FGF21 or AAV-Control (CTRL, expressing GFP) via tail vein, as reported [46].

#### OLA intraperitoneal (i.p.) treatment

WT, PTP1B-KO, AAV-Cre, AAV-GFP, AAV-FGF21 and AAV-CTRL mice received vehicle (VEH) (2 % v/v DMSO in 0.9 % NaCl) or 10 mg/kg/day OLA via i.p. injection (10-12 am) for 8 weeks as reported for injectable treatments [24, 47–50], including two recent studies by our group [16, 39].

#### OLA intrahypothalamic injections

During the light phase of the diurnal cycle (8 am-1 pm), C57BL/6J, Alb-Cre and L-JNK1/2-KO mice were anesthetized with isoflurane for ∼5 min prior to the intrahypothalamic injection and placed in the stereotaxic apparatus. Then, mice received bilaterally a single injection of OLA (15 nmol, dose used in previous studies [16, 39, 51]) or VEH (DMSO)) in the VMH at the aforementioned coordinates [42]. During the injection, mice remained anesthetized with isoflurane. To minimize backflow up the needle track, the needle remained inserted for approximately 3-5 min after injection. Mice were sacrificed at different time points post injection (30 min, 8 h and 48 h).

### 2.2. Data analysis

Statistical analysis was performed with GraphPad Prism version-7.0 (GraphPad Software, San Diego, CA, USA). Data are reported as mean and standard error of the mean (SEM). Comparisons between groups were made using Student’s t-test if 2 groups were considered. If more than 2 groups were studied with one variable taken in consideration (*i.e.,* treatment or genotype) One Way-ANOVA (α = 0.05) was used, with Bonferroni’s post-hoc test carried for multiple comparisons between the groups. If not mentioned in the figure legend, it must be considered that Student’s t-test was used for the presented analysis.

## 3. Results

### 3.1. Treatment of male mice with OLA via i.p. induces peripheral insulin resistance and impairs hepatic insulin signaling

To analyze whole-body glucose homeostasis, male mice received daily i.p. injections of OLA (10 mg/kg) or vehicle (VEH) for 8 weeks as previously reported [25, 26], and glucose, insulin, and pyruvate tolerance tests (GTT, ITT, and PTT, respectively) were performed at the end of treatment. As previously reported [16, 52], body weight loss was observed (Figure 1A). GTT did not reveal glucose intolerance (Figure 1B); however, despite the previously reported body weight loss [25], OLA-injected mice showed peripheral insulin resistance (Figure 1C) and pyruvate intolerance (Figure 1D). Also, an increase in fasting blood glucose was observed in mice receiving OLA compared to VEH controls, although within the physiological range (Figure S1A), without changes in fed blood glucose. Likewise, no differences in fasting or fed plasma insulin levels were found in OLA-treated mice (Figure S1B). At the molecular level, the analysis of hepatic insulin signaling revealed a decrease in insulin-induced phosphorylation of the insulin receptor (IR) and AKT (S473 and T308) in OLA-treated mice when compared with VEH-injected controls (Figure 1E). Defective insulin-induced AKT phosphorylation was confirmed in primary hepatocytes isolated at the end of the OLA i.p. treatment (Figure 1F). This metabolic memory has previously been found in primary hepatocytes from mice fed a high fat diet (HFD) [53] and from *db/db* mice [54]. Moreover, in hepatocytes from mice treated with OLA via i.p., glucose uptake was significantly reduced when compared to the VEH controls (Figure 1G). Overall, our results show that, despite the absence of body weight gain [16] and hepatosteatosis [39], OLA i.p.-treated mice presented systemic insulin resistance and impaired insulin signaling in the liver.

**Figure 1.**
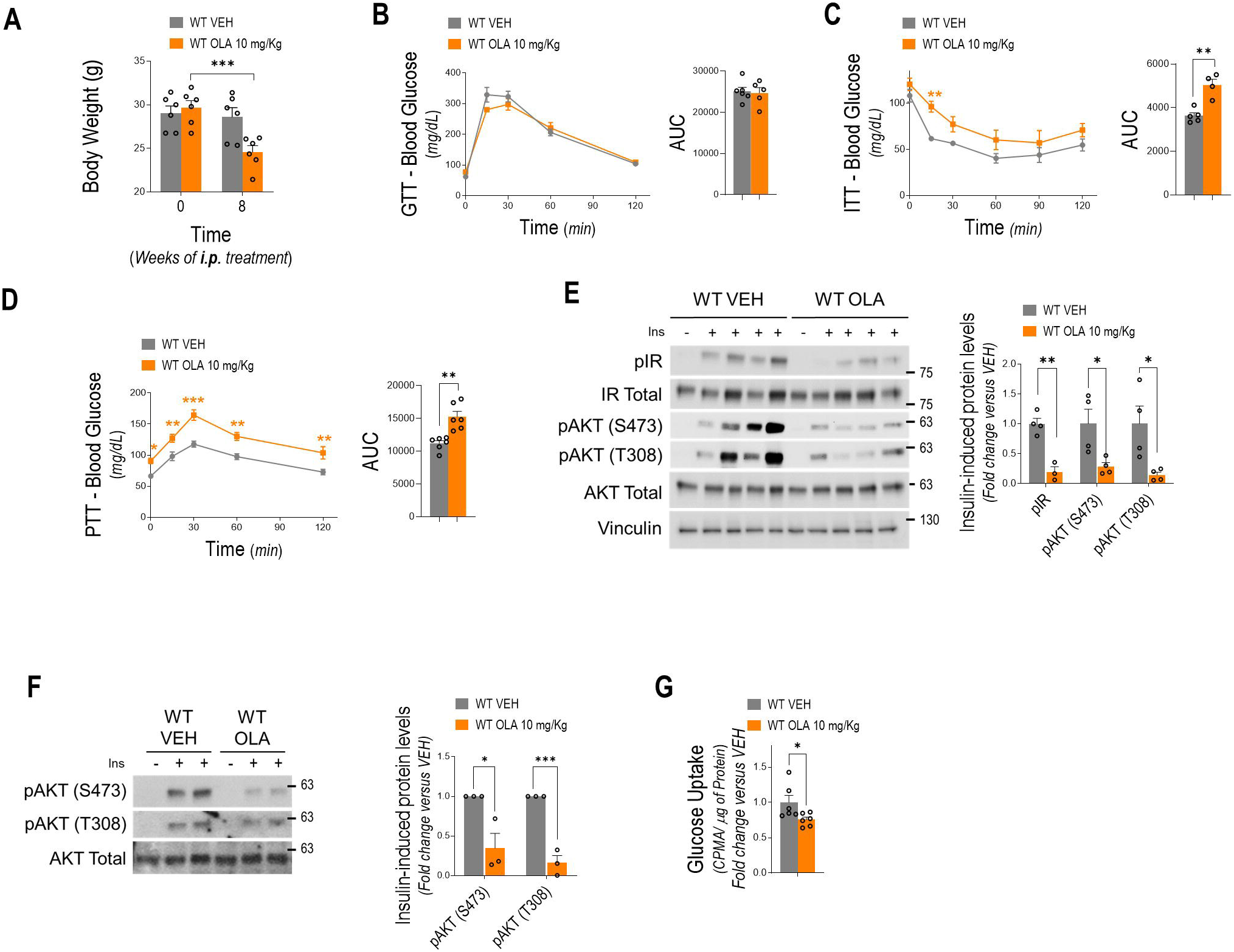
Treatment with OLA via i.p. attenuates systemic and hepatic insulin sensitivity in male mice. WT male mice received a daily i.p. injection of OLA (10 mg/kg) or VEH during 8 weeks. **A**. Body weight (WT VEH: n=6; WT OLA: n=6). **B.** GTT (WT VEH: n=6; WT OLA: n=5), **C.** ITT (WT VEH: n=5; WT OLA: n=4) and **D.** PTT (WT VEH: n=6; WT OLA: n=6) at the end of the treatment and respective area under the curve (AUC) for each group. **E.** Representative Western blots of IR and AKT phosphorylation (Ser473 and Thr308) in liver of mice treated with OLA via i.p. and upon 15 min of insulin (0.75 U/kg) stimulation. Densitometric quantification normalized for total IR, AKT and vinculin levels (WT VEH: n=4; WT OLA: n=4). **F.** Representative Western blots of AKT (Ser473 and Thr308) phosphorylation in primary hepatocytes isolated from male mice treated with VEH or OLA mice and densitometric quantification normalized for total AKT and vinculin levels (WT VEH: n=3; WT OLA: n=3). **G.** Glucose uptake (fold of change versus VEH) of primary hepatocytes from OLA or VEH-treated male mice (WT VEH: n=6; WT OLA: n=6). Each point/bar corresponds to mean ± SEM; comparisons between groups: p < 0.05; p < 0.01; p < 0.001.

### 3.2. OLA i.p. treatment in male mice increases JNK phosphorylation in the liver

Since, on the one hand, the liver is the primary site of OLA metabolism [55] and, on the other, JNK activation has been extensively associated with inflammation, oxidative stress and defective insulin signaling in this organ [56], we next analyzed hepatic JNK phosphorylation under our experimental settings. Figure 2A shows augmented phospho-JNK in the liver of OLA-treated male mice. Interestingly, when primary hepatocytes were treated *in vitro* with OLA at 12.5 µM, concentration that preserves cellular viability [16, 39], activation of JNK was not observed (Figure 2B and S2A), suggesting that OLA-induced JNK phosphorylation in the liver may not be cell autonomous. Furthermore, the immune profile of the liver revealed no alterations by the OLA i.p. treatment in macrophage, neutrophil or T lymphocyte populations as shown by F4/80, MRP14 and CD3 immunostaining of liver sections, respectively (Figure 2C). Of relevance, the number of recruited monocytes positive for Ly6C was slightly increased in the livers of WT mice receiving OLA as assessed by immunofluorescence and immunohistochemistry analysis (Figure 2D). Since, as mentioned above, JNK is activated by reactive oxygen species (ROS), we evaluated features of oxidative stress in livers from male mice receiving OLA via i.p. during 8 weeks and, as shown in Figure 2E-F, no marked differences were found in 4-Hydroxynonenal (4-HNE) staining among the experimental groups. In this line, no differences were found in *Hmox1* mRNA levels (encoding Hemeoxigenase-1) (Figure 2G), pointing that hepatic JNK activation and insulin resistance during OLA treatment are not related to oxidative stress. In line with these features, signs of fibrosis were absent in the livers from OLA-treated mice (Figure 2E-G). These results suggest that the activation of JNK in the liver of WT male mice during OLA i.p. treatment is likely independent of intrahepatic signals.

**Figure 2.**
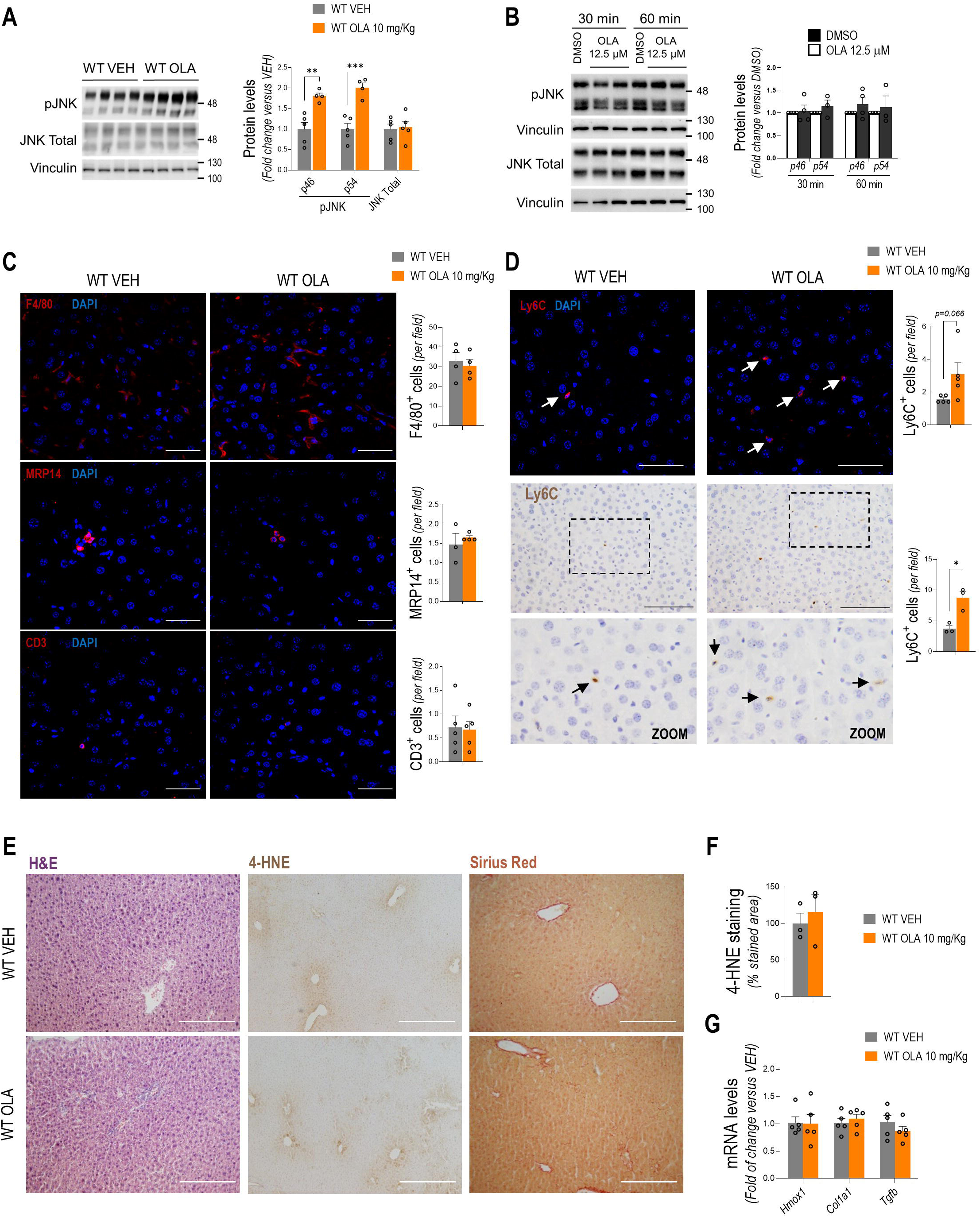
Effect of the treatment with OLA via i.p. in hepatic JNK activation in male mice. **A.** Representative Western blots of JNK phosphorylation in liver of mice treated with OLA via i.p. and densitometric quantification normalized for total JNK and vinculin levels (WT VEH: n=5; WT OLA: n=4). **B.** Representative Western blots of JNK phosphorylation in primary mouse hepatocytes treated *in vitro* with OLA (12.5 µM) for several time periods and densitometric quantification normalized for total JNK and vinculin levels (DMSO: n=4; OLA: n=4). **C.** Representative confocal images of (*upper to lower panel*) F4/80 (WT VEH: n=4; WT OLA: n=4), MRP14 (WT VEH: n=3; WT OLA: n=4) and CD3 (WT VEH: n=5; WT OLA: n=5) immunofluorescence in liver sections (40x, scale bars-50 µm) of mice receiving VEH (*left panel*) or OLA (*right panel*) via i.p. and their respective quantifications. **D.** Representative images of Ly6C immunofluorescence (*upper panel,* 40x, scale bars-50 µm; WT VEH: n=5; WT OLA: n=5) and immunohistochemistry (*down panel,* 40x, scale bars-100 µm; WT VEH: n=3; WT OLA: n=3) analysis in liver sections of mice receiving VEH (*left panel*) or OLA (*right panel*) via i.p. and their respective quantifications. **E.** Representative images of H&E (WT VEH: n=4; WT OLA: n=4), 4-HNE immunohistochemistry (WT VEH: n=3; WT OLA: n=3) and Sirius Red staining (WT VEH: n=4; WT OLA: n=4). **F.** Hepatic 4-HNE staining quantification (WT VEH: n=3; WT OLA: n=3). **G.** Hepatic mRNA levels of *Hmox1*, *Col1a1* and *Tgfb* (WT VEH: n= 5; WT OLA: n= 5), normalized to *Actb* as a housekeeping gene. Each point/bar corresponds to mean ± SEM; comparisons between groups: p < 0.05; p < 0.01; p < 0.001.

### 3.3. Hypothalamic JNK1 controls hepatic insulin signaling through the vagus nerve in male mice treated with OLA via i.p

In line with the absence of cell autonomous effects in response to OLA in hepatocyte JNK activation (Figure 2B and S2A), the pretreatment of primary hepatocytes with 12.5 μM before insulin stimulation (10 nM for 15 min) did not decrease AKT phosphorylation (Figure 3A). Therefore, we hypothesized that a central-peripheral inter-organ crosstalk might control OLA-induced impairments in hepatic insulin signaling. To achieve this, C57BL/6J male mice received an intrahypothalamic injection of OLA and insulin signaling was analyzed in the liver 48 h post-injection. Interestingly, insulin-induced AKT phosphorylation (S473 and T308) was attenuated in the liver of mice receiving OLA via intrahypothalamic injection (Figure 3B). Since it has been extensively reported that JNK phosphorylates IRS1 at serine residues [53] which, in turn, decreases its tyrosine phosphorylation upon insulin stimulation, this cross-talk was analyzed. As shown in Figure 3C, JNK phosphorylation was increased in the liver of mice at 30 min after OLA intrahypothalamic injection. This response paralleled an increase in IRS1 serine phosphorylation (Figure 3D) and, in this line, insulin-induced IRS1 tyrosine phosphorylation was reduced (Figure 3E). Interestingly, elevated IRS1 serine phosphorylation was also found in the liver of mice receiving chronic OLA i.p. treatment (Figure 3F) concurrently with JNK phosphorylation levels (Figure 2A) and, in agreement, those mice showed a decrease in IRS1 tyrosine phosphorylation in response to insulin (Figure 3G).

**Figure 3.**
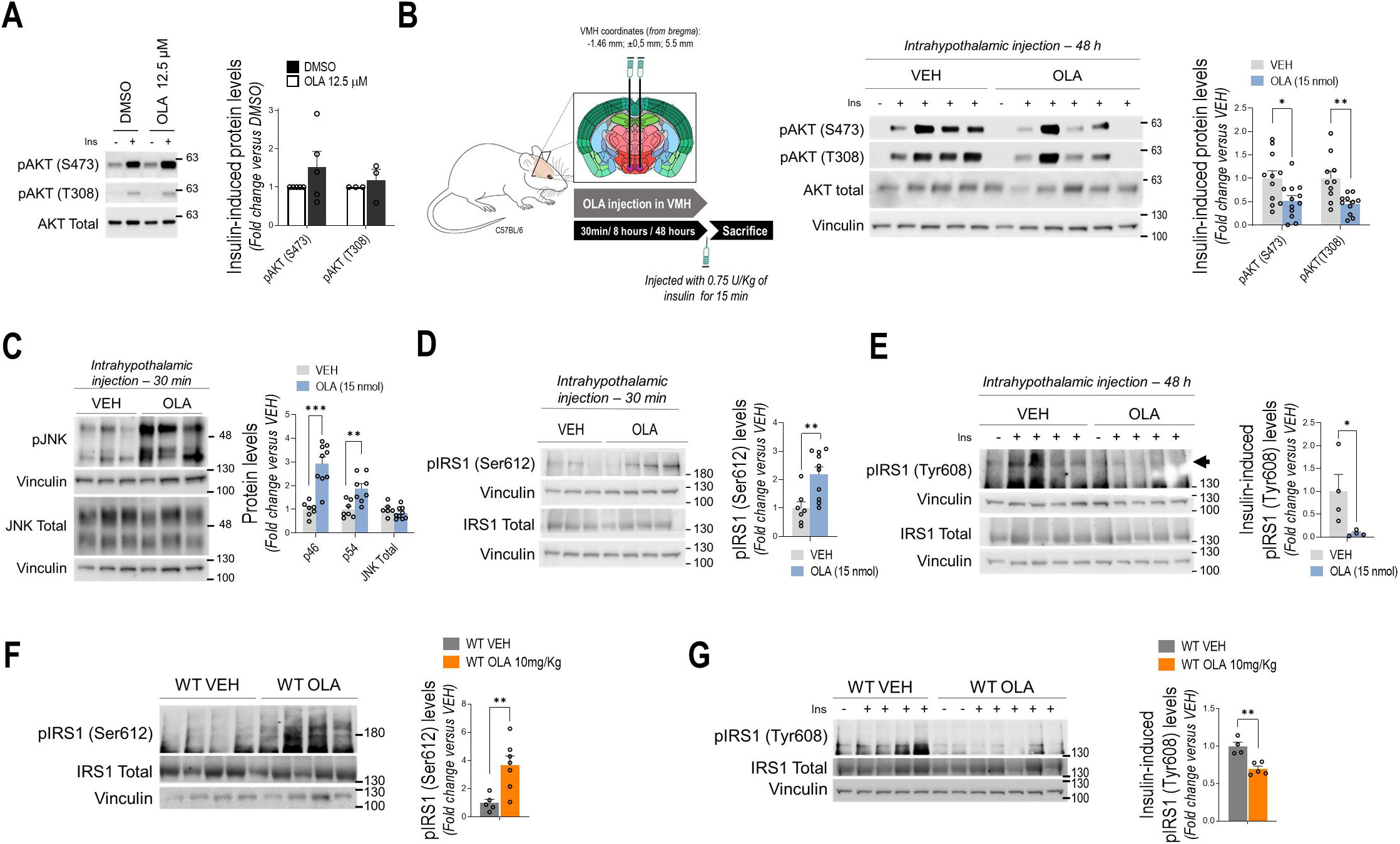
The defective hepatic insulin signaling induced by OLA is associated with an increase in IRS1 serine phosphorylation and a decrease in IRS1 tyrosine phosphorylation in male mice. **A.** Representative Western blots of insulin -induced AKT (Ser473 and Thr308) phosphorylation in primary mouse hepatocytes treated *in vitro* with OLA (12.5 µM) for 48 h prior to insulin stimulation (10 nM, 15 min). Densitometric quantification normalized for total AKT and vinculin levels (DMSO: n=4; OLA: n=4). **B.** Experimental design of OLA-intrahypothalamic injection (*left panel*). Representative Western blots of AKT phosphorylation (Ser473 and Thr308) in liver of mice 48 h after an OLA (15 nmol) intrahypothalamic injection and upon 15 min of insulin (0.75 U/kg) stimulation. Densitometric quantification normalized for total AKT and vinculin levels (DMSO: n=11; OLA: n=13). **C.** Representative Western blots of JNK phosphorylation in liver of mice 30 min after an OLA (15 nmol) intrahypothalamic injection and densitometric quantification normalized for total JNK and vinculin levels (DMSO: n=7; WT OLA: n=9). **D.** Representative Western blots of IRS1 Ser612 phosphorylation in liver of mice 30 min after an OLA (15 nmol) intrahypothalamic injection and densitometric quantification normalized for total IRS1 and vinculin levels (DMSO: n=7; OLA: n=10). **E.** Representative Western blots of insulin-induced IRS1 Tyr608 phosphorylation in liver of mice 48 h after an OLA (15 nmol) intrahypothalamic injection and upon 15 min of insulin (0.75 U/kg) stimulation. Densitometric quantification normalized for total IRS1 and vinculin levels (DMSO: n=4; OLA: n=4). **F.** Representative Western blots of IRS1 Ser612 phosphorylation in liver of mice receiving OLA via i.p. and densitometric quantification normalized for total IRS1 and vinculin levels (WT VEH: n=5; WT OLA: n=7). **G.** Representative Western blots of insulin-induced IRS1 Tyr608 phosphorylation in liver of mice receiving OLA via i.p. upon 15 min of insulin (0.75 U/kg) stimulation. Densitometric quantification normalized for total IRS1 and vinculin levels (WT VEH: n=4; WT OLA: n=5). Each point/bar corresponds to mean ± SEM; comparisons between groups: p < 0.05; p < 0.01; p < 0.001.

As we previously demonstrated that hypothalamic JNK is activated in response to i.p. or central OLA administration [39], and to further investigate the role of hypothalamic JNK1 in hepatic insulin resistance induced by OLA, we conducted experiments in a cohort of male mice in which JNK was deleted specifically in this brain region by injecting AAV-Cre particles in the VMH of JNK2-KO/JNK1*^flox/flox^* mice prior to OLA i.p. treatment (10 mg/kg/day) for 8 weeks, as we previously reported [39]. Figures 4A and S3A show that deletion of JNK1/2 isoforms in the hypothalamus prevented OLA-induced attenuation of AKT (S473 and T308) phosphorylation in the liver. Surprisingly, insulin-induced AKT phosphorylation was enhanced in mice with deletion of hypothalamic JNK receiving OLA treatment (Figure S3A). Moreover, deletion of hypothalamic JNK1/2 reduced OLA-induced JNK phosphorylation in the liver, an effect accompanied by a decrease in IRS1 serine phosphorylation, thereby protecting against the reduction of insulin-induced IRS1 tyrosine phosphorylation by this SGA (Figure 4B-D). These data envision hypothalamic JNK1 as a key controller of hepatic insulin signaling in response to OLA i.p. treatment.

**Figure 4.**
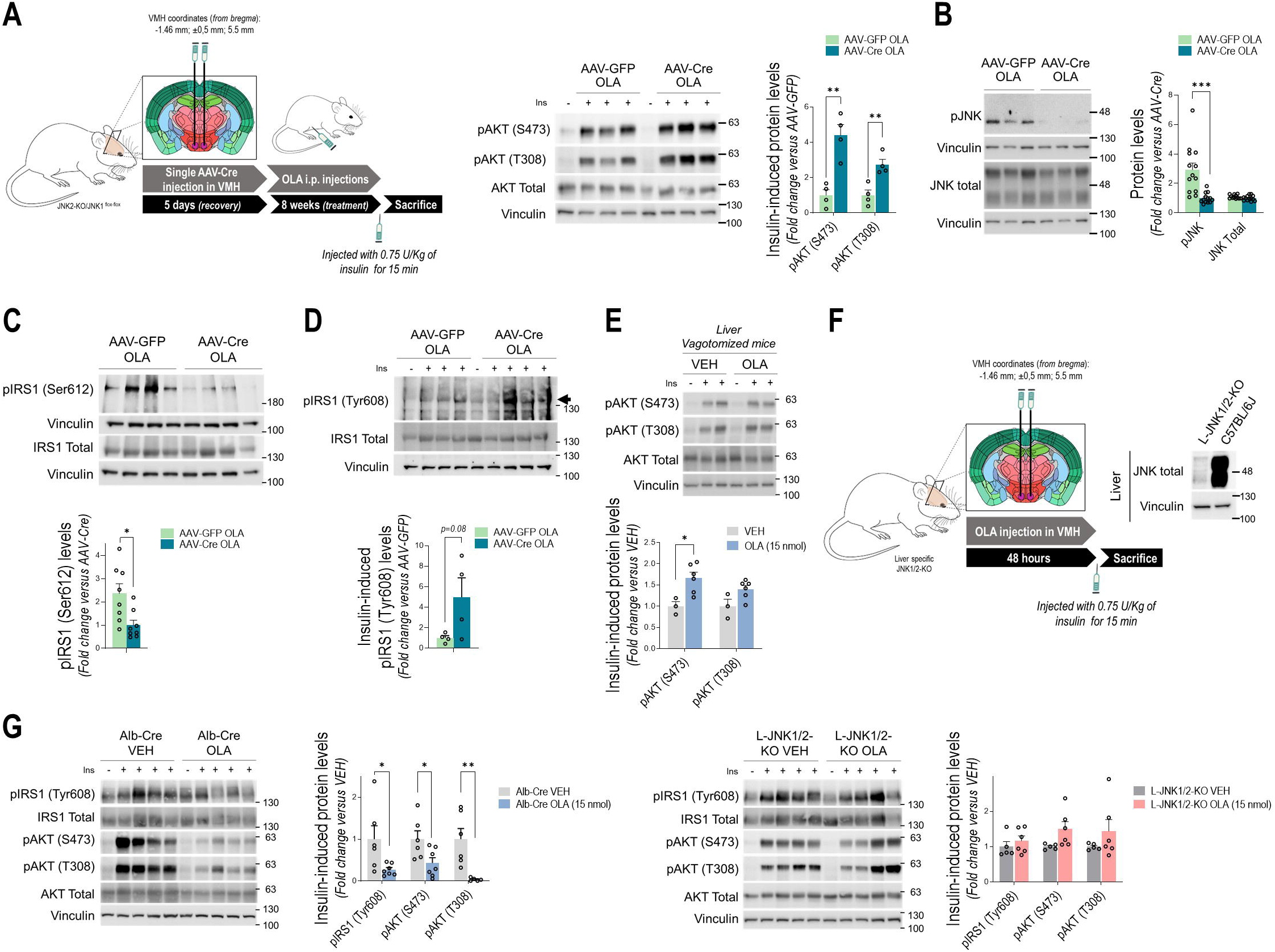
Hypothalamic or hepatic JNK1 deletion prevents OLA-induced attenuation in hepatic insulin signaling in male mice. **A.** Experimental design of the intrahypothalamic injection of adeno-associated viruses encoding Cre recombinase in C57BL/6J male JNK2-KO/JNK1^flox-flox^ mice 5 days prior to OLA i.p. daily injections for 8 weeks. Representative Western blots of insulin-induced AKT phosphorylation (Ser473 and Thr308) in liver of mice with hypothalamic JNK1 deletion (AAV-Cre) or AAV-GFP control mice receiving OLA via i.p. Densitometric quantification normalized for total AKT and vinculin levels (AAV-GFP OLA: n=4; AAV-Cre OLA: n=4). **B.** Representative Western blots of JNK phosphorylation in liver of AAV-Cre or AAV-GFP control mice treated with OLA via i.p. and densitometric quantification normalized for total JNK and vinculin levels (AAV-GFP OLA: n=13; AAV-Cre OLA: n=13). **C.** Representative Western blots of IRS1 Ser612 phosphorylation in liver of OLA-treated AAV-Cre or AAV-GFP control mice and densitometric quantification normalized for total IRS1 and vinculin levels (AAV-GFP OLA: n=8; AAV-Cre OLA: n=8). **D.** Representative Western blots of insulin-induced IRS1 Tyr608 phosphorylation in liver of OLA-treated AAV-Cre or AAV-GFP control mice receiving an insulin (0.75 U/kg) injection for 15 min. Densitometric quantification normalized for total IRS1 and vinculin levels (AAV-GFP OLA: n=4; AAV-Cre OLA: n=4).). **E.** Representative Western blots of insulin-induced AKT phosphorylation (Ser473 and Thr308) in liver of vagotomized mice 48 h after an OLA (15 nmol) intrahypothalamic injection and treated with insulin (0.75 U/kg, 15 min), and densitometric quantification normalized for total AKT and vinculin levels (DMSO: n=3; OLA: n=6). **F.** Experimental design of the intrahypothalamic injection of OLA (15 nmol) in a liver-specific JNK1/2-KO model (L-JNK1/2-KO) (*left panel*). Representative Western blot images of hepatic total JNK and Vinculin protein levels (*right panel*) to assess JNK1/2 deletion in the liver. **G.** Representative Western blots of insulin-induced IRS1 phosphorylation in Tyr608 residues and AKT phosphorylation (S473 and T308) in liver of Alb-Cre mice (*left panel*) and L-JNK1/2-KO (*right panel*) 48 h after receiving an OLA intrahypothalamic injection and an insulin (0.75 U/kg) injection for the last 15 min. Densitometric quantification normalized for total IRS1, AKT and vinculin levels (Alb-Cre DMSO: n= 6; Alb-Cre OLA: n= 7; L-JNK1/2-KO DMSO: n=5; L-JNK1/2-KO OLA: n=6). Each point/bar corresponds to mean ± SEM; comparisons between groups: p < 0.05; p < 0.01; p < 0.001.

In our previous study we found that OLA-mediated signals modulating hepatic lipid metabolism originate from the hypothalamus and are transmitted to the liver via the vagus nerve [39]. To further investigate the relevance of the vagal signals to the hypothalamic-driven effects of OLA in hepatic insulin signaling, OLA or VEH was injected into the VMH of vagotomized C57BL/6J mice. The effectiveness of the vagotomy was assessed at the end of the study through a post-mortem analysis of the stomach. Only mice that showed a clear increase in stomach size following vagotomy due to motor dysfunction (Figure S3B) were included in the analysis. As shown in Figure 4E, intrahypothalamic injection of OLA did not attenuate AKT phosphorylation in the liver of vagotomized mice. Indeed, AKT S473 phosphorylation was increased. These results point to a central-peripheral communication via the vagus nerve in this response. In this line, our results suggest that OLA-induced insulin resistance in the liver likely results from an inter-organ crosstalk dependent on hypothalamic signals that promote the activation of hepatic JNK which, in turn phosphorylates IRS1 on serine, a well-known molecular signature of defective insulin signaling.

### 3.4. Liver-specific JNK1 deletion prevents the impairment of insulin signaling induced by OLA i.p. treatment

Since we found, on the one hand, increased JNK phosphorylation in the liver of male mice receiving OLA either by i.p. or intrahypothalamic injection (Figure 2A, 3C), both of which exhibit hepatic insulin resistance (Figure 1E, 3B), and on the other, decreased hepatic JNK activation in mice lacking JNK1 in the hypothalamus (Figure 4B) which is associated with protection against OLA-induced impairment in insulin signaling (Figure 4A), we postulated that central signals driven by OLA activate hepatic JNK1 which is required to impair insulin signaling at the level of IRS1 tyrosine phosphorylation. To address this, we performed experiments in a cohort of mice in which JNK1 was specifically deleted in the liver of JNK2-KO/JNK1*^flox/flox^* mice, as detailed in Material and Methods. As shown in Figure 4F, both isoforms were absent in the liver of JNK2-KO/JNK1*^AlbCre^* (L-JNK1/2-KO) animals. Next, hepatic insulin signaling was analyzed 48 h after an intrahypothalamic OLA injection. As shown in Figure 4G, L-JNK1/2-KO mice receiving an intrahypothalamic injection of OLA retained the response to insulin in inducing IRS1 (Tyr608) and AKT (S473 and T308) phosphorylation in the liver similar to their vehicle treated-controls. Additionally, the response of the livers from L-JNK1/2-KO mice regarding AKT phosphorylation was compared with that of Alb-Cre control mice receiving an intrahypothalamic injection of OLA or VEH. As shown in Figure S4A, no differences in insulin-induced AKT phosphorylation were observed between the VEH groups, whereas L-JNK1/2-KO mice showed increased AKT phosphorylation in the liver upon OLA injection compared to their control Alb-Cre mice. Overall, these findings emphasize the pivotal role of hepatic JNK in mediating OLA-induced deficits in insulin signaling in the liver.

### 3.5. OLA administration via i.p. reduces hepatic and circulating FGF21 in male mice

Since, skeletal muscle accounts for around 85% of the glucose usage during an euglycemic-hyperinsulinemic clamp [57, 58], we evaluated insulin signaling in skeletal muscle in male mice treated with OLA via i.p. injection during 8 weeks. As shown in Figure 5A, a significant decrease in AKT phosphorylation was found in the skeletal muscle of OLA-treated mice. Skeletal muscle insulin resistance was corroborated in mice (Alb-Cre) receiving a single intrahypothalamic injection of OLA since they also showed reduced AKT phosphorylation 48 h post-injection (Figure 5B). To understand if the effect of OLA in the skeletal muscle was secondary to the activation of the JNK1-driven hypothalamus-liver axis, insulin signaling was analyzed in skeletal muscle of mice with specific hypothalamic JNK1 deletion treated with OLA. As shown in Figure 5C, in mice with hypothalamic JNK1 deficiency under OLA i.p. treatment, AKT phosphorylation levels in skeletal muscle were higher compared to those of OLA-treated AAV-GFP control mice. A step further, the essential role of hepatic JNK activation for the observed skeletal muscle insulin resistance induced by OLA was assessed in L-JNK1/2-KO mice. In those mice OLA administered via intrahypothalamic injection did not impair AKT phosphorylation (Figure 5D). The response of skeletal muscle to AKT phosphorylation was also compared between L-JNK1/2-KO and Alb-Cre control mice following an intrahypothalamic injection of VEH or OLA. As shown in Figure S4B, a slight increase in insulin-induced AKT phosphorylation at Ser473 was observed in skeletal muscle ofL-JNK1/2-KO mice receiving VEH, whereas no differences were found at the Thr308 residue. Notably, comparisons between Alb-Cre and L-JNK1/2-KO mice following an OLA intrahypothalamic injection (same comparison as in Figure 5C), confirmed that L-JNK-1/2-KO mice exhibited higher insulin-induced AKT phosphorylation in skeletal muscle (Fig S4B), demonstrating the involvement of hepatic JNK1/2 in OLA-induced insulin resistance in this tissue. We also investigated whether disrupting the central-peripheral effects of OLA via the vagus nerve affects the responsiveness of skeletal muscle to insulin. As shown in Figure 5E, intrahypothalamic injection of OLA did not reduce AKT phosphorylation in skeletal muscle of vagotomized mice. Intriguingly, insulin-induced AKT phosphorylation in skeletal muscle was higher in vagotomized mice receiving OLA via intrahypothalamic injection than in those receiving VEH.

**Figure 5.**
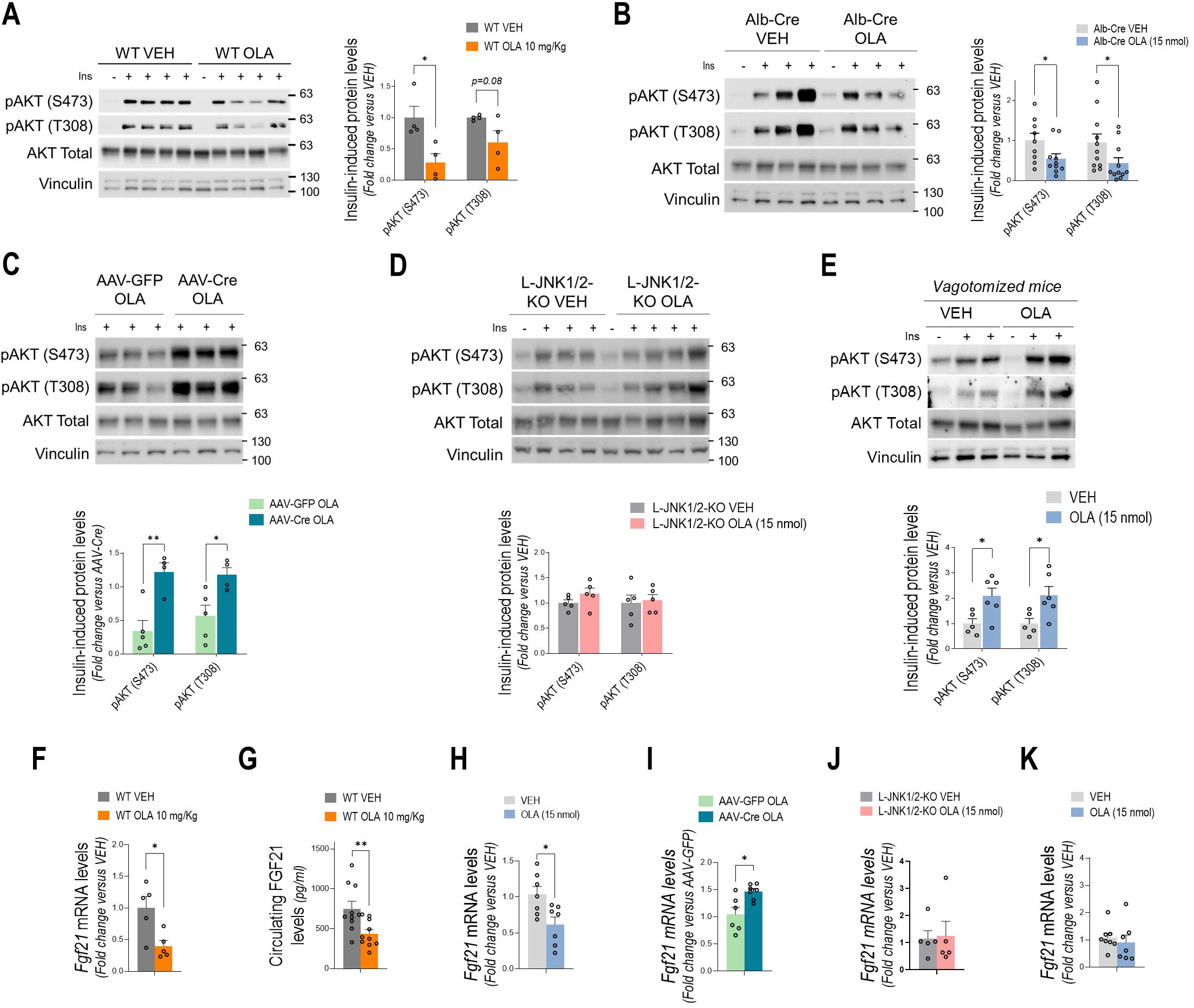
The hypothalamic JNK1-hepatic JNK1-FGF21 axis drive the impairment in skeletal muscle insulin signaling in male mice receiving an i.p. treatment with OLA. **A.** Representative Western blots of insulin-induced AKT phosphorylation (Ser473 and Thr308) in skeletal muscle of mice receiving OLA via i.p. and stimulated with insulin (0.75 U/kg) for the last 15 min, and densitometric quantification normalized for total AKT and vinculin levels (WT VEH: n=4; WT OLA: n=4). **B.** Representative Western blots of insulin-induced AKT phosphorylation (Ser473 and Thr308) in skeletal muscle of Alb-Cre mice 48 h after an OLA intrahypothalamic injection and stimulated with insulin (0.75 U/kg) for the last 15 min, and densitometric quantification normalized for total AKT and vinculin levels (Alb-Cre VEH: n=9; Alb-Cre OLA: n=10). **C.** Representative Western blots of insulin-induced AKT phosphorylation (Ser473 and Thr308) in skeletal muscle of mice with specific JNK1 deletion in the hypothalamus receiving OLA via i.p. and stimulated with insulin (0.75 U/kg) for the last 15 min, and densitometric quantification normalized for total AKT and vinculin levels (AAV-GFP OLA: n=5; AAV-Cre OLA: n=4). **D.** Representative Western blots of insulin-induced AKT phosphorylation (Ser473 and Thr308) in skeletal muscle of L-JNK1/2-KO mice 48 h after an OLA intrahypothalamic injection and stimulated with insulin (0.75 U/kg) for the last 15 min, and densitometric quantification normalized for total AKT and vinculin levels (L-JNK1/2-KO DMSO: n=5; L-JNK1/2-KO OLA: n=4). **E.** Representative Western blots of insulin-induced AKT phosphorylation (Ser473 and Thr308) in skeletal muscle of vagotomized mice 48 h after an OLA intrahypothalamic injection and stimulated with insulin (0.75 U/kg) for the last 15 min, and densitometric quantification normalized for total AKT and vinculin levels (DMSO: n=5; OLA: n=5). F. *Fgf21* mRNA levels using *Actb* as housekeeping gene in the liver of mice receiving OLA via i.p. for 8 weeks (WT VEH: n= 5; WT OLA: n= 5). **G.** FGF21 circulating levels in mice receiving OLA via i.p. for 8 weeks (WT VEH: n= 10; WT OLA: n= 10). H. *Fgf21* mRNA levels using *Actb* as housekeeping gene in the liver of mice 8 h after an OLA intrahypothalamic injection (DMSO: n= 7; OLA: n= 7). I. *Fgf21* mRNA levels using *Actb* as housekeeping gene in the liver of mice with JNK1 specifically deleted from the hypothalamus and receiving OLA via i.p. for 8 weeks (AAV-GFP VEH: n= 6; AAV-Cre OLA: n= 7). J. *Fgf21* mRNA levels using *Actb* as housekeeping gene in the liver of L-JNK1/2-KO mice 8 h after an OLA intrahypothalamic injection (L-JNK1/2-KO DMSO: n= 5; L-JNK1/2-KO OLA: n=5). K. *Fgf21* mRNA levels using *Actb* as housekeeping gene in the liver of vagotomized mice 8 h after an OLA intrahypothalamic injection (DMSO: n= 8; OLA: n= 7). Each point/bar corresponds to mean ± SEM; comparisons between groups: p < 0.05; p < 0.01; p < 0.001.

Since Vernia *et al.* reported that activation of hepatic JNK represses *Fgf21* expression via peroxisome proliferator-activated receptor α (PPARα) [59], we measured *Fgf21* mRNA levels in the liver and found a significant reduction in male mice receiving OLA via i.p (Figure 5F).

We measured circulating FGF21 levels, which were also reduced in OLA-treated mice (Figure 5G). The ability of OLA to downregulate hepatic *Fgf21* mRNA was also evident in mice 8 h after receiving a single intrahypothalamic injection (Figure 5H). Importantly, mice lacking either hypothalamic (Figure 5I) or hepatic (Figure 5J) JNK1, as well as vagotomized mice (Figure 5K), were protected from the OLA-induced decrease in hepatic *Fgf21*. Collectively, our results suggest that OLA activates JNK1 in the hypothalamus, which transmits signals through the vagus nerve that induce JNK activation in the liver, thereby reducing *Fgf21* expression and circulating FGF21 levels, which may account for OLA-induced insulin resistance in skeletal muscle. To our knowledge, this is the first study linking central JNK1 to a liver-muscle axis via FGF21 in the context of antipsychotic-induced metabolic dysfunction.

### 3.6. Fgf21 overexpression in the liver prevents insulin resistance in skeletal muscle induced by the OLA i.p. treatment

To directly demonstrate the relevance of FGF21 in the effect of OLA on insulin sensitivity, FGF21 was overexpressed in the liver of mice by adenoviral delivery prior to i.p. OLA treatment. Hepatic *Fgf21* overexpression was assessed by measuring FGF21 levels in circulation (Figure 6A). Evaluation of body weight through the treatment revealed that mice with hepatic *Fgf21* overexpression had similar weight loss as their control mice without FGF21 overexpression after OLA treatment (Figure 6B and S5A). As previously reported [60], under basal conditions the body weight of mice overexpressing *Fgf21* in the liver was lower than control mice. Moreover, hepatic *Fgf21* overexpression protected against whole body insulin resistance induced by OLA, but not against pyruvate intolerance, reinforcing that skeletal muscle insulin sensitization relays on liver-derived FGF21 (Figures 6C-E). Importantly, these mice showed increased JNK phosphorylation in both the hypothalamus and liver (Figures 6F-G) after i.p. OLA treatment, similar to their controls without hepatic FGF21 overexpression (Figures S5B-C). At the molecular level, mice overexpressing hepatic *Fgf21* exhibited impairments in hepatic (Figure 6H) but not skeletal muscle (Figure 6I) insulin signaling, consistent with the lack of decrease in liver *Fgf21* expression upon OLA treatment (Figure 6J). As expected, AAV-CTRL mice exhibited impaired insulin signaling in both liver and skeletal muscle (Figures S5D-E). Collectively, these results support the reduction of hepatic FGF21 as a driver of OLA-induced insulin resistance in skeletal muscle as a consequence of the hypothalamus-liver signals described above.

**Figure 6.**
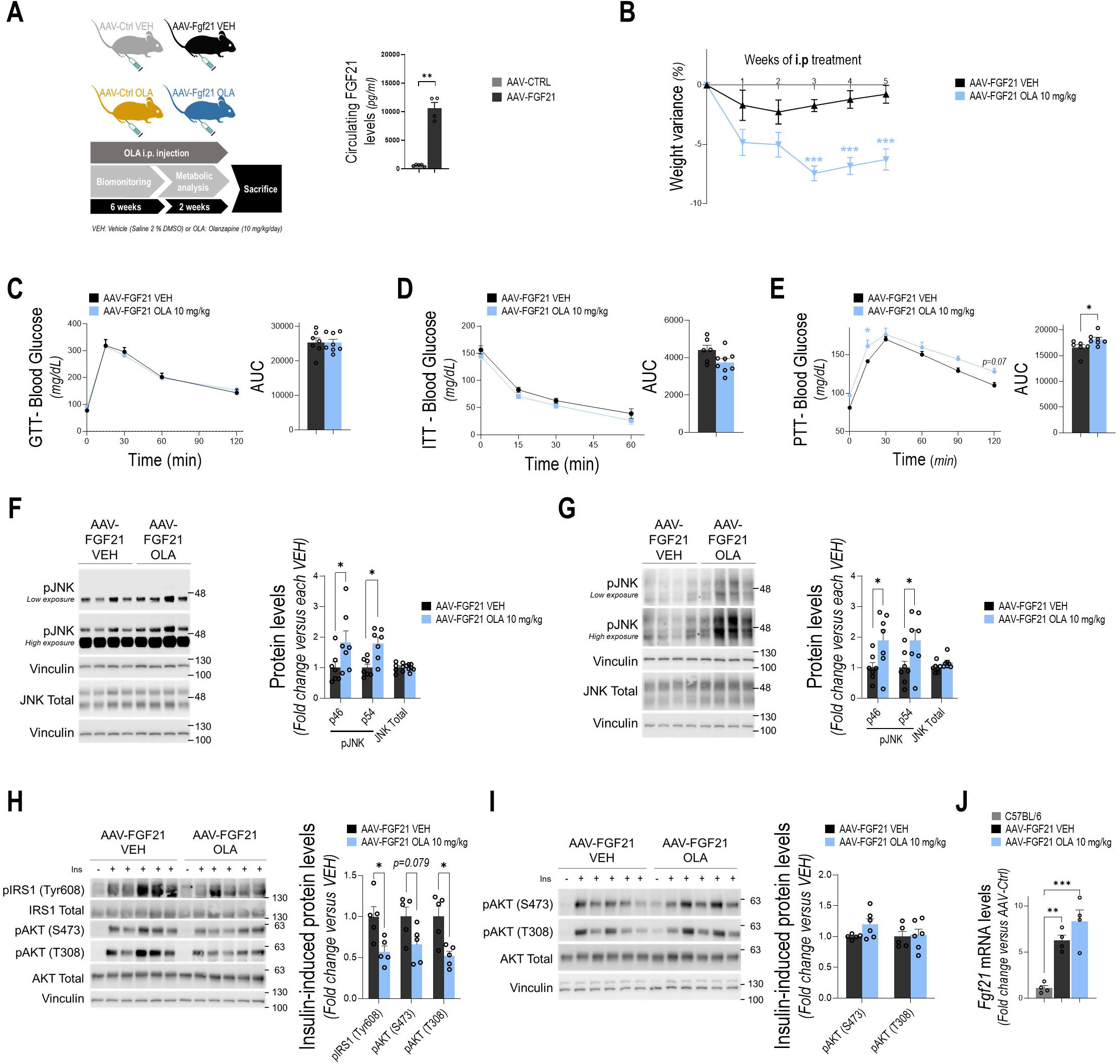
Hepatic FGF21 overexpression prevents insulin resistance in skeletal muscle, but not in the liver, in male mice receiving OLA via i.p. **A.** Experimental design of *Fgf21* overexpression in the liver prior to OLA i.p. daily injections for 8 weeks. FGF21 circulating levels showing that the virogenetic approach result in a significant increase of FGF21 circulating levels (AAV-CTRL: n= 5; AAV-FGF21: n= 4). **B.** Body weight change of mice with hepatic FGF21 overexpression receiving OLA via i.p. (AAV-FGF21 VEH: n= 7; AAV-FGF21 OLA: n= 8). **C.** GTT (AAV-FGF21 VEH: n= 7; AAV-FGF21 OLA: n= 8), **D.** ITT (AAV-FGF21 VEH: n= 6; AAV-FGF21 OLA: n= 8) and **E.** PTT (AAV-FGF21 VEH: n= 6; AAV-FGF21 OLA: n= 7) at the end of the treatment and their respective AUC for each group. **F.** Representative Western blots of JNK phosphorylation in the hypothalamus of AAV-FGF21 injected mice treated with OLA via i.p. and densitometric quantification normalized for total JNK and vinculin levels (AAV-FGF21 VEH: n= 7; AAV-FGF21 OLA: n= 7). **G.** Representative Western blots of JNK phosphorylation in liver of AAV-FGF21 injected mice treated with OLA via i.p. and densitometric quantification normalized for total JNK and vinculin levels (AAV-FGF21 VEH: n= 7; AAV-FGF21 OLA: n= 7). **H.** Representative Western blots of insulin-induced IRS1 phosphorylation in Tyr608 residues and AKT phosphorylation (Ser473 and Thr308) in liver of AAV-FGF21 mice receiving OLA via i.p. and stimulated with insulin (0.75 U/kg) for the last 15 min, and densitometric quantification normalized for total IRS1, AKT and vinculin levels (AAV-FGF21 VEH: n= 5; AAV-FGF21 OLA: n= 5). **I.** Representative Western blots of insulin-induced AKT phosphorylation (S473 and T308) in skeletal muscle of AAV-FGF21 injected mice receiving OLA via i.p. and stimulated with insulin (0.75 U/kg) for the last 15 min, and densitometric quantification normalized for total IRS1, AKT and vinculin levels (AAV-FGF21 VEH: n= 5; AAV-FGF21 OLA: n= 6). J. *Fgf21* mRNA levels using *Actb* as housekeeping gene in the liver of AAV-FGF21 injected mice receiving OLA via i.p. for 8 weeks (C57BL/6: n= 4; AAV-FGF21 VEH: n= 4; AAV-FGF21 OLA: n= 4, groups were compared using One Way-ANOVA with Bonferroni’s post-hoc test carried for multiple comparisons between the groups). Each point/bar corresponds to mean ± SEM; comparisons between groups: p < 0.05; p < 0.01; p < 0.001.

### 3.7. PTP1B-deficient mice are protected against insulin resistance induced by OLA i.p. treatment

To confirm the inter-organ crosstalk as responsible for OLA-induced insulin resistance driven by hypothalamic JNK1, we conducted experiments in PTP1B-deficient mice treated with OLA via i.p. during 8 weeks since, as we reported, they are protected against OLA-induced JNK activation in this brain region [39]. As expected, male mice lacking PTP1B were protected against OLA-induced disturbances in glucose homeostasis (Figure 7A-C). In this line, hepatic insulin signaling was preserved in PTP1B KO mice receiving OLA, an effect also confirmed in primary hepatocytes isolated at the end of treatment (Figure 7D and Figure S6A). In addition, basal glucose uptake was higher in hepatocytes from OLA-treated PTP1B KO mice compared to hepatocytes from vehicle-treated mice (Figure S6B). Consistent with our hypothesis, OLA-treated PTP1B KO mice did not show activation of hepatic JNK (Figure 7E), preserved IRS1 tyrosine phosphorylation (Figure 7F), and were protected against the decrease in FGF21 levels both in the liver (Figure 7G) and in the circulation (Figure 7H). In this line, PTP1B KO mice treated with OLA responded to insulin by inducing AKT (S473 and T308) phosphorylation in skeletal muscle to a similar extent as control PTP1B KO mice (Figure 7I).

**Figure 7.**
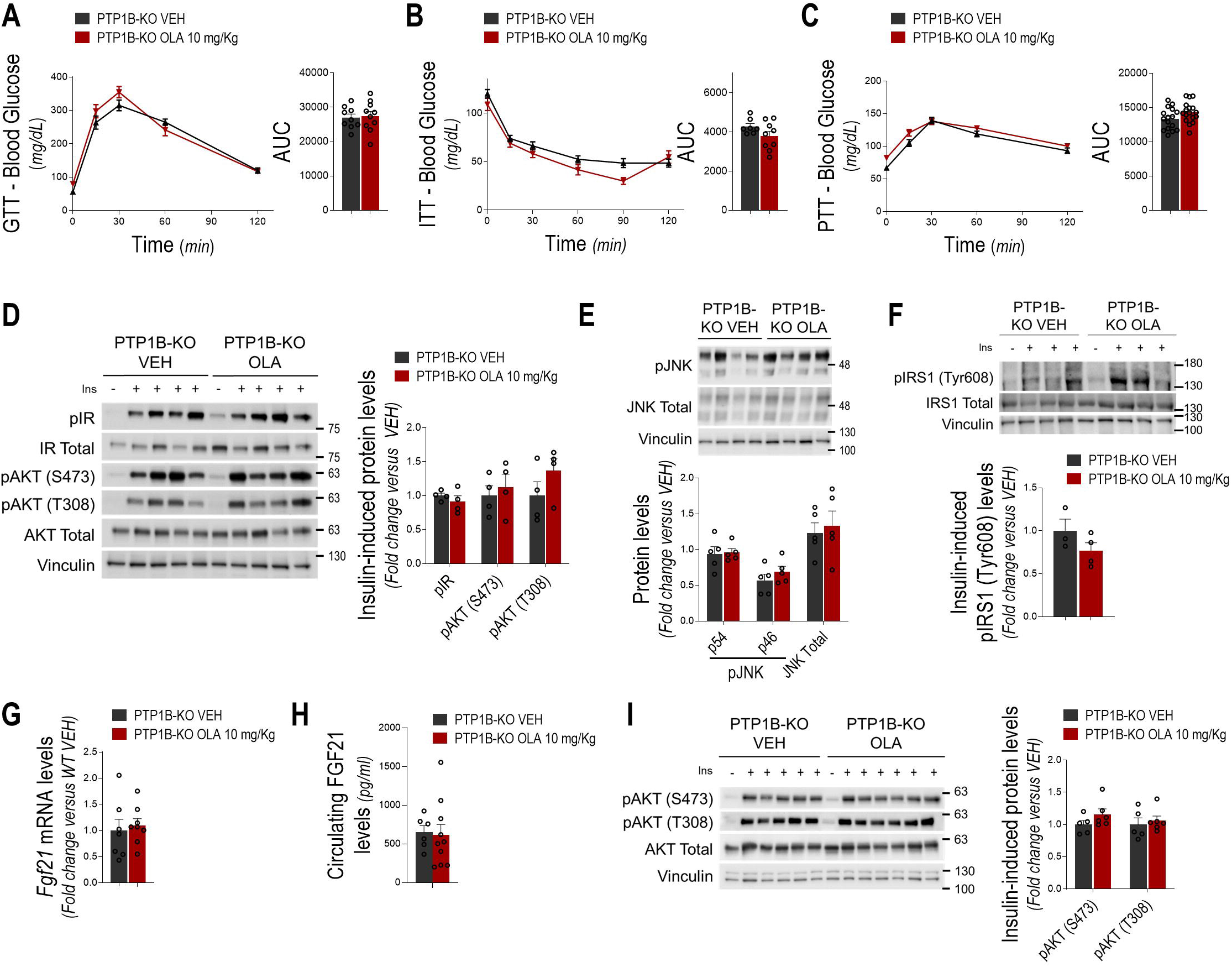
PTP1B deficiency protects male mice against induced insulin resistance induced by an i.p. treatment with OLA. PTP1B-KO mice received OLA via i.p. daily for 8 weeks. **A.** GTT (PTP1B-KO VEH: n= 9; PTP1B-KO OLA: n= 10), **B.** ITT (PTP1B-KO VEH: n= 7; PTP1B-KO OLA: n= 9) and **C.** PTT (PTP1B-KO VEH: n= 16; PTP1B-KO OLA: n= 17) at the end of the treatment and respective AUC for each group. **D.** Representative Western blots of IR and AKT phosphorylation (Ser473 and Thr308) in liver of PTP1B-KO mice receiving OLA via i.p. and stimulated with insulin (0.75 U/kg) for the last 15 min, and densitometric quantification normalized for total IR, AKT and vinculin levels (PTP1B-KO VEH: n= 4; PTP1B-KO OLA: n= 4). **E.** Representative Western blots of hepatic JNK in PTP1B-KO mice treated with OLA via i.p. and densitometric quantification normalized for total JNK and vinculin levels (PTP1B-KO VEH: n= 5; PTP1B-KO OLA: n= 5). **F.** Representative Western blots of insulin-induced IRS1 phosphorylation in Tyr608 residues in liver of PTP1B-KO mice receiving OLA via i.p. and stimulated with insulin (0.75 U/kg) for the last 15 min, and densitometric quantification normalized for total IRS1 and vinculin levels (PTP1B-KO VEH: n= 3; PTP1B-KO OLA: n= 4). **G.** *Fgf21* mRNA levels using *Actb* as housekeeping gene in the liver of PTP1B-KO mice receiving OLA via i.p. for 8 weeks (PTP1B-KO VEH: n= 7; PTP1B-KO OLA: n= 8). **H.** FGF21 circulating levels of PTP1B-KO mice receiving OLA via i.p. for 8 weeks (PTP1B-KO VEH: n= 6; PTP1B-KO OLA: n= 10). **I.** Representative Western blots of AKT phosphorylation (Ser473 and Thr308) in liver of PTP1B-KO mice receiving OLA via i.p. and stimulated with insulin (0.75 U/kg) for the last 15 min, and densitometric quantification normalized for total AKT and vinculin levels (PTP1B-KO VEH: n= 5; PTP1B-KO OLA: n= 6). Each point/bar corresponds to mean ± SEM.

## 4. Discussion

Although accumulating evidence has linked SGA therapy to metabolic disturbances [13, 14], at the molecular level there is still a gap regarding the knowledge of tissue-specific effects of these drugs leading to insulin resistance and the identification of potential therapeutic targets to ameliorate this deleterious side-effect. Our results herein have uncovered a mechanistic link between central signals originating from the hypothalamus and relevant peripheral insulin target tissues in male mice receiving OLA via i.p. In particular, we identified: i) a hypothalamus-liver-skeletal muscle axis controlled by hypothalamic JNK1 which activates hepatic JNK1 via the vagus nerve; ii) JNK1 activation in the liver leading, on the one hand, to hepatic insulin resistance by increasing IRS1 serine phosphorylation and, on the other, to downregulation of hepatic *Fgf21* which, in turn, impairs insulin signaling in skeletal muscle; iii) the inhibition of PTP1B as an effective strategy to ameliorate OLA-induced insulin resistance by targeting this central-peripheral axis.

Our recent study has demonstrated the benefit of administering OLA via i.p. to male mice, showing that it reduces body weight by increasing energy expenditure (EE) without altering food intake, through modulation of the hypothalamic AMPK-BAT UCP1 axis [16]. This highlights the therapeutic advantage of this administration route compared to oral formulations, which are associated with obesity [13, 14, 16]. However, it should be noted that other studies administering OLA orally [61] or via intramuscular injections [62] found increases in BAT UCP1 and EE without reductions in food intake or body weight. This strongly suggests that, under circumstances that are not yet fully understood, BAT thermogenesis might not be sufficient to counteract the hyperphagia associated with OLA oral treatment. In a subsequent study, we identified the liver as another peripheral tissue targeted by hypothalamic signals induced by OLA through rewiring of lipid metabolism [39], implicating the hypothalamus as a major controller of the peripheral metabolic responses during OLA i.p. treatment in male mice. Interestingly, low-grade inflammatory features were found in the hypothalamus of male mice receiving an i.p. treatment with OLA that were attenuated by deleting JNK1 in this brain region. Since neuroinflammation is closely linked with insulin resistance and T2D [40], in the present study we have extensively evaluated the peripheral insulin responses to the OLA i.p. treatment, particularly in the liver and skeletal muscle.

Despite protection against body weight gain and hepatic steatosis, our results demonstrated that male mice receiving OLA treatment via i.p. developed systemic insulin resistance and pyruvate intolerance. Consistent with this metabolic phenotype, reduced insulin responsiveness was observed in the critical nodes of the insulin signaling cascade in the liver and skeletal muscle. The importance of central signals that control hepatic insulin sensitivity is supported by the lack of effect of OLA in impairing insulin signaling in hepatocytes treated with this SGA in vitro, as well as in the livers of vagotomized mice. With regard to the latter, although vagotomy was sufficient to blunt OLA-induced hepatic insulin resistance, it cannot be ruled out that other nerves such, as the sympathetic nerves, or hormonal systems, may also be involved in transmitting of signals from the hypothalamus to the periphery. Indeed, insulin-induced AKT phosphorylation was higher in the liver and skeletal muscle of vagotomized mice receiving OLA, than in VEH-treated mice (Figures 4E and 5E, respectively). Of note, this higher effect was also observed in the liver of mice with JNK deletion in the hypothalamus compared to VEH-treated mice (Figure S3A). Therefore, it is tempting to speculate that OLA might target additional signaling pathways that become evident when the hypothalamic JNK-driven hepatic JNK axis is disrupted. The fact that OLA does not affect insulin sensitivity in hepatocytes *in vitro* suggests that the enhancement of insulin signaling observed *in vivo* under these conditions is not the result of a direct effect of this SGA in the liver. Furthermore, the absence of significant inflammatory and/or oxidative stress markers in the liver eliminates the intrahepatic microenvironment as a trigger of insulin resistance in this organ. These results are also supported by the lack of OLA’s direct effect on activating JNK in primary hepatocytes, a phenomenon observed in vivo when male mice were treated with OLA via i.p. Notably, the lack of effect of OLA in altering hepatocytes’ insulin response is consistent with previous reports by Levkovitz *et al.* in FAO hepatoma cells [63].

JNK plays a central role in oxidative stress and inflammatory responses [64, 65] both of which were previously found to be elevated in the hypothalamus of male mice receiving OLA via i.p. treatment concomitant with JNK activation. At the cellular level, OLA increases JNK phosphorylation in neurons, astrocytes and microglia in culture [39]. Therefore, the combined response of these cell populations in the hypothalamus of OLA-treated mice may explain the central signals that connect to the liver. Our results herein show a significant increase in JNK phosphorylation in the livers of mice treated with OLA via i.p., likely due to hypothalamic activation of this kinase. This is supported by the absence of hepatic JNK activation in mice with specific deletion of JNK1 in the hypothalamus. On the other hand, JNK is a main hub in obesity and insulin resistance in the central nervous system (CNS) and the periphery [66–68]. Indeed, JNK activity is elevated in muscle, fat, and liver of obese mice, and obese mice with systemic ablation of JNK1, but not JNK2, exhibit a dramatically improved metabolic phenotype [66]. Regarding the CNS, mice with specific deletion of JNK1 in nestin-expressing neurons are protected against HFD-induced insulin resistance concurrently with improvements in glucose homeostasis, hepatic insulin signaling and reduction of hepatosteatosis [69].

JNK has been proposed to drive insulin resistance in obesity by distinct mechanisms, including the direct inhibitory phosphorylation of IRS proteins, the promotion of adiposity and metabolic inflammation, and the negative regulation of the PPARα-FGF21 axis [66, 70]. As reviewed [71], JNK has been identified in many studies as an IRS1 kinase that targets serine and threonine residues. Indeed, JNK1-deficient mice are protected against insulin resistance, which correlates with enhanced insulin-stimulated IRS1 tyrosine phosphorylation and reduced IRS1 serine phosphorylation in the liver [66]. In the context of OLA effects in the liver of male mice receiving an i.p. treatment, a marked increase in phospho-JNK was observed alongside an increase in IRS1 serine phosphorylation and a decrease in insulin-induced IRS1 tyrosine phosphorylation. This molecular signature of hepatic insulin resistance was also observed when OLA was administered via intrahypothalamic injection, supporting the pivotal role of central-peripheral connections in OLA-induced impairments of insulin sensitivity and glucose homeostasis. Of note, Li *et al.* reported that the i.p. treatment with OLA increased JNK phosphorylation in adipose tissue of female mice and in a cell-autonomous manner in 3T3L1 adipocytes, the latter being associated with increased IRS1 serine phosphorylation [72]. The authors of this study claimed that OLA-mediated increases in JNK/IRS1 serine phosphorylation induce insulin resistance in adipocytes; however, experiments with insulin stimulation in cells treated with OLA were not provided. The relevance of hepatic JNK1 activation in OLA-induced insulin resistance was supported by protection against a decrease in insulin signaling in mice with specific JNK1 deletion in hepatocytes. Consistent with the absence of hepatic JNK phosphorylation, deletion of hypothalamic JNK1 also prevented OLA-induced impairment of insulin signaling in the liver. This strongly supports the idea that modulation of the JNK1-driven hypothalamic-liver axis is pivotal for OLA-induced impairment of hepatic insulin signaling.

FGF21 is mainly secreted by the hepatocytes and elicits a variety of metabolic benefits in obese mouse models by enhancing lipid metabolism, reducing blood lipids, ameliorating insulin resistance, and improving metabolic syndrome [73]. FGF21 is a potent inducer of BAT thermogenesis and iWAT browning and also improves weight loss [74]. At the molecular level, the study of Vernia *et al.* [59] demonstrated that hepatic JNK represses the nuclear hormone receptor PPARα, thereby decreasing the expression of its target genes including FGF21. Silencing *Fgf21* in the liver prevented the beneficial effects of hepatic JNK inhibition, including increased ketogenesis and improved systemic insulin sensitivity [59]. These results highlight JNK signaling as a critical mediator of this hepatokine. Consistent with this previous study, our results herein provide evidence of protection against the decline in skeletal muscle insulin signaling and peripheral insulin resistance in mice overexpressing FGF21 in the liver and receiving i.p. OLA treatment. Notably, in these mice, the impairment of hepatic insulin signaling by OLA was not counteracted by FGF21 overexpression, implicating FGF21 as a downstream JNK target in the liver during OLA treatment via i.p. A step further, our study links hypothalamic JNK activation with FGF21 secretion by the liver and its insulin-sensitizing effect in skeletal muscle in the context of OLA-induced metabolic disturbances. Further research is needed to reveal the specific FGF21-mediated signaling pathways in skeletal muscle during OLA i.p. treatment.

In addition to providing mechanistic insights into the central and peripheral effects of the treatment with OLA via i.p. on insulin sensitivity and glucose homeostasis, the present study demonstrates the beneficial effects of PTP1B inhibition in protecting against liver and skeletal muscle insulin resistance induced by i.p. OLA treatment in male mice. Both tissues are extensively studied targets of PTP1B actions in insulin signaling [37, 75–77]. These findings are not surprising since protection against insulin resistance by PTP1B inhibition in different physiopathological conditions, including obesity and aging, has been reported by our group [34–36] and others [38, 76]. Nevertheless, since PTP1B-deficient mice did not show hypothalamic inflammation/oxidative stress and thus JNK activation in this brain region, the experiments performed in these mice in the present study model confirm the mechanistic insights into JNK1-driven central-peripheral crosstalk that leads to impaired insulin sensitivity in male mice receiving OLA via i.p. This was particularly evident in the protection of PTP1B-deficient mice against intrahepatic (e.g., JNK activation, impaired insulin signaling, and *Fgf21* downregulation) and extrahepatic (e.g., decrease in serum FGF21 and impaired insulin signaling in skeletal muscle) OLA effects. However, mice with specific deficiency of PTP1B in central and peripheral compartments would be useful in further studies to determine the tissue-specific relevance of this phosphatase in the metabolic side-effects of OLA i.p. treatment. Nevertheless, PTP1B inhibition is a current therapeutic strategy for improving insulin sensitivity as demonstrated by several groups including ours [35, 37, 38, 75, 77]. Therefore, we propose combining OLA treatment with a PTP1B inhibitor could address this metabolic comorbidity in patients under OLA treatment. Of note, clinical trials ofPTP1B inhibitors such as trodusquemine (i.e., NCT00606112) are underway and, surprisingly, trodusquemine alleviated schizophrenia-like symptoms in mice [78]. Taken together, these findings and our results suggest that targeting PTP1B could be a therapeutic strategy to prevent insulin resistance in patients receiving OLA either orally or by long-term injectable treatments.

It is also noteworthy to highlight that a previous study by Kowalchuk *et al.* [79] showed that OLA impaired the ability of a central insulin infusion to suppress hepatic glucose production raising the possibility that the OLA-induced insulin resistance may also result from additional central alterations. In a subsequent study they found that attenuation of the activation of central ATP-sensitive potassium (K_ATP_) channels, key metabolic sensors downstream of hypothalamic insulin signaling that are closely linked to the maintenance of systemic glucose homeostasis [20], is likely the trigger for OLA-induced insulin resistance. These results suggest that additional mechanisms linking central and peripheral tissues may be relevant to OLA-induced disturbances of glucose homeostasis and should be considered.

Although it is highly relevant to uncover the molecular mechanisms by which OLA induces insulin resistance independently of body weight gain, our study has some limitations. In particular, future studies should investigate alternative mechanisms and molecular mediators in experimental models in which OLA promotes weight gain without activating the hypothalamic JNK-hepatic JNK axis [39]. Another limitation concerns the translation of our preclinical findings to humans. In clinical studies where OLA was administered at high doses (up to 50 mg/day), the amount of drug reaching the VMH was not determined. Furthermore, differences in the half-life of OLA across species (approximately 3 h in mice [80], 2.5-3 h in rats [81] and 21-54 h in humans [82]) further complicate the extrapolation of preclinical data to a clinical setting.

## 5. Conclusion

Despite of the beneficial effects of OLA administered via i.p. in preventing weight gain and hepatic steatosis, it induces peripheral insulin resistance through a hypothalamus-liver axis controlled by hypothalamic JNK1 activation, which, in turn, activates hepatic JNK via the vagus nerve. This activation suppresses FGF21 production, thereby promoting insulin resistance in the liver and skeletal muscle, respectively. Our results have uncovered OLA-mediated hypothalamic signaling as a driver of skeletal muscle insulin resistance secondary to hepatic dysfunction, as this inter-organ crosstalk is prevented when central signals are abolished. In this context, PTP1B deficiency protects against OLA-induced insulin resistance by targeting hypothalamic JNK1, supporting the beneficial effects of inhibiting PTP1B in ameliorating the metabolic side-effects of this SGA and offering therapeutic benefits.

## Abbreviations

SGA: Second generation antipsychotics
OLA: olanzapine
T2D: Type 2 Diabetes
GTT: glucose tolerance test
BAT: brown adipose tissue
UCP-1: uncoupling protein 1
GLUT4: Glucose transporter 4
WAT: White adipose tissue
NF_κ_B: nuclear factor kappa B
IRS1: insulin receptor substrate 1
GSK-3: glycogen synthase kinase 3
PTP1B: Protein tyrosine phosphatase 1B
AMPK: AMP-activated protein kinase
JNK: c-jun n-terminal kinase
FAS: fatty acid synthase
EE: energy expenditure
PTP1B-KO: PTP1B-Knockout
WT: Wild-type
L-JNK1/2-KO: liver-specific JNK1/2-KO
i.p.: intraperitoneal
AAV: adeno-associated viruses
VMH: ventromedial nucleus of the hypothalamus
AUC: area under the curve
ITT: insulin tolerance test
PTT: pyruvate tolerance test
KRP: Krebs Ringer-phosphate buffer

## CRediT authorship contribution statement

The study was designed by VF, PR and ÁMV. Mice treatments, data acquisition, analysis and interpretation were performed by VF, PR and ÁMV. Immunohistochemistry and immunofluorescence analysis were performed by VF. Protein expression analysis was performed by VF, ABH, AMSL, PR and ÁMV. Primary hepatocytes were isolated by PR and VF. Central injections were performed by VF, CF, ÁE and GS. Central adenoviral injections and posterior treatment with OLA were done by CF, VF, GS and RJD. VF, PR and ÁMV wrote the first draft of the manuscript. CF, ABH, AMSL, ML, GS and RJD critically revised the manuscript for important intellectual content. All authors gave final approval of the manuscript and gave consent to its publication. ÁMV and PR coordinated the study and ÁMV is the guarantor of the work.

## Acknowledgements

The authors would like to thank all members of AMV’s laboratory for helpful discussions. We also acknowledge M. Martín Belinchón (IIBM, CSIC) for the technical assistance with the confocal microscopy.

## Data availability

Data presented in this manuscript are available upon request from the corresponding author.

---

***Supplementary figure 1. Plasma glucose and insulin levels of male mice receiving an OLA i.p. treatment for 8 weeks.* A.** Plasma glucose levels in fast (16-18 h) and fed state of mice receiving OLA via i.p. daily for 8 weeks (WT VEH: n= 5-6; WT OLA: n=5-6). **B.** Plasma insulin levels in fast (16-18 h) and fed state of mice receiving OLA via i.p. daily for 8 weeks (WT VEH: n=9; WT OLA: n=8). Each point/bar corresponds to mean ± SEM; comparisons between groups: p < 0.05; p < 0.01; p < 0.001.

***Supplementary Figure 2. OLA treatment does not affect JNK phosphorylation in primary mouse hepatocytes.* A.** Representative Western blots of JNK phosphorylation in primary hepatocytes treated *in vitro* with OLA (12.5 µM) for several time periods and densitometric quantification normalized for total JNK and vinculin levels (DMSO: n=4; OLA: n=4). Each point/bar corresponds to mean ± SEM.

***Supplementary Figure 3. Hypothalamic JNK1 deletion prevents OLA-induced attenuation in hepatic insulin signaling in male mice.* A.** Representative Western blots of insulin-induced AKT phosphorylation (Ser473 and Thr308) in liver of AAV-GFP control mice (left panel) or mice with hypothalamic JNK1 deletion (AAV-Cre, right panel) receiving VEH or OLA via i.p. Densitometric quantification normalized for total AKT and vinculin levels (AAV-GFP VEH: n=6; AAV-GFP OLA: n=4; AAV-Cre VEH: n=4; AAV-Cre OLA: n=4). **B.** Stomachs from vagotomized and sham-operated mice showing an evident increase in size after vagotomy due to motoric dysfunction. Each point/bar corresponds to mean ± SEM; comparisons between groups: p < 0.05; p < 0.01; p < 0.001.

***Supplementary Figure 4. Hepatic JNK1 deletion prevents OLA-induced attenuation in liver and skeletal muscle insulin signaling in male mice.* A.** Representative Western blots of insulin-induced AKT phosphorylation (Ser473 and Thr308) in liver of mice with liver JNK1 deletion (L-JNK1/2-KO) or Alb-Cre control mice receiving VEH (left panel) or OLA (right panel) via intrahypothalamic. Densitometric quantification normalized for total AKT and vinculin levels (Alb-Cre VEH: n=6; L-JNK1/2-KO VEH: n=4; Alb-Cre OLA: n=5; L-JNK1/2-KO OLA: n=5). **B.** Representative Western blots of insulin-induced AKT phosphorylation (Ser473 and Thr308) in skeletal muscle of mice with liver JNK1 deletion (L-JNK1/2-KO) or Alb-Cre control mice receiving VEH (left panel) or OLA (right panel) via intrahypothalamic. Densitometric quantification normalized for total AKT and vinculin levels (Alb-Cre VEH: n=6; L-JNK1/2-KO VEH: n=5; Alb-Cre OLA: n=5; L-JNK1/2-KO OLA: n=5). Each point/bar corresponds to mean ± SEM; comparisons between groups: p < 0.05.

***Supplementary Figure 5. The effect of an i.p. treatment with OLA in systemic and hepatic insulin sensitivity in AAV-CTRL injected male mice is similar to that of WT mice.* A.** Body weight change of mice injected AAV-CTRL and receiving OLA via i.p. (AAV-CTRL VEH: n= 7; AAV-CTRL OLA: n= 8). **B.** Representative Western blots of JNK phosphorylation in the hypothalamus of AAV-CTRL injected mice treated with OLA via i.p. and densitometric quantification normalized for total JNK and vinculin levels (AAV-CTRL VEH: n= 6-7; AAV-CTRL OLA: n= 7-8). **C.** Representative Western blots of JNK phosphorylation in liver of AAV-CTRL injected mice treated with OLA via i.p. and densitometric quantification normalized for total JNK and vinculin levels (AAV-CTRL VEH: n= 6-7; AAV-CTRL OLA: n= 7-8). **D.** Representative Western blots of insulin-induced IRS1 phosphorylation in Tyr608 residues and AKT phosphorylation (Ser473 and Thr308) in liver of AAV-CTRL mice receiving OLA via i.p. and stimulated with insulin (0.75 U/kg) for the last 15 min, and densitometric quantification normalized for total IRS1, AKT and vinculin levels (AAV-CTRL VEH: n= 4-5; AAV-CTRL OLA: n= 7). **E.** Representative Western blots of insulin-induced AKT phosphorylation (Ser473 and Thr308) in skeletal muscle of AAV-CTRL injected mice receiving OLA via i.p. and stimulated with insulin (0.75 U/kg) for the last 15 min, and densitometric quantification normalized for total IRS1, AKT and vinculin levels (AAV-CTRL VEH: n= 4-5; AAV-CTRL OLA: n= 7). Each point/bar corresponds to mean ± SEM; comparisons between groups: p < 0.05; p < 0.01; p < 0.001.

***Supplementary Figure 6. PTP1B-deficient hepatocytes are protected against insulin resistance induced by the treatment with OLA via i.p.* A.** Representative Western blots of insulin-induced AKT (Ser473 and Thr308) phosphorylation in primary hepatocytes isolated from VEH- or OLA-treated PTP1B-KO mice via i.p. and stimulated with insulin (10 nM, 15 min). Densitometric quantification normalized for total AKT levels (PTP1B-KO VEH: n=3; PTP1B-KO OLA: n=2). **F.** Glucose uptake (fold of change versus VEH) of primary hepatocytes from OLA-treated male mice (PTP1B-KO VEH: n=6; PTP1B-KO OLA: n=6). Each point/bar corresponds to mean ± SEM; comparisons between groups: p < 0.05; p < 0.01; p < 0.001.

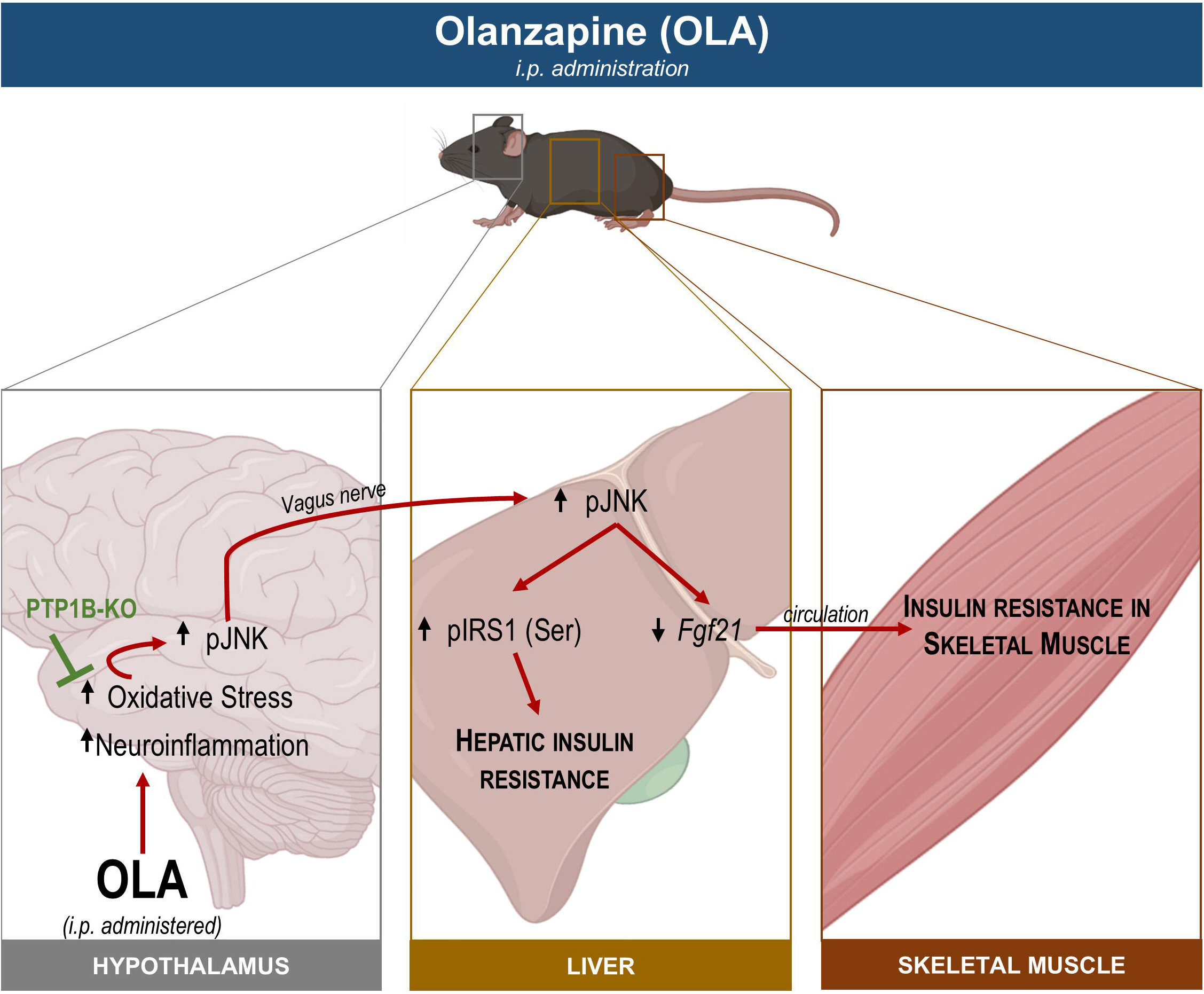

## References

[1] K.R. Patel, J. Cherian, K. Gohil, D. Atkinson, Schizophrenia: overview and treatment options, P T 39(9) (2014) 638–45.

[2] G. Remington, D. Addington, W. Honer, Z. Ismail, T. Raedler, M. Teehan, Guidelines for the Pharmacotherapy of Schizophrenia in Adults, Can J Psychiatry 62(9) (2017) 604–616.

[3] M. Fonseca, F. Carmo, F. Martel, Metabolic effects of atypical antipsychotics: Molecular targets, J Neuroendocrinol 35(12) (2023) e13347.

[4] D. Cohen, R.P. Stolk, D.E. Grobbee, C.C. Gispen-de Wied, Hyperglycemia and diabetes in patients with schizophrenia or schizoaffective disorders, Diabetes Care 29(4) (2006) 786–91.

[5] D.E. Casey, Dyslipidemia and atypical antipsychotic drugs, J Clin Psychiatry 65 Suppl 18 (2004) 27–35.

[6] I. Kurzthaler, W.W. Fleischhacker, The clinical implications of weight gain in schizophrenia, J Clin Psychiatry 62 Suppl 7 (2001) 32–7.

[7] S. Leucht, A. Cipriani, L. Spineli, D. Mavridis, D. Orey, F. Richter, M. Samara, C. Barbui, R.R. Engel, J.R. Geddes, W. Kissling, M.P. Stapf, B. Lassig, G. Salanti, J.M. Davis, Comparative efficacy and tolerability of 15 antipsychotic drugs in schizophrenia: a multiple-treatments meta-analysis, Lancet 382(9896) (2013) 951–62.

[8] A.P. Rajkumar, H.T. Horsdal, T. Wimberley, D. Cohen, O. Mors, A.D. Borglum, C. Gasse, Endogenous and Antipsychotic-Related Risks for Diabetes Mellitus in Young People With Schizophrenia: A Danish Population-Based Cohort Study, Am J Psychiatry 174(7) (2017) 686–694.

[9] P. Oriot, J.L. Feys, S. Mertens de Wilmars, A. Misson, L. Ayache, O. Fagnart, D. Gruson, A. Luts, J. Jamart, M.P. Hermans, M. Buysschaert, Insulin sensitivity, adjusted beta-cell function and adiponectinaemia among lean drug-naive schizophrenic patients treated with atypical antipsychotic drugs: a nine-month prospective study, Diabetes Metab 34(5) (2008) 490–6.

[10] B. Galling, A. Roldan, R.E. Nielsen, J. Nielsen, T. Gerhard, M. Carbon, B. Stubbs, D. Vancampfort, M. De Hert, M. Olfson, K.G. Kahl, A. Martin, J.J. Guo, H.Y. Lane, F.C. Sung, C.H. Liao, C. Arango, C.U. Correll, Type 2 Diabetes Mellitus in Youth Exposed to Antipsychotics: A Systematic Review and Meta-analysis, JAMA Psychiatry 73(3) (2016) 247–59.

[11] C.F. Ebenbichler, M. Laimer, U. Eder, B. Mangweth, E. Weiss, A. Hofer, M. Hummer, G. Kemmler, M. Lechleitner, J.R. Patsch, W.W. Fleischhacker, Olanzapine induces insulin resistance: results from a prospective study, J Clin Psychiatry 64(12) (2003) 1436–9.

[12] F.G.S. Toledo, W.F. Martin, L. Morrow, C. Beysen, D. Bajorunas, Y. Jiang, B.L. Silverman, D. McDonnell, M.N. Namchuk, J.W. Newcomer, C. Graham, Insulin and glucose metabolism with olanzapine and a combination of olanzapine and samidorphan: exploratory phase 1 results in healthy volunteers, Neuropsychopharmacology 47(3) (2022) 696–703.

[13] V. Ferreira, D. Grajales, A.M. Valverde, Adipose tissue as a target for second-generation (atypical) antipsychotics: A molecular view, Biochim Biophys Acta Mol Cell Biol Lipids 1865(2) (2020) 158534.

[14] D. Grajales, V. Ferreira, A.M. Valverde, Second-Generation Antipsychotics and Dysregulation of Glucose Metabolism: Beyond Weight Gain, Cells 8(11) (2019).

[15] S. Mukherjee, S. Skrede, E. Milbank, R. Andriantsitohaina, M. Lopez, J. Ferno, Understanding the Effects of Antipsychotics on Appetite Control, Front Nutr 8 (2021) 815456.

[16] V. Ferreira, C. Folgueira, M. Guillen, P. Zubiaur, M. Navares, A. Sarsenbayeva, P. Lopez-Larrubia, J.W. Eriksson, M.J. Pereira, F. Abad-Santos, G. Sabio, P. Rada, A.M. Valverde, Modulation of hypothalamic AMPK phosphorylation by olanzapine controls energy balance and body weight, Metabolism 137 (2022) 155335.

[17] V.L. Albaugh, C.R. Henry, N.T. Bello, A. Hajnal, S.L. Lynch, B. Halle, C.J. Lynch, Hormonal and metabolic effects of olanzapine and clozapine related to body weight in rodents, Obesity (Silver Spring) 14(1) (2006) 36–51.

[18] T. Baptista, A. Mata, L. Teneud, M. de Quijada, H.W. Han, L. Hernandez, Effects of long-term administration of clozapine on body weight and food intake in rats, Pharmacol Biochem Behav 45(1) (1993) 51–4.

[19] S. Choi, B. DiSilvio, J. Unangst, J.D. Fernstrom, Effect of chronic infusion of olanzapine and clozapine on food intake and body weight gain in male and female rats, Life Sci 81(12) (2007) 1024–30.

[20] A.J. Goudie, J.A. Smith, J.C. Halford, Characterization of olanzapine-induced weight gain in rats, J Psychopharmacol 16(4) (2002) 291–6.

[21] J. Minet-Ringuet, P.C. Even, M. Goubern, D. Tome, R. de Beaurepaire, Long term treatment with olanzapine mixed with the food in male rats induces body fat deposition with no increase in body weight and no thermogenic alteration, Appetite 46(3) (2006) 254–62.

[22] S. Skrede, J. Ferno, B. Bjorndal, W.R. Brede, P. Bohov, R.K. Berge, V.M. Steen, Lipid-lowering effects of tetradecylthioacetic acid in antipsychotic-exposed, female rats: challenges with long-term treatment, PLoS One 7(11) (2012) e50853.

[23] R. Coccurello, A. Moles, Potential mechanisms of atypical antipsychotic-induced metabolic derangement: clues for understanding obesity and novel drug design, Pharmacol Ther 127(3) (2010) 210–51.

[24] H.N. Boyda, L. Tse, R.M. Procyshyn, D. Wong, T.K. Wu, C.C. Pang, A.M. Barr, A parametric study of the acute effects of antipsychotic drugs on glucose sensitivity in an animal model, Prog Neuropsychopharmacol Biol Psychiatry 34(6) (2010) 945–54.

[25] N. Davoodi, M. Kalinichev, S.A. Korneev, P.G. Clifton, Hyperphagia and increased meal size are responsible for weight gain in rats treated sub-chronically with olanzapine, Psychopharmacology (Berl) 203(4) (2009) 693–702.

[26] M.J. Fell, N. Anjum, K. Dickinson, K.M. Marshall, L.M. Peltola, S. Vickers, S. Cheetham, J.C. Neill, The distinct effects of subchronic antipsychotic drug treatment on macronutrient selection, body weight, adiposity, and metabolism in female rats, Psychopharmacology (Berl) 194(2) (2007) 221–31.

[27] J. Wang, Q. Wu, Y. Zhou, L. Yu, L. Yu, Y. Deng, C. Tu, W. Li, The mechanisms underlying olanzapine-induced insulin resistance via the brown adipose tissue and the therapy in rats, Adipocyte 11(1) (2022) 84–98.

[28] H. Li, S. Peng, S. Li, S. Liu, Y. Lv, N. Yang, L. Yu, Y.H. Deng, Z. Zhang, M. Fang, Y. Huo, Y. Chen, T. Sun, W. Li, Chronic olanzapine administration causes metabolic syndrome through inflammatory cytokines in rodent models of insulin resistance, Sci Rep 9(1) (2019) 1582.

[29] C. Guo, J. Liu, H. Li, Metformin ameliorates olanzapine-induced insulin resistance via suppressing macrophage infiltration and inflammatory responses in rats, Biomed Pharmacother 133 (2021) 110912.

[30] P. Dipta, A. Sarsenbayeva, M. Shmuel, F. Forno, J.W. Eriksson, M.J. Pereira, X.M. Abalo, M. Wabitsch, M. Thaysen-Andersen, B. Tirosh, Macrophage-derived secretome is sufficient to confer olanzapine-mediated insulin resistance in human adipocytes, Compr Psychoneuroendocrinol 7 (2021) 100073.

[31] E.M. Girault, A. Alkemade, E. Foppen, M.T. Ackermans, E. Fliers, A. Kalsbeek, Acute peripheral but not central administration of olanzapine induces hyperglycemia associated with hepatic and extra-hepatic insulin resistance, PLoS One 7(8) (2012) e43244.

[32] L. Ren, X. Zhou, X. Huang, C. Wang, Y. Li, The IRS/PI3K/Akt signaling pathway mediates olanzapine-induced hepatic insulin resistance in male rats, Life Sci 217 (2019) 229–236.

[33] J. Engl, M. Laimer, A. Niederwanger, M. Kranebitter, M. Starzinger, M.T. Pedrini, W.W. Fleischhacker, J.R. Patsch, C.F. Ebenbichler, Olanzapine impairs glycogen synthesis and insulin signaling in L6 skeletal muscle cells, Mol Psychiatry 10(12) (2005) 1089–96.

[34] A. Gonzalez-Rodriguez, J.A. Mas Gutierrez, S. Sanz-Gonzalez, M. Ros, D.J. Burks, A.M. Valverde, Inhibition of PTP1B restores IRS1-mediated hepatic insulin signaling in IRS2-deficient mice, Diabetes 59(3) (2010) 588–99.

[35] A. Gonzalez-Rodriguez, J.A. Mas-Gutierrez, M. Mirasierra, A. Fernandez-Perez, Y.J. Lee, H.J. Ko, J.K. Kim, E. Romanos, J.M. Carrascosa, M. Ros, M. Vallejo, C.M. Rondinone, A.M. Valverde, Essential role of protein tyrosine phosphatase 1B in obesity-induced inflammation and peripheral insulin resistance during aging, Aging Cell 11(2) (2012) 284–96.

[36] A. Gonzalez-Rodriguez, B. Santamaria, J.A. Mas-Gutierrez, P. Rada, E. Fernandez-Millan, V. Pardo, C. Alvarez, A. Cuadrado, M. Ros, M. Serrano, A.M. Valverde, Resveratrol treatment restores peripheral insulin sensitivity in diabetic mice in a sirt1-independent manner, Mol Nutr Food Res 59(8) (2015) 1431–42.

[37] M. Delibegovic, D. Zimmer, C. Kauffman, K. Rak, E.G. Hong, Y.R. Cho, J.K. Kim, B.B. Kahn, B.G. Neel, K.K. Bence, Liver-specific deletion of protein-tyrosine phosphatase 1B (PTP1B) improves metabolic syndrome and attenuates diet-induced endoplasmic reticulum stress, Diabetes 58(3) (2009) 590–9.

[38] L.D. Klaman, O. Boss, O.D. Peroni, J.K. Kim, J.L. Martino, J.M. Zabolotny, N. Moghal, M. Lubkin, Y.B. Kim, A.H. Sharpe, A. Stricker-Krongrad, G.I. Shulman, B.G. Neel, B.B. Kahn, Increased energy expenditure, decreased adiposity, and tissue-specific insulin sensitivity in protein-tyrosine phosphatase 1B-deficient mice, Mol Cell Biol 20(15) (2000) 5479–89.

[39] V. Ferreira, C. Folgueira, M. Garcia-Altares, M. Guillen, M. Ruiz-Rosario, G. DiNunzio, I. Garcia-Martinez, R. Alen, C. Bookmeyer, J.G. Jones, J.C. Cigudosa, P. Lopez-Larrubia, X. Correig-Blanchar, R.J. Davis, G. Sabio, P. Rada, A.M. Valverde, Hypothalamic JNK1-hepatic fatty acid synthase axis mediates a metabolic rewiring that prevents hepatic steatosis in male mice treated with olanzapine via intraperitoneal: Additional effects of PTP1B inhibition, Redox Biol 63 (2023) 102741.

[40] S. Asslih, O. Damri, G. Agam, Neuroinflammation as a Common Denominator of Complex Diseases (Cancer, Diabetes Type 2, and Neuropsychiatric Disorders), Int J Mol Sci 22(11) (2021).

[41] I. Garcia-Ruiz, N. Blanes Ruiz, P. Rada, V. Pardo, L. Ruiz, A. Blas-Garcia, M.P. Valdecantos, M. Grau Sanz, J.A. Solis Herruzo, A.M. Valverde, Protein tyrosine phosphatase 1b deficiency protects against hepatic fibrosis by modulating nadph oxidases, Redox Biol 26 (2019) 101263.

[42] K.B.J. Franklin, G. Paxinos, Paxinos and Franklin’s The mouse brain in stereotaxic coordinates, Fourth edition. ed., Academic Press, an imprint of Elsevier, Amsterdam, 2013.

[43] M. Lopez, L. Varela, M.J. Vazquez, S. Rodriguez-Cuenca, C.R. Gonzalez, V.R. Velagapudi, D.A. Morgan, E. Schoenmakers, K. Agassandian, R. Lage, P.B. Martinez de Morentin, S. Tovar, R. Nogueiras, D. Carling, C. Lelliott, R. Gallego, M. Oresic, K. Chatterjee, A.K. Saha, K. Rahmouni, C. Dieguez, A. Vidal-Puig, Hypothalamic AMPK and fatty acid metabolism mediate thyroid regulation of energy balance, Nat Med 16(9) (2010) 1001–8.

[44] P.B. Martinez de Morentin, I. Gonzalez-Garcia, L. Martins, R. Lage, D. Fernandez-Mallo, N. Martinez-Sanchez, F. Ruiz-Pino, J. Liu, D.A. Morgan, L. Pinilla, R. Gallego, A.K. Saha, A. Kalsbeek, E. Fliers, P.H. Bisschop, C. Dieguez, R. Nogueiras, K. Rahmouni, M. Tena-Sempere, M. Lopez, Estradiol regulates brown adipose tissue thermogenesis via hypothalamic AMPK, Cell Metab 20(1) (2014) 41–53.

[45] N. Martinez-Sanchez, P. Seoane-Collazo, C. Contreras, L. Varela, J. Villarroya, E. Rial-Pensado, X. Buque, I. Aurrekoetxea, T.C. Delgado, R. Vazquez-Martinez, I. Gonzalez-Garcia, J. Roa, A.J. Whittle, B. Gomez-Santos, V. Velagapudi, Y.C.L. Tung, D.A. Morgan, P.J. Voshol, P.B. Martinez de Morentin, T. Lopez-Gonzalez, L. Linares-Pose, F. Gonzalez, K. Chatterjee, T. Sobrino, G. Medina-Gomez, R.J. Davis, N. Casals, M. Oresic, A.P. Coll, A. Vidal-Puig, J. Mittag, M. Tena-Sempere, M.M. Malagon, C. Dieguez, M.L. Martinez-Chantar, P. Aspichueta, K. Rahmouni, R. Nogueiras, G. Sabio, F. Villarroya, M. Lopez, Hypothalamic AMPK-ER Stress-JNK1 Axis Mediates the Central Actions of Thyroid Hormones on Energy Balance, Cell Metab 26(1) (2017) 212–229 e12.

[46] M. Crespo, I. Nikolic, A. Mora, E. Rodriguez, L. Leiva-Vega, A. Pintor-Chocano, D. Horrillo, L. Hernandez-Cosido, J.L. Torres, E. Novoa, R. Nogueiras, G. Medina-Gomez, M. Marcos, M. Leiva, G. Sabio, Myeloid p38 activation maintains macrophage-liver crosstalk and BAT thermogenesis through IL-12-FGF21 axis, Hepatology 77(3) (2023) 874–887.

[47] Y.E. Savoy, M.A. Ashton, M.W. Miller, F.M. Nedza, D.K. Spracklin, M.H. Hawthorn, H. Rollema, F.F. Matos, E. Hajos-Korcsok, Differential effects of various typical and atypical antipsychotics on plasma glucose and insulin levels in the mouse: evidence for the involvement of sympathetic regulation, Schizophr Bull 36(2) (2010) 410–8.

[48] H.N. Boyda, A. Ramos-Miguel, R.M. Procyshyn, E. Topfer, N. Lant, H.H. Choy, R. Wong, L. Li, C.C. Pang, W.G. Honer, A.M. Barr, Routine exercise ameliorates the metabolic side-effects of treatment with the atypical antipsychotic drug olanzapine in rats, Int J Neuropsychopharmacol 17(1) (2014) 77–90.

[49] V. Mondelli, C. Anacker, A.C. Vernon, A. Cattaneo, S. Natesan, M. Modo, P. Dazzan, S. Kapur, C.M. Pariante, Haloperidol and olanzapine mediate metabolic abnormalities through different molecular pathways, Transl Psychiatry 3 (2013) e208.

[50] A. Calevro, M.C. Cotel, S. Natesan, M. Modo, A.C. Vernon, V. Mondelli, Effects of chronic antipsychotic drug exposure on the expression of Translocator Protein and inflammatory markers in rat adipose tissue, Psychoneuroendocrinology 95 (2018) 28–33.

[51] M. Ikegami, H. Ikeda, T. Ohashi, M. Kai, M. Osada, A. Kamei, J. Kamei, Olanzapine-induced hyperglycemia: possible involvement of histaminergic, dopaminergic and adrenergic functions in the central nervous system, Neuroendocrinology 98(3) (2013) 224–32.

[52] V. Ferreira, C. Folgueira, A. Montes-San Lorenzo, A. Rodriguez-Lopez, E. Gonzalez-Iglesias, P. Zubiaur, F. Abad-Santos, G. Sabio, P. Rada, A.M. Valverde, Estrogens prevent the hypothalamus-periphery crosstalk induced by olanzapine intraperitoneal treatment in female mice: Effects on brown/beige adipose tissues and liver, Biochim Biophys Acta Mol Basis Dis 1870(5) (2024) 167227.

[53] P. Rada, E. Carceller-Lopez, A.B. Hitos, B. Gomez-Santos, C. Fernandez-Hernandez, E. Rey, J. Pose-Utrilla, C. Garcia-Monzon, A. Gonzalez-Rodriguez, G. Sabio, A. Garcia, P. Aspichueta, T. Iglesias, A.M. Valverde, Protein kinase D2 modulates hepatic insulin sensitivity in male mice, Mol Metab 90 (2024) 102045.

[54] P. Rada, A. Mosquera, J. Muntane, F. Ferrandiz, L. Rodriguez-Manas, F. de Pablo, J. Gonzalez-Canudas, A.M. Valverde, Differential effects of metformin glycinate and hydrochloride in glucose production, AMPK phosphorylation and insulin sensitivity in hepatocytes from non-diabetic and diabetic mice, Food Chem Toxicol 123 (2019) 470–480.

[55] P. Kolli, G. Kelley, M. Rosales, J. Faden, R. Serdenes, Olanzapine Pharmacokinetics: A Clinical Review of Current Insights and Remaining Questions, Pharmgenomics Pers Med 16 (2023) 1097–1108.

[56] J.H.M. Yung, A. Giacca, Role of c-Jun N-terminal Kinase (JNK) in Obesity and Type 2 Diabetes, Cells 9(3) (2020).

[57] J.C. Bruning, M.D. Michael, J.N. Winnay, T. Hayashi, D. Horsch, D. Accili, L.J. Goodyear, C.R. Kahn, A muscle-specific insulin receptor knockout exhibits features of the metabolic syndrome of NIDDM without altering glucose tolerance, Mol Cell 2(5) (1998) 559–69.

[58] K.E. Merz, D.C. Thurmond, Role of Skeletal Muscle in Insulin Resistance and Glucose Uptake, Compr Physiol 10(3) (2020) 785–809.

[59] S. Vernia, J. Cavanagh-Kyros, L. Garcia-Haro, G. Sabio, T. Barrett, D.Y. Jung, J.K. Kim, J. Xu, H.P. Shulha, M. Garber, G. Gao, R.J. Davis, The PPARalpha-FGF21 hormone axis contributes to metabolic regulation by the hepatic JNK signaling pathway, Cell Metab 20(3) (2014) 512–25.

[60] V. Jimenez, C. Jambrina, E. Casana, V. Sacristan, S. Munoz, S. Darriba, J. Rodo, C. Mallol, M. Garcia, X. Leon, S. Marco, A. Ribera, I. Elias, A. Casellas, I. Grass, G. Elias, T. Ferre, S. Motas, S. Franckhauser, F. Mulero, M. Navarro, V. Haurigot, J. Ruberte, F. Bosch, FGF21 gene therapy as treatment for obesity and insulin resistance, EMBO Mol Med 10(8) (2018).

[61] C.C. Lord, S.C. Wyler, R. Wan, C.M. Castorena, N. Ahmed, D. Mathew, S. Lee, C. Liu, J.K. Elmquist, The atypical antipsychotic olanzapine causes weight gain by targeting serotonin receptor 2C, J Clin Invest 127(9) (2017) 3402–3406.

[62] S. Skrede, I. Gonzalez-Garcia, L. Martins, R.K. Berge, R. Nogueiras, M. Tena-Sempere, G. Mellgren, V.M. Steen, M. Lopez, J. Ferno, Lack of Ovarian Secretions Reverts the Anabolic Action of Olanzapine in Female Rats, Int J Neuropsychopharmacol 20(12) (2017) 1005–1012.

[63] Y. Levkovitz, G. Ben-Shushan, A. Hershkovitz, R. Isaac, I. Gil-Ad, D. Shvartsman, D. Ronen, A. Weizman, Y. Zick, Antidepressants induce cellular insulin resistance by activation of IRS-1 kinases, Mol Cell Neurosci 36(3) (2007) 305–12.

[64] J. Wu, C. Mei, H. Vlassara, G.E. Striker, F. Zheng, Oxidative stress-induced JNK activation contributes to proinflammatory phenotype of aging diabetic mesangial cells, Am J Physiol Renal Physiol 297(6) (2009) F1622–31.

[65] G. Verdile, K.N. Keane, V.F. Cruzat, S. Medic, M. Sabale, J. Rowles, N. Wijesekara, R.N. Martins, P.E. Fraser, P. Newsholme, Inflammation and Oxidative Stress: The Molecular Connectivity between Insulin Resistance, Obesity, and Alzheimer’s Disease, Mediators Inflamm 2015 (2015) 105828.

[66] G. Solinas, B. Becattini, JNK at the crossroad of obesity, insulin resistance, and cell stress response, Mol Metab 6(2) (2017) 174–184.

[67] G. Sabio, R.J. Davis, cJun NH2-terminal kinase 1 (JNK1): roles in metabolic regulation of insulin resistance, Trends Biochem Sci 35(9) (2010) 490–6.

[68] R. Nogueiras, G. Sabio, Brain JNK and metabolic disease, Diabetologia 64(2) (2021) 265–274.

[69] B.F. Belgardt, J. Mauer, F.T. Wunderlich, M.B. Ernst, M. Pal, G. Spohn, H.S. Bronneke, S. Brodesser, B. Hampel, A.C. Schauss, J.C. Bruning, Hypothalamic and pituitary c-Jun N-terminal kinase 1 signaling coordinately regulates glucose metabolism, Proc Natl Acad Sci U S A 107(13) (2010) 6028–33.

[70] G. Sabio, R.J. Davis, TNF and MAP kinase signalling pathways, Semin Immunol 26(3) (2014) 237–45.

[71] K.D. Copps, M.F. White, Regulation of insulin sensitivity by serine/threonine phosphorylation of insulin receptor substrate proteins IRS1 and IRS2, Diabetologia 55(10) (2012) 2565–2582.

[72] H. Li, C. Wang, J. Zhao, C. Guo, JNK downregulation improves olanzapine-induced insulin resistance by suppressing IRS1(Ser307) phosphorylation and reducing inflammation, Biomed Pharmacother 142 (2021) 112071.

[73] S. Li, T. Zou, J. Chen, J. Li, J. You, Fibroblast growth factor 21: An emerging pleiotropic regulator of lipid metabolism and the metabolic network, Genes Dis 11(3) (2024) 101064.

[74] C. Schlein, S. Talukdar, M. Heine, A.W. Fischer, L.M. Krott, S.K. Nilsson, M.B. Brenner, J. Heeren, L. Scheja, FGF21 Lowers Plasma Triglycerides by Accelerating Lipoprotein Catabolism in White and Brown Adipose Tissues, Cell Metab 23(3) (2016) 441–53.

[75] M. Elchebly, P. Payette, E. Michaliszyn, W. Cromlish, S. Collins, A.L. Loy, D. Normandin, A. Cheng, J. Himms-Hagen, C.C. Chan, C. Ramachandran, M.J. Gresser, M.L. Tremblay, B.P. Kennedy, Increased insulin sensitivity and obesity resistance in mice lacking the protein tyrosine phosphatase-1B gene, Science 283(5407) (1999) 1544–8.

[76] J.M. Zabolotny, Y.B. Kim, L.A. Welsh, E.E. Kershaw, B.G. Neel, B.B. Kahn, Protein-tyrosine phosphatase 1B expression is induced by inflammation in vivo, J Biol Chem 283(21) (2008) 14230–41.

[77] M. Delibegovic, K.K. Bence, N. Mody, E.G. Hong, H.J. Ko, J.K. Kim, B.B. Kahn, B.G. Neel, Improved glucose homeostasis in mice with muscle-specific deletion of protein-tyrosine phosphatase 1B, Mol Cell Biol 27(21) (2007) 7727–34.

[78] Z. Qin, L. Zhang, M.A. Zasloff, A.F.R. Stewart, H.H. Chen, Ketamine’s schizophrenia-like effects are prevented by targeting PTP1B, Neurobiol Dis 155 (2021) 105397.

[79] C. Kowalchuk, C. Teo, V. Wilson, A. Chintoh, L. Lam, S.M. Agarwal, A. Giacca, G.J. Remington, M.K. Hahn, In male rats, the ability of central insulin to suppress glucose production is impaired by olanzapine, whereas glucose uptake is left intact, J Psychiatry Neurosci 42(6) (2017) 424–431.

[80] E. Mattiuz, R. Franklin, T. Gillespie, A. Murphy, J. Bernstein, A. Chiu, T. Hotten, K. Kassahun, Disposition and metabolism of olanzapine in mice, dogs, and rhesus monkeys, Drug Metab Dispos 25(5) (1997) 573–83.

[81] M. Aravagiri, Y. Teper, S.R. Marder, Pharmacokinetics and tissue distribution of olanzapine in rats, Biopharm Drug Dispos 20(8) (1999) 369–77.

[82] J.T. Callaghan, R.F. Bergstrom, L.R. Ptak, C.M. Beasley, Olanzapine. Pharmacokinetic and pharmacodynamic profile, Clin Pharmacokinet 37(3) (1999) 177–93.

