## Supplementary data and figures for "A hypothalamus-liver-skeletal muscle axis controlled by JNK1 and FGF21 mediates olanzapine-induced insulin resistance in an intraperitoneal treatment in male mice"

**Supplementary Materials and Methods**

### 1.1. **Glucose, insulin and pyruvate tolerance tests**

For glucose tolerance test (GTT), mice were fasted overnight and i.p. injected D-glucose (2 g/Kg; G8270, Sigma, Madrid, Spain) and blood glucose was measured before and at 15, 30, 60 and 120 min after glucose injection with a glucometer (AccuCheck Aviva, SB_RDC_2019_06, Roche Diagnostics, Rotkreuz, Switzerland). For insulin tolerance test (ITT), mice were fasted for 4 h and then i.p. injected 0.75 U/Kg of human insulin (Actrapid, 775502, Novo Nordisk, Bagsværd, Dinamarca). Blood glucose was measured before and at 15, 30, 60 and 120 min post-injection. For pyruvate tolerance test (PTT), mice were fasted overnight and i.p. injected sodium pyruvate (1.5 g/Kg; #P2256, Sigma) and blood glucose was measured before and after 15, 30, 60 and 120 min. The area under the curve (AUC) was calculated.

### 1.2. **Primary culture of mouse hepatocytes**

Primary hepatocytes were isolated from mice treated with OLA or vehicle (VEH), or from naive C57BL/6J mice by using a modified two-step collagenase perfusion method [1, 2]. The livers were first perfused with a calcium-free HBSS solution to disrupt cell-cell junctions, followed by a second perfusion with a collagenase (C5138, Sigma-Aldrich)-containing E-William’s media to digest the tissue into a suspension of single cells. The livers were perfused through the inferior vena cava, with the perfusate flowing toward the heart. Partially digested livers were placed in petri dishes, minced and disaggregated in Attachment Media (AM, DMEM and Ham’s F-12 medium (1:1) with 10 % heat-inactivated fetal bovine serum (FBS, 10270-106, Life Technologies, Carlsbad, CA, USA) supplemented with 2 mM glutamine (25030-024, Life Technologies), 15 mM glucose, 20 mM HEPES, 100 U/ml penicillin, 100 μg/ml streptomycin (15140-122, Life Technologies) and 1 mM sodium pyruvate (11360-039, Life Technologies). After that, liver homogenates were filtered with 100-µm cell strainers, centrifuged at 50 *x g* for 5 min at RT and supernatants containing non-parenchymal cells were discarded. Pellets containing hepatocytes were resuspended in AM. An isotonic Percoll solution (Percoll, 17-0891-01, GE Healthcare, HBSS 1x, 75 mM NaCl) was used to purify viable hepatocytes by exploiting density differences between live and dead cells. Viable hepatocytes were pelleted by centrifugation at 200 *x g* for 10 min, RT, and then, were then resuspended in AM, washed by centrifugation at at 50 *x g* for 5 min, RT, and then cells were cultured in 6- or 12-well collagen IV (C3867-1VL, Sigma-Aldrich) pre-coated plates in AM for 24 h. Then, medium was replaced by DMEM w/o phenol red, w/o glucose (11966-025, Life Technologies) supplemented with 5.5 mM glucose for 1 h and after that, primary hepatocytes were stimulated with 10 nM insulin for 15 min.

Primary hepatocytes isolated from non-treated mice were exposed to 12.5 µM OLA for 48 h in 5.5 mM glucose DMEM and then stimulated with 10 nM insulin for 15 min to test insulin response. To analyze JNK phosphorylation, hepatocytes were exposed to 12.5 µM OLA for 30 to 240 min in 5.5 mM glucose DMEM.

### 1.3. **Glucose uptake assay**

After 24 h in attachment medium, primary mouse hepatocytes were maintained in FBS-free 5 mM glucose DMEM for 2 h. Afterwards, cells were washed with ice-cold Krebs Ringer-phosphate buffer (KRP) (135 mM NaCl, 5.4 mM KCl, 1.4 mM CaCl_2_, 1.4 mM MgSO_4_, 10 mM sodium pyrophosphate, pH 7.4) and 2-Deoxy-D [1-^3^H] glucose (500 nCi/ml) in KRP was added for 10 min at 37 °C. Cells were then washed with ice-cold KRP buffer and solubilized in 1% sodium dodecyl sulfate (SDS) as previously described [3]. Total protein was determined by using the Pierce™ BCA Protein Assay Kit (#23225, Thermo Fisher Scientific, Massachusetts, USA) with bovine serum albumin (BSA) as standard. Glucose uptake was expressed as cpm per µg of protein.

### 1.4. **Tissue sample collection**

At week 8 (for the i.p. treatment) or 30 min, 8 h and 48 h post-injection (for the intrahypothalamic treatment), mice were euthanized by decapitation. Blood was collected with EDTA to prevent clotting, and immediately centrifuged for 20 min at 15 600 g at 4 ºC. Plasma was collected, frozen and stored at -80 ºC. The brain was removed from the skull and the hypothalamus was isolated and snap-frozen. The liver and skeletal muscle were dissected and snap-frozen. The brain and liver samples were placed in 4 % (w/v) paraformaldehyde (PFA, 16005, Sigma-Aldrich) for paraffin inclusion. For the analysis of insulin signaling *in vivo*, mice were fasted for 4 h and then i.p. injected 0.75 U/Kg of human insulin for 15 min before sacrifice.

### 1.5. **Protein extraction from cells and tissues**

To obtain total cell lysates, cells were scraped off in lysis buffer (40 mM Tris-HCl, 5 mM EDTA, 30 μM sodium pyrophosphate tetrabasic (Na_4_P_2_O_7_), 50 mM sodium fluoride (NaF), 100 mM sodium o-vanadate (Na_3_VO_4_), 1% Triton X-100, 1 mM phenylmethylsulphonyl fluoride (PMSF), and 10 μg/ml protease inhibitors cocktail, at final pH 7.4-7.6). Afterwards, samples were cleared by centrifugation at 13 600 g for 20 min at 4 ºC. Supernatants were stored at -20 ºC. Protein concentration was measured by the Bradford dye (#5000006, Bio-Rad Laboratories, Hercules, CA, USA) method using BSA as standard.

To extract total protein from tissues (hypothalamus, liver and skeletal muscle), samples were homogenized in ice-cold lysis buffer (50 mM HEPES, 1 % Triton X-100, 50 mM Na_4_P_2_O_7_, 0.1 M NaF, 10 mM EDTA, 10 mM Na_4_P_2_O_7_, 1 mM PMSF, and 10 μg/ml of protease inhibitors cocktail, at final pH 7.4-7.6) using a polytron tissue homogenizer (0003737000, IKA, Staufen, Germany). Extracts were kept on ice during the process. Extracts were cleared twice by centrifugation at 40 000 g for 40 min at 4 ºC. Supernatants were then aliquoted and stored at -80 ºC. Protein quantification was performed by the BCA dye method using BCA Protein Assay Kit (23227, Thermo Fisher Scientific, Waltham, MA, USA) and BSA as standard.

### 1.6. **Western blot**

After protein content determination, samples were prepared with equal amount of protein, boiled at 98 ºC for 5 min and subjected to SDS-polyacrylamide gel electrophoresis (SDS-PAGE). Then gels were transferred to PVDF membranes (Millipore, Burlington, MA, USA) previously activated with 100 % methanol for 5 min. Afterwards membranes were blocked using 4 % non-fat dried milk in TTBS (10 mM Tris-HCl, 150 mM NaCl pH 7.5, Tween 20 (0.05% w/v)) for 2 h, and incubated overnight at 4 ºC with primary antibodies (Supplementary Table 1) in TTBS. Membranes were then washed and incubated with the appropriate secondary antibody diluted in blocking solution. Immunoreactive bands were visualized in a ChemiDoc™ MP digital imager (Bio-Rad Laboratories) or with radiographic films in a radiology cassette (AGFA, Mortsel, Belgium) by using the Clarity Western ECL Substrate (#170-5061, Bio-Rad Laboratories). Blots were normalized using antibodies against housekeeping proteins. Densitometry values were determined using Fiji Software [4].

1.7. **Quantitative Real-Time PCR analysis (RT-qPCR)**

Total RNA from tissues and cells was extracted with Tri reagent (AM9738, Life Technologies) and reverse transcribed using a SuperScript III First-Strand Synthesis System (18080051, Life Technologies) for real time quantitative PCR (RT-qPCR) following the manufacturer’s indications. RT-qPCR was performed with an ABI 7900 sequence detector (Applied Biosystems, Waltham. MA, USA). The primers sequences/probes are included in Supplementary Table 2. Data analysis is based on the ΔΔCt method with normalization of the raw data to housekeeping genes as described in the manufacturer's manual (Applied Biosystems).

### 1.8. **Tissue histological analysis, immunohistochemistry and immunofluorescence**

**Immunohistochemistry:** Tissue sections (5 µm) were de-waxed and rehydrated through a descending series of ethanol dilutions (100, 96, 75, 50 %). Antigen retrieval was achieved by microwaving for 5 min at 600 W in 100 mM citrate buffer (pH 6) with 0.05 % (w/v) Tween-20, followed by three washes in phosphate buffered saline (PBS). Tissue sections were permeabilized with PBS-Triton X-100 (0.3 % w/v; T8787, Sigma-Aldrich). Non-specific binding was blocked with 3 % (v/v) horse serum (H0146, Sigma-Aldrich) in PBS-Triton X-100 (0.3 % v/v). Sections were incubated overnight at 4 °C with primary antibodies (Supplementary Table 3) in blocking buffer and then incubated with a biotinylated secondary antibody combined with streptavidin-HRP by the ABC (Avidin Biotin Complex) method. DAB kit was used following the manufacturer’s instructions. Images were collected with an Axiophot light microscope (Carl Zeiss, Oberkochen, Germany) with 20x or 40x objectives.

**Immunofluorescence in tissue sections:** After rehydrating and antigen retrieval, non-specific binding was blocked with 0.5 % (w/v) BSA (MB04602, NZYTech, Lisbon, Portugal) in PBS-Triton X-100 (0.3 % v/v). Sections were incubated overnight at 4 °C with the corresponding primary antibody (Supplementary Table 3) in blocking buffer and then incubated with the appropriated fluorescent secondary antibody. Nuclei were stained for 5 min with 4,6-diamidino-2-phenylindole (DAPI) in PBS (1:1000) and sections were mounted in Fluoromont (Sigma-Aldrich). Samples were analyzed using confocal microscopy (Carl Zeiss).

***Hematoxylin and eosin (H&E) staining:*** H&E staining was performed in paraffin-embedded liver sections. Liver was fixed in 4% (w/v) PFA for 24 h, washed twice with PBS, dehydrated with ascending ethanol solutions, incubated with xylene and then embedded in paraffin. Blocks were cut in 5 μm sections. Prior to H&E staining, sections were deparaffinized in xylene and hydrated in descending ethanol solutions and distilled water. The slides were stained with Mayer's hematoxylin (MHS32-1L, Sigma-Aldrich) for 15 min and eosin (1.15935.0025, Merck) for 1 min. After mounted and dried, images were captured with an Axiophot light microscope (Carl Zeiss) using a 20x objective.

***Sirius red staining:*** Sirius red staining was performed in paraffin-embedded liver sections. Liver was fixed in 4% (w/v) PFA for 24 h, washed twice with PBS, dehydrated with ascending ethanol solutions, incubated with xylene and then embedded in paraffin. Blocks were cut in 5 μm sections. Prior to Sirius red staining, sections were deparaffinized in xylene and hydrated in descending ethanol solutions and distilled water. Slides were submerged in Picro-Sirius red solution for 1 h and, next, they were washed twice with acidified water (0.5 % acetic acid in tap water). Samples were then washed with distilled water, dehydrated with ascending ethanol solutions and cleared in xylene. After mounted in a resinous medium and dried, images were captured with an Axiophot light microscope (Carl Zeiss) using a 20x objective.

1.9. **Analysis of plasma insulin levels**

Plasma insulin concentrations were determined with a Mouse Insulin ELISA kit (10-1247-01; Mercodia, Sweden) following the manufacturer’s indications. Briefly, plasma samples were loaded on the ELISA plate and incubated for 2 h in an orbital shaker at RT. After washing 6 times, 200 µL of substrate solution were added to each well and incubated for 15 min at RT before adding 50 µL of stop solution. The absorbance of each well was measured by using a microplate reader set to 450 nm. Afterwards, these values were used to assess insulin plasma levels in plasma samples from WT mice chronically treated with OLA (10 mg/kg) or VEH daily for 8 weeks.

### 1.10. **Analysis of Fibroblast Growth Factor 21 (FGF21) plasma levels.**

FGF21 plasma levels were assessed by an ELISA kit for mouse and rat plasma samples (RD291108200R, R&D Systems) following the manufacturer’s indications. Briefly, samples were loaded on the ELISA plate and incubated for 1 h in an orbital shaker (300 rpm) at RT. After washing 3 times, 100 µL of the biotin labelled antibody solution was added into each well and incubated for 1 h in the orbital shaker (300 rpm) at RT. Subsequently, wells were washed for 3 times and incubated for 30 min in the orbital shaker (300 rpm) at RT with 100 µL Streptavidin-HRP conjugate solution. After washing 3 times, 100 µL of substrate solution were added to each well and incubated for 10 min at RT before adding 100 µL of stop solution. The absorbance of each well was measured by using a microplate reader set to 450 nm, with the reference wavelength set to 630 nm. Afterwards, readings at 630 nm were subtracted from the readings at 450 nm and these values were used to assess FGF21 levels in plasma samples from WT and PTP1B-KO mice chronically treated with OLA (10 mg/kg) or VEH daily for 8 weeks.

**References**

[1] A. Gonzalez-Rodriguez, J.A. Mas Gutierrez, S. Sanz-Gonzalez, M. Ros, D.J. Burks, A.M. Valverde, Inhibition of PTP1B restores IRS1-mediated hepatic insulin signaling in IRS2-deficient mice, Diabetes, 59 (2010) 588-599.

[2] R. Benveniste, T.M. Danoff, J. Ilekis, H.R. Craig, Epidermal growth factor receptor numbers in male and female mouse primary hepatocyte cultures, Cell Biochem Funct, 6 (1988) 231-235.

[3] E. Garcia-Casarrubios, C. de Moura, A.I. Arroba, N. Pescador, M. Calderon-Dominguez, L. Garcia, L. Herrero, D. Serra, S. Cadenas, F. Reis, E. Carvalho, M.J. Obregon, A.M. Valverde, Rapamycin negatively impacts insulin signaling, glucose uptake and uncoupling protein-1 in brown adipocytes, Biochim Biophys Acta, 1861 (2016) 1929-1941.

[4] J. Schindelin, I. Arganda-Carreras, E. Frise, V. Kaynig, M. Longair, T. Pietzsch, S. Preibisch, C. Rueden, S. Saalfeld, B. Schmid, J.Y. Tinevez, D.J. White, V. Hartenstein, K. Eliceiri, P. Tomancak, A. Cardona, Fiji: an open-source platform for biological-image analysis, Nat Methods, 9 (2012) 676-682.

**Supplementary figures**

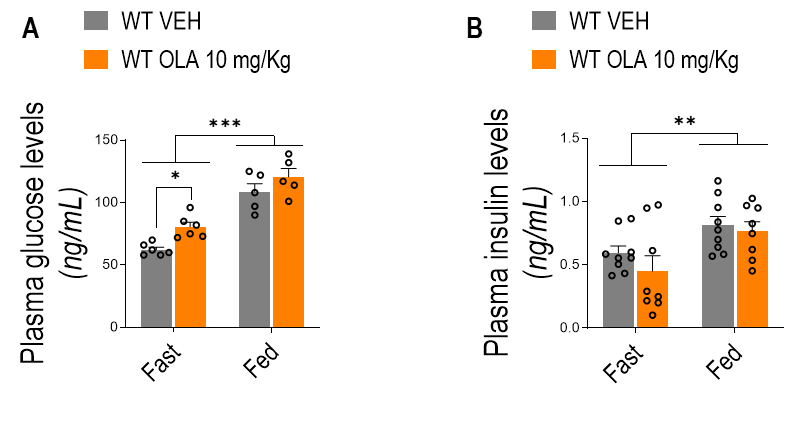

**Supplementary figure 1. *Plasma glucose and insulin levels of male mice receiving an OLA i.p. treatment for 8 weeks.* A.** Plasma glucose levels in fast (16-18 h) and fed state of mice receiving OLA via i.p. daily for 8 weeks (WT VEH: n= 5-6; WT OLA: n=5-6). **B.** Plasma insulin levels in fast (16-18 h) and fed state of mice receiving OLA via i.p. daily for 8 weeks (WT VEH: n=9; WT OLA: n=8). Each point/bar corresponds to mean ± SEM; comparisons between groups: ^⁎^p < 0.05; ^⁎⁎^p < 0.01; ^⁎⁎⁎^p < 0.001.

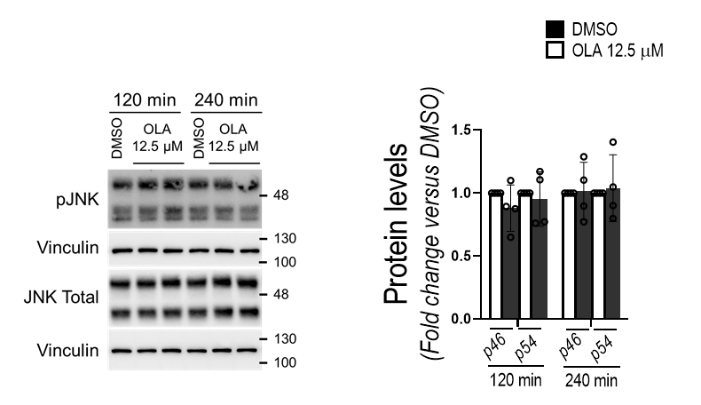

**Supplementary Figure 2. *OLA treatment does not affect JNK phosphorylation in primary mouse hepatocytes. A*.** Representative Western blots of JNK phosphorylation in primary hepatocytes treated *in vitro* with OLA (12.5 µM) for several time periods and densitometric quantification normalized for total JNK and vinculin levels (DMSO: n=4; OLA: n=4). Each point/bar corresponds to mean ± SEM.

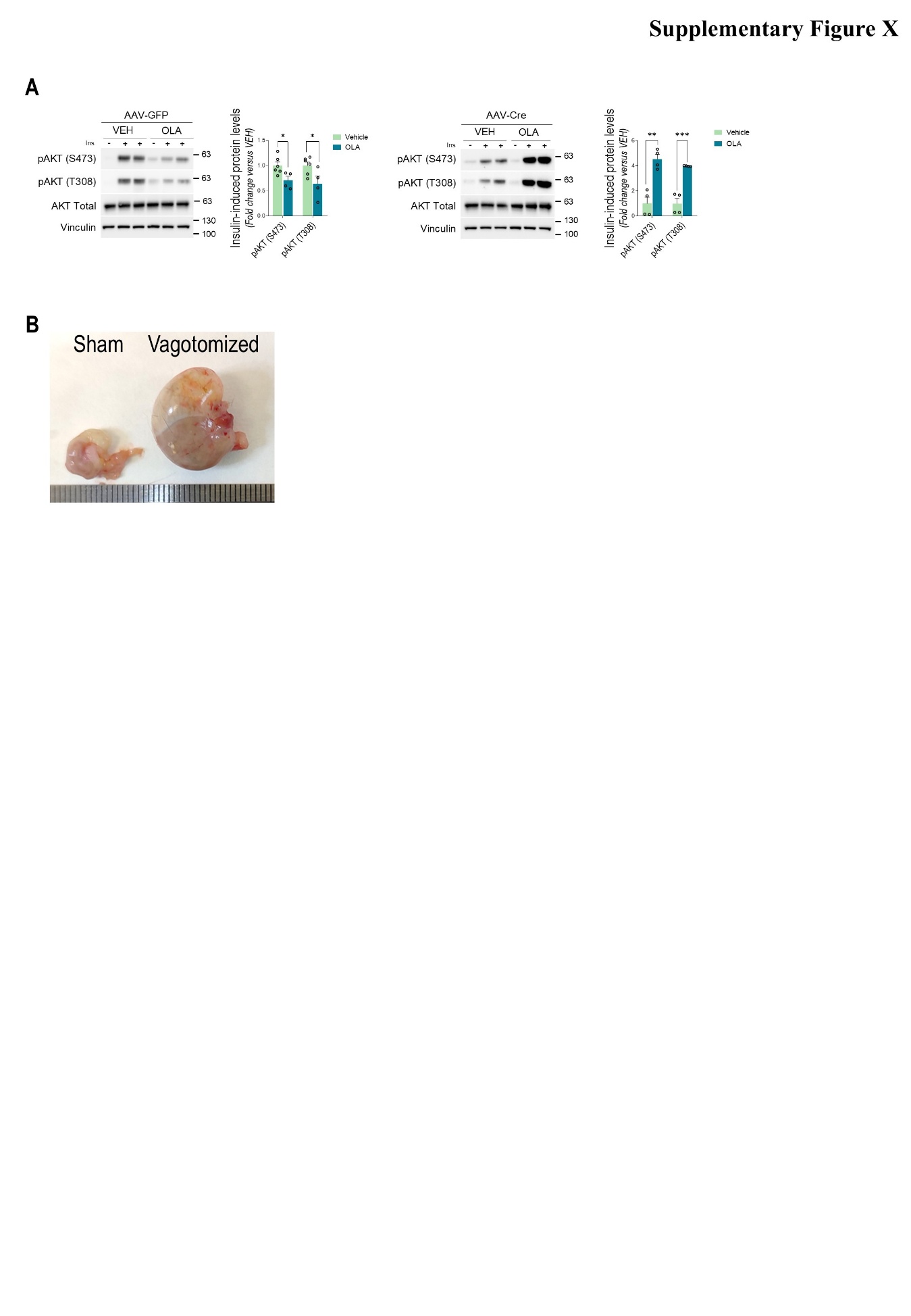
**Supplementary Figure 3. *Hypothalamic JNK1 deletion prevents OLA-induced attenuation in hepatic insulin signaling in male mice.* A*.*** Representative Western blots of insulin-induced AKT phosphorylation (Ser473 and Thr308) in liver of AAV-GFP control mice (left panel) or mice with hypothalamic JNK1 deletion (AAV-Cre, right panel) receiving VEH or OLA via i.p. Densitometric quantification normalized for total AKT and vinculin levels (AAV-GFP VEH: n=6; AAV-GFP OLA: n=4; AAV-Cre VEH: n=4; AAV-Cre OLA: n=4). **B.** Stomachs from vagotomized and sham-operated mice showing an evident increase in size after vagotomy due to motoric dysfunction. Each point/bar corresponds to mean ± SEM; comparisons between groups: ^⁎^p < 0.05; ^⁎⁎^p < 0.01; ^⁎⁎⁎^p < 0.001.

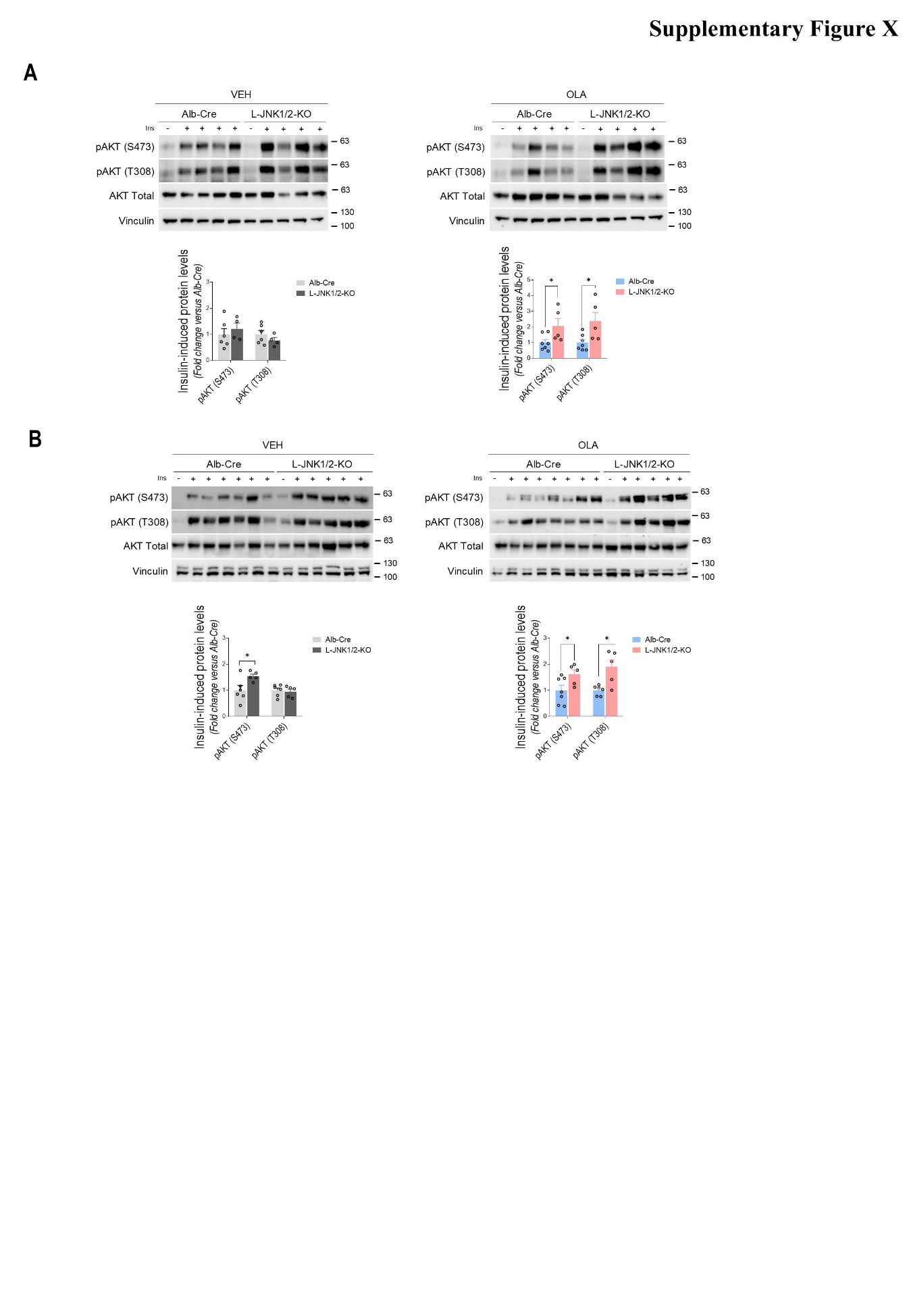

**Supplementary Figure 4. *Hepatic JNK1 deletion prevents OLA-induced attenuation in liver and skeletal muscle insulin signaling in male mice.* A*.*** Representative Western blots of insulin-induced AKT phosphorylation (Ser473 and Thr308) in liver of mice with liver JNK1 deletion (L-JNK1/2-KO) or Alb-Cre control mice receiving VEH (left panel) or OLA (right panel) via intrahypothalamic. Densitometric quantification normalized for total AKT and vinculin levels (Alb-Cre VEH: n=6; L-JNK1/2-KO VEH: n=4; Alb-Cre OLA: n=5; L-JNK1/2-KO OLA: n=5). **B*.*** Representative Western blots of insulin-induced AKT phosphorylation (Ser473 and Thr308) in skeletal muscle of mice with liver JNK1 deletion (L-JNK1/2-KO) or Alb-Cre control mice receiving VEH (left panel) or OLA (right panel) via intrahypothalamic. Densitometric quantification normalized for total AKT and vinculin levels (Alb-Cre VEH: n=6; L-JNK1/2-KO VEH: n=5; Alb-Cre OLA: n=5; L-JNK1/2-KO OLA: n=5). Each point/bar corresponds to mean ± SEM; comparisons between groups: ^⁎^p < 0.05.

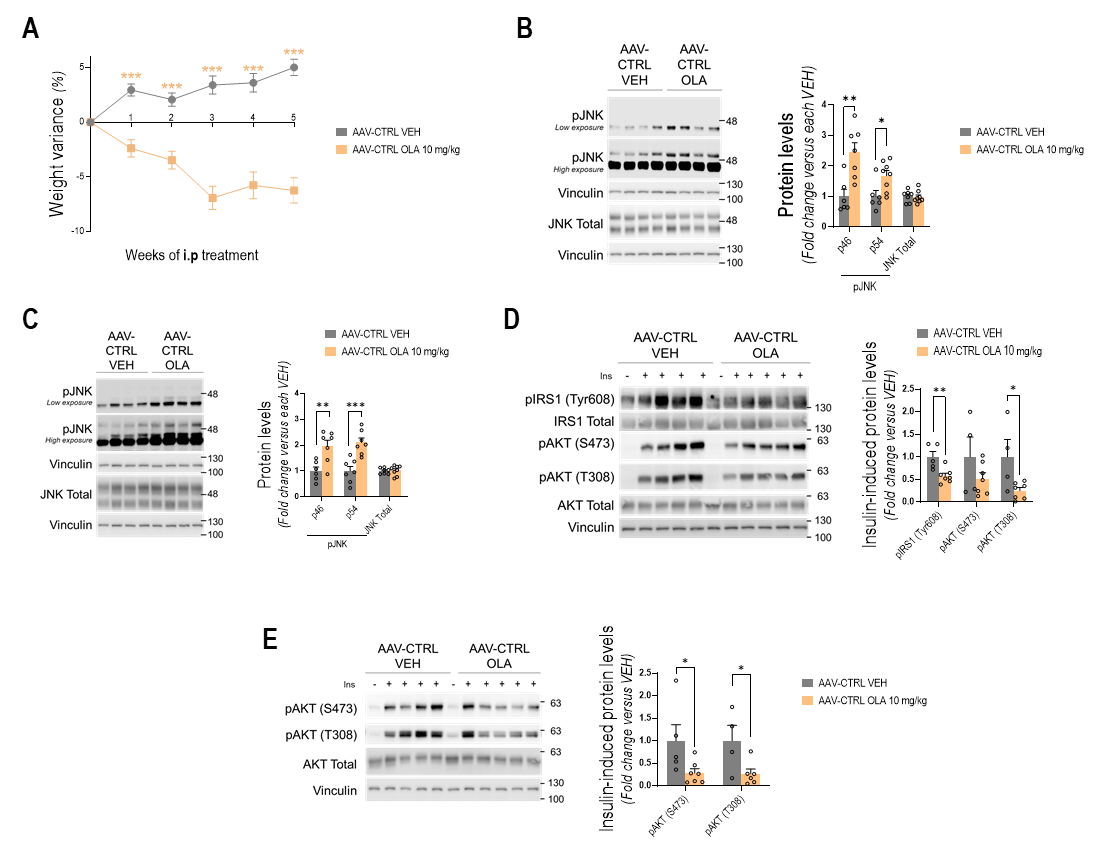

**Supplementary Figure 5. *The effect of an i.p. treatment with OLA in systemic and hepatic insulin sensitivity in AAV-CTRL injected male mice is similar to that of WT mice.* A.** Body weight change of mice injected AAV-CTRL and receiving OLA via i.p. (AAV-CTRL VEH: n= 7; AAV-CTRL OLA: n= 8). **B.** Representative Western blots of JNK phosphorylation in the hypothalamus of AAV-CTRL injected mice treated with OLA via i.p. and densitometric quantification normalized for total JNK and vinculin levels (AAV-CTRL VEH: n= 6-7; AAV-CTRL OLA: n= 7-8). **C.** Representative Western blots of JNK phosphorylation in liver of AAV-CTRL injected mice treated with OLA via i.p. and densitometric quantification normalized for total JNK and vinculin levels (AAV-CTRL VEH: n= 6-7; AAV-CTRL OLA: n= 7-8). **D.** Representative Western blots of insulin-induced IRS1 phosphorylation in Tyr608 residues and AKT phosphorylation (Set473 and Thr308) in liver of AAV-CTRL mice receiving OLA via i.p. and stimulated with insulin (0.75 U/kg) for the last 15 min, and densitometric quantification normalized for total IRS1, AKT and vinculin levels (AAV-CTRL VEH: n= 4-5; AAV-CTRL OLA: n= 7). **E.** Representative Western blots of insulin-induced AKT phosphorylation (Ser473 and Thr308) in skeletal muscle of AAV-CTRL injected mice receiving OLA via i.p. and stimulated with insulin (0.75 U/kg) for the last 15 min, and densitometric quantification normalized for total IRS1, AKT and vinculin levels (AAV-CTRL VEH: n= 4-5; AAV-CTRL OLA: n= 7). Each point/bar corresponds to mean ± SEM; comparisons between groups: ^⁎^p < 0.05; ^⁎⁎^p < 0.01; ^⁎⁎⁎^p < 0.001.

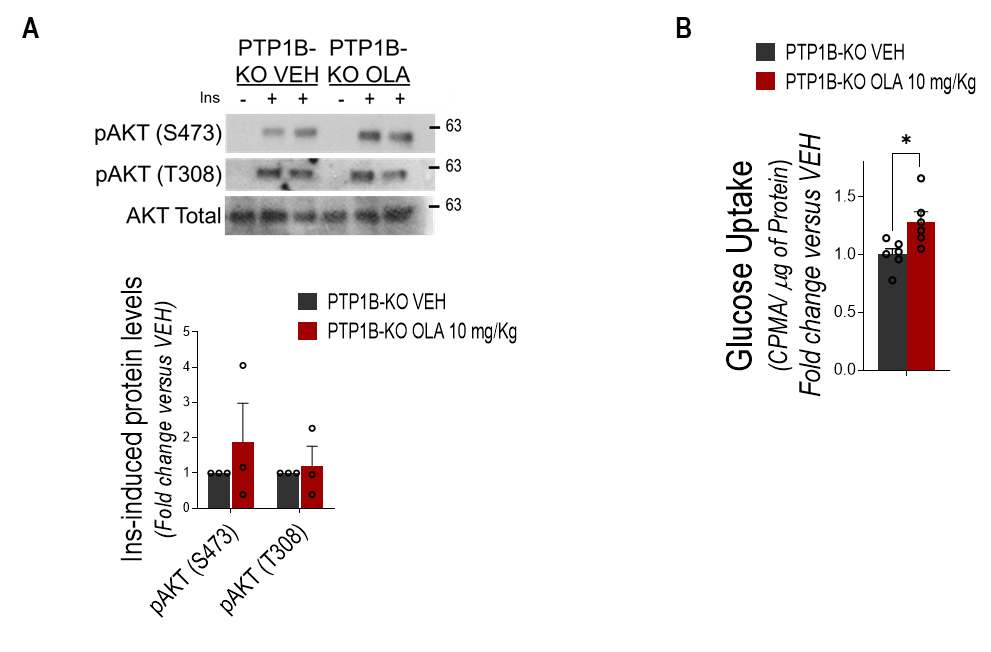

**Supplementary Figure 6. *PTP1B-deficient hepatocytes are protected against insulin resistance induced by the treatment with OLA via i.p.* A.** Representative Western blots of insulin-induced AKT (Ser473 and Thr308) phosphorylation in primary hepatocytes isolated from VEH- or OLA-treated PTP1B-KO mice via i.p. and stimulated with insulin (10 nM, 15 min). Densitometric quantification normalized for total AKT levels (PTP1B-KO VEH: n=3; PTP1B-KO OLA: n=2). **F.** Glucose uptake (fold of change versus VEH) of primary hepatocytes from OLA-treated male mice (PTP1B-KO VEH: n=6; PTP1B-KO OLA: n=6). Each point/bar corresponds to mean ± SEM; comparisons between groups: ^⁎^p < 0.05; ^⁎⁎^p < 0.01; ^⁎⁎⁎^p < 0.001.

**Uncropped Western blots membranes**

**Figure 1D**

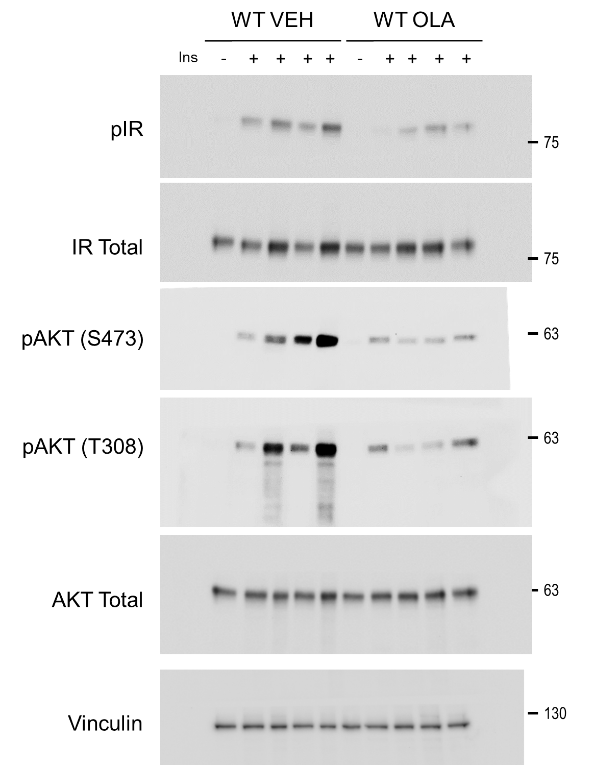

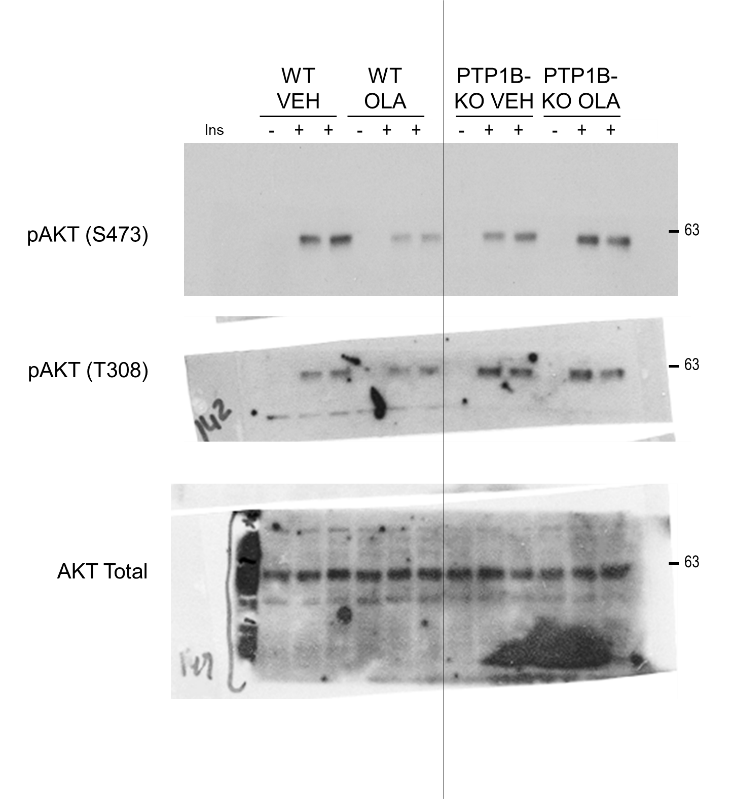

| **Figure 1E** | **Sup. Figure 6A** |
| --- | --- |

**Figure 2A**

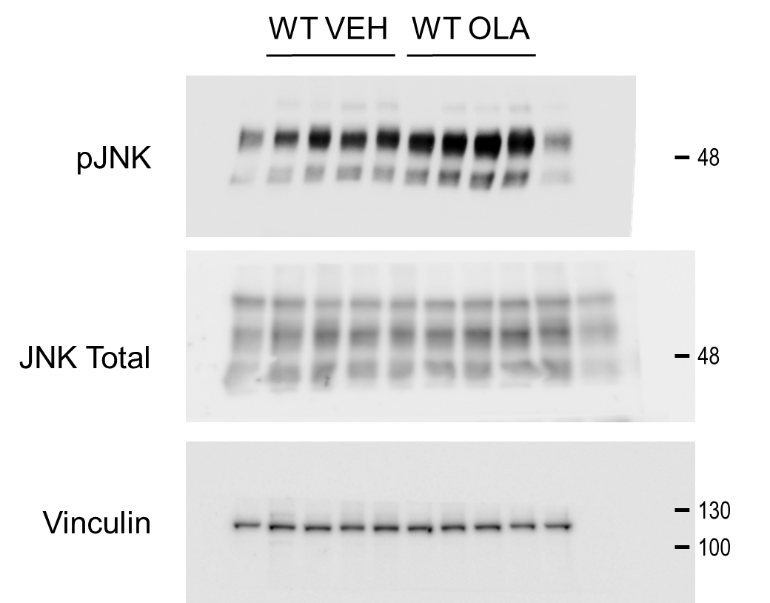

**Figure 2B**

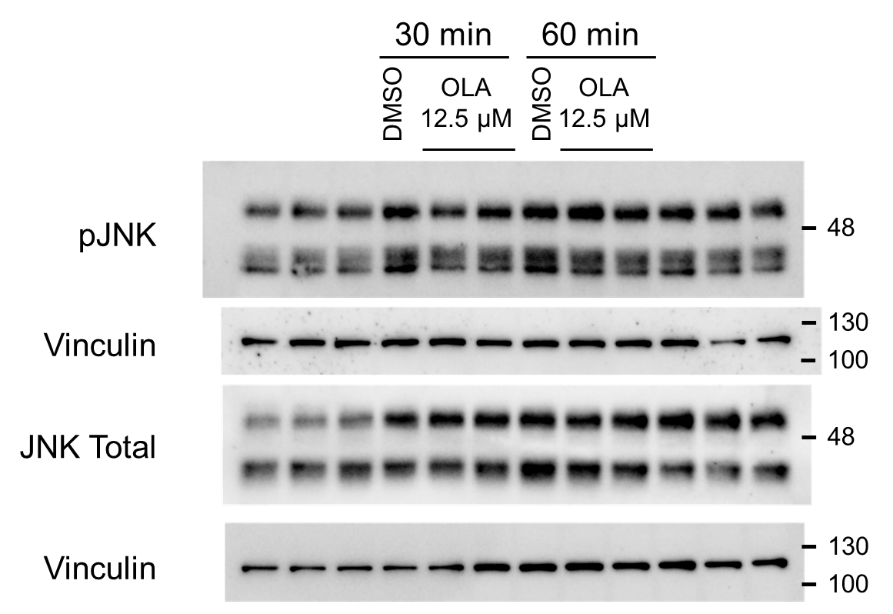

**Figure 3A**

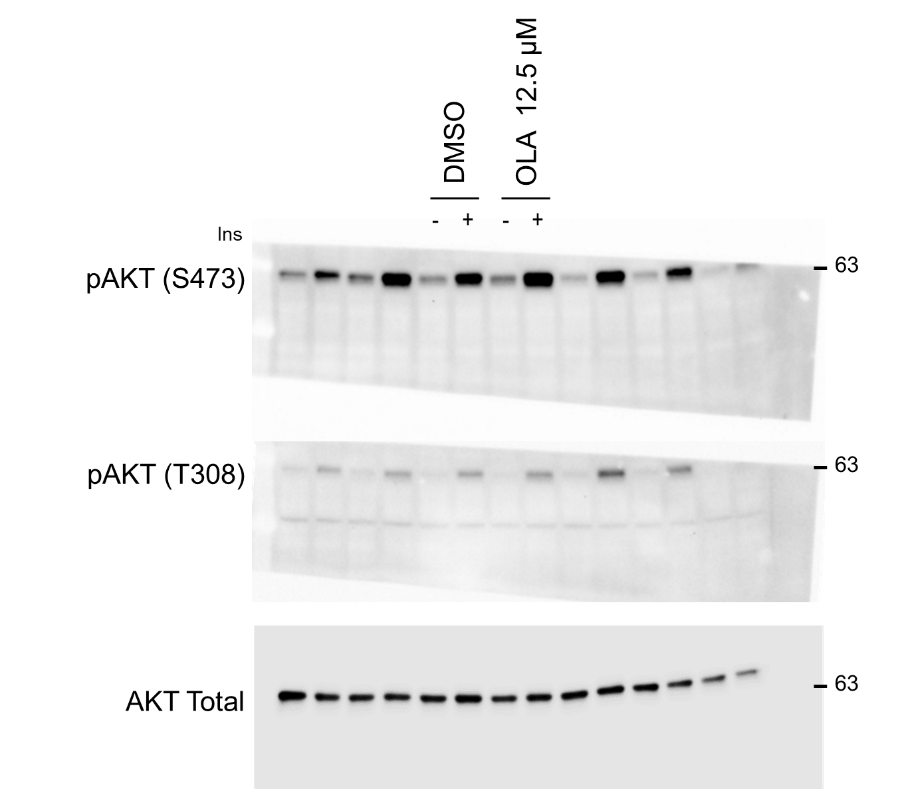

**Figure 3B**

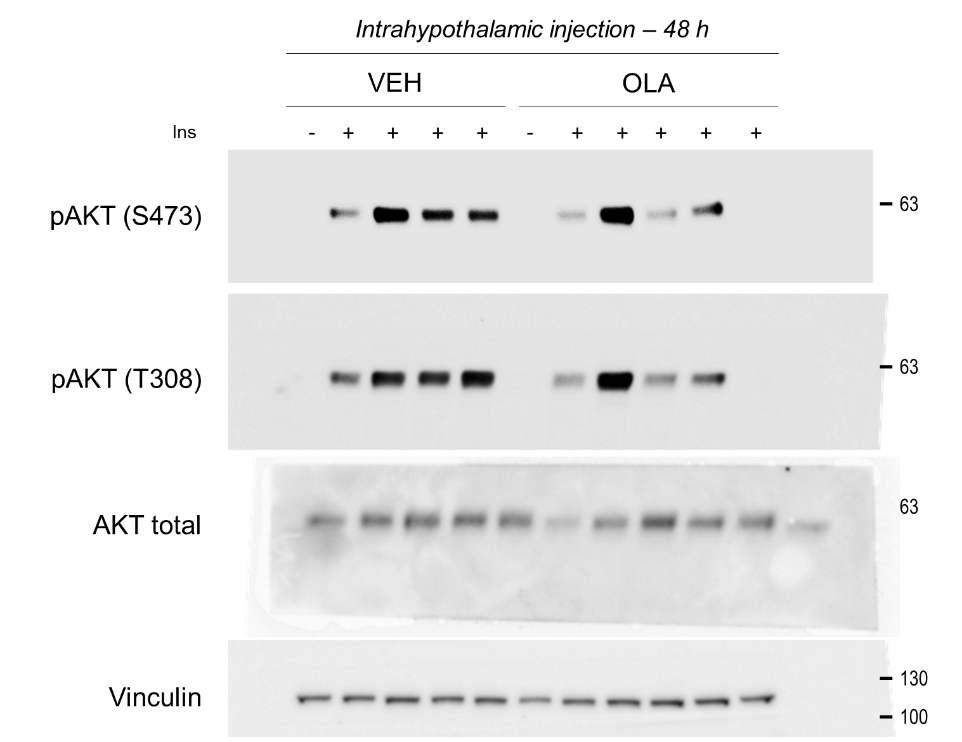

**Figure 3C**

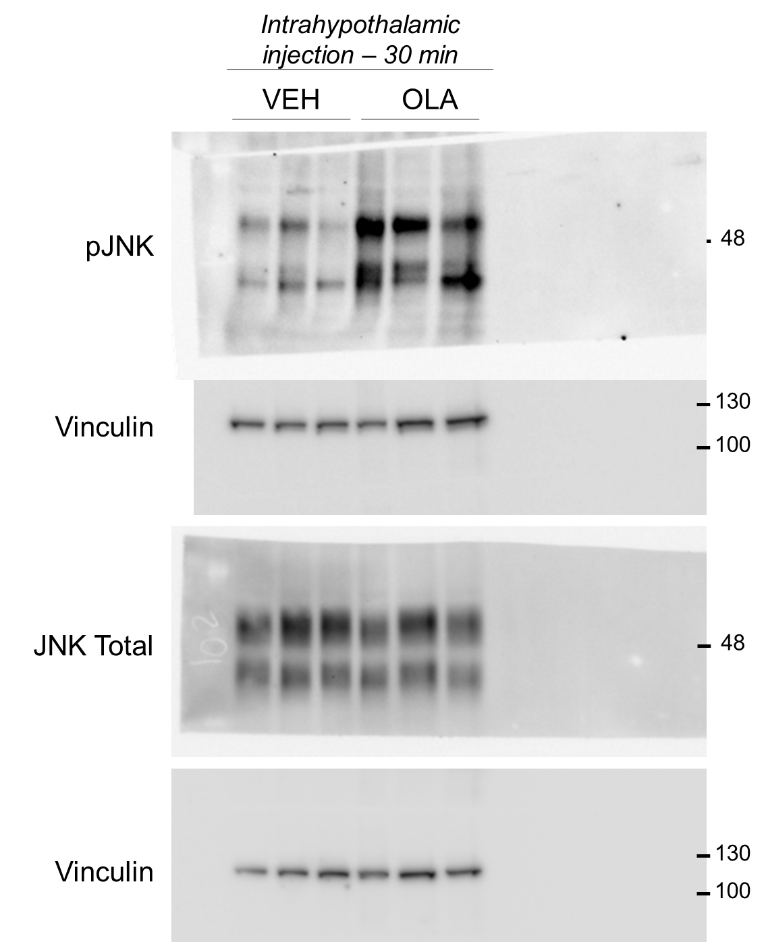

**Figure 3D**

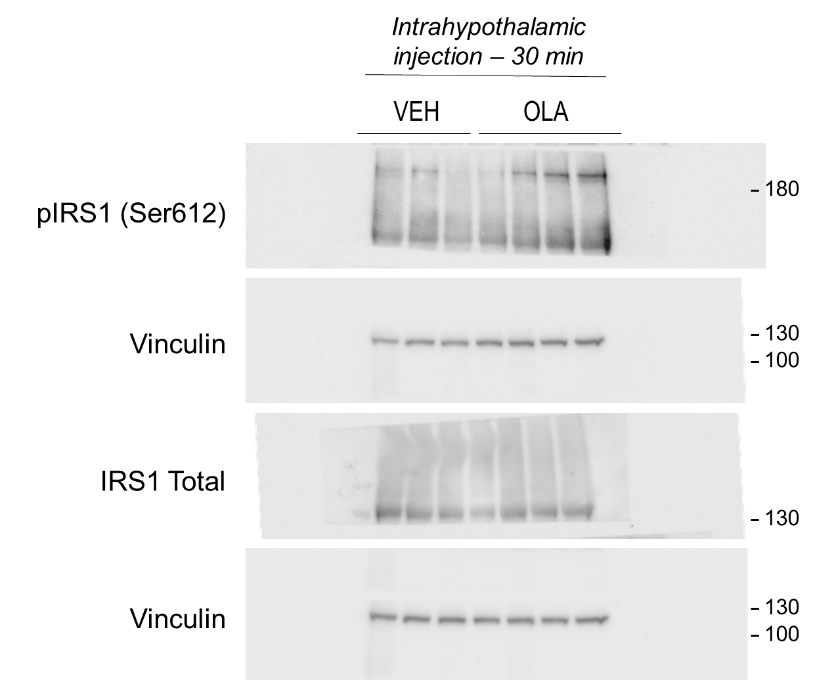

**Figure 3E**

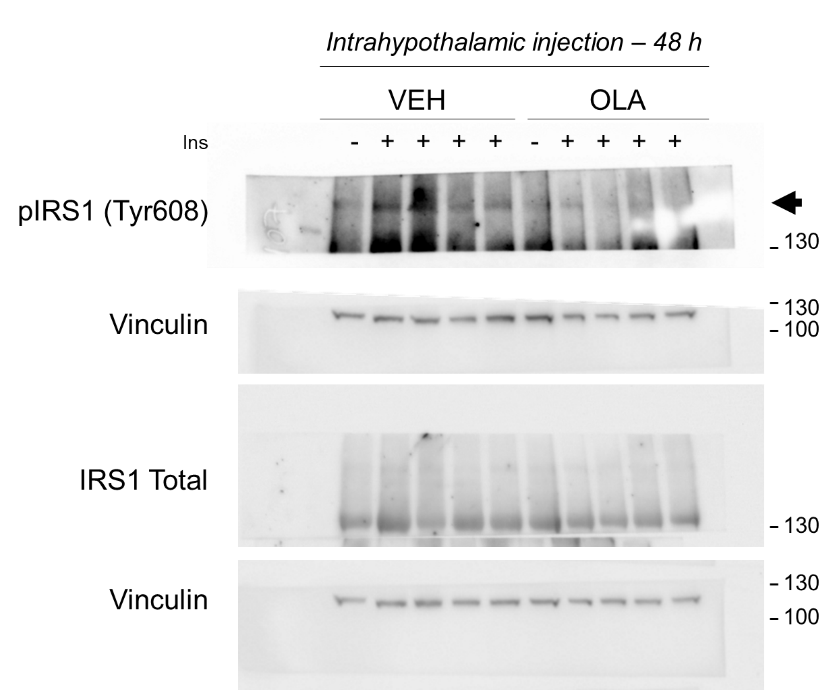

**Figure 3F**

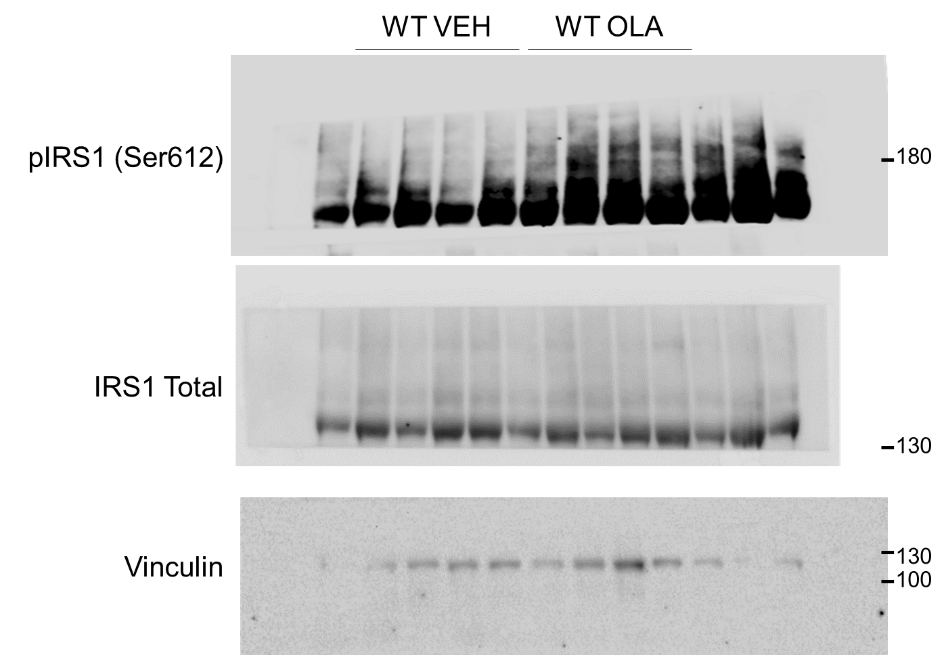

**Figure 3G**

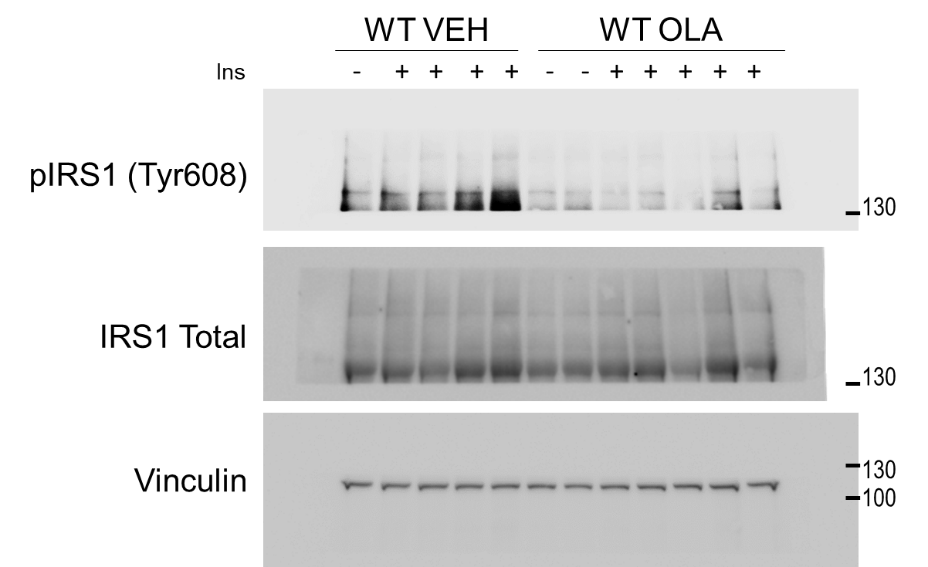

**Figure 4A**

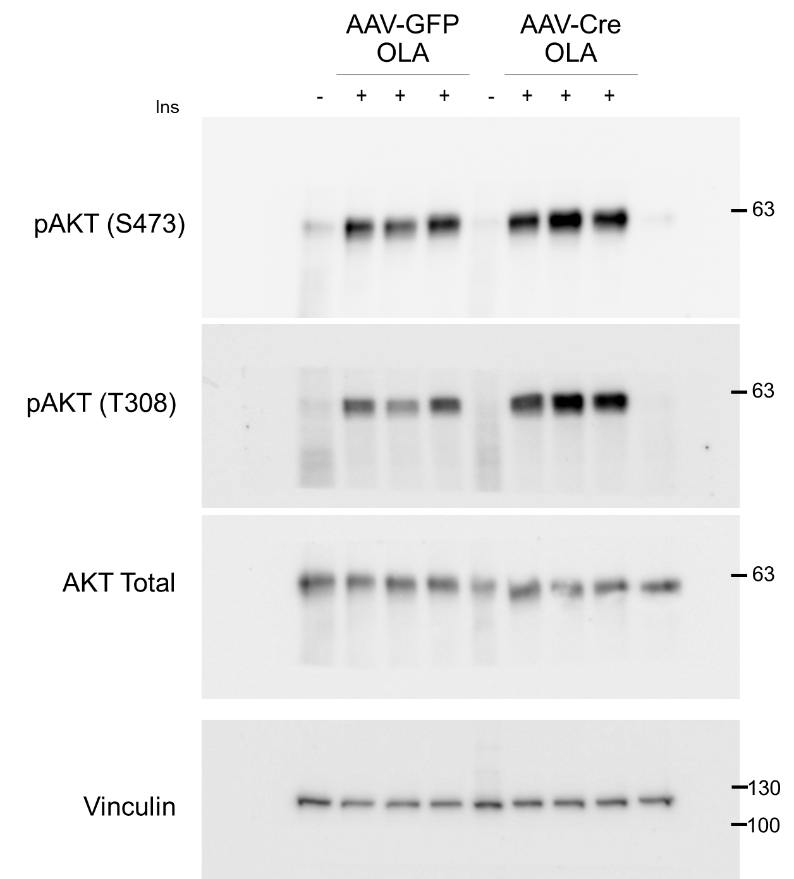

**Figure 4B**

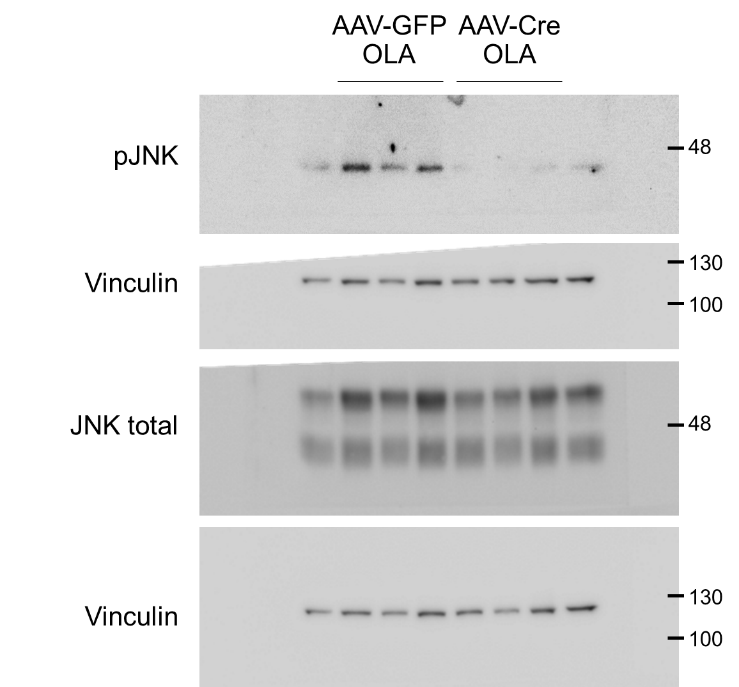

**Figure 4C**

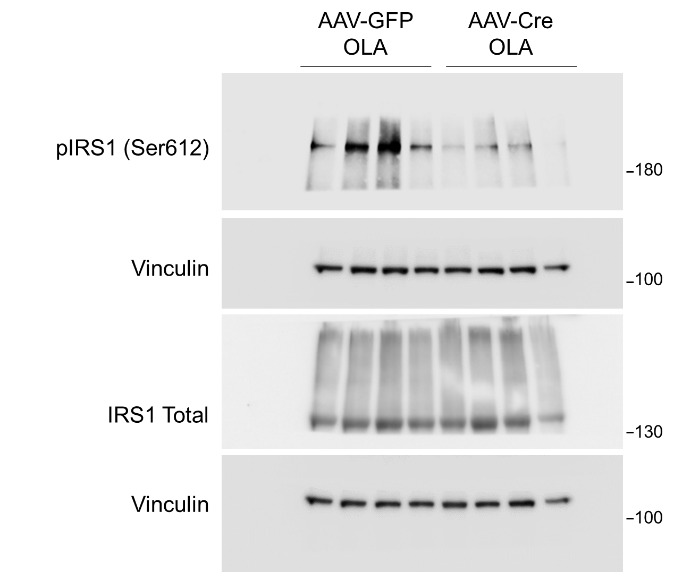

**Figure 4D**

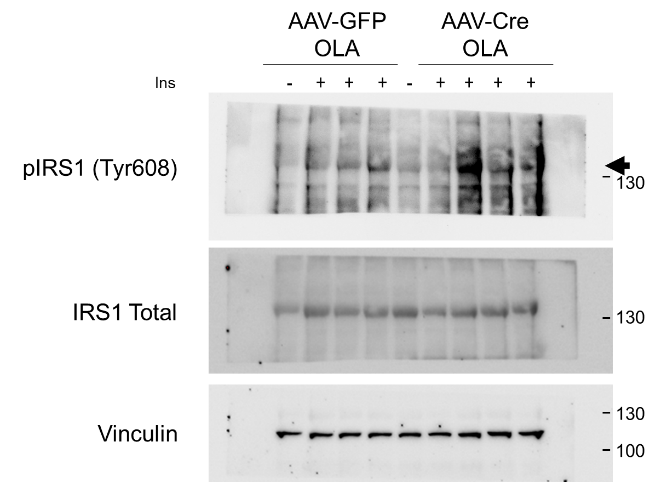

**Figure 4E**

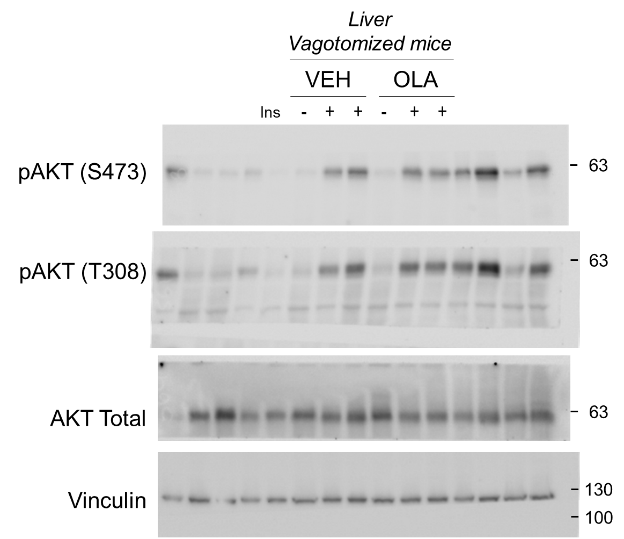

**Figure 4F**

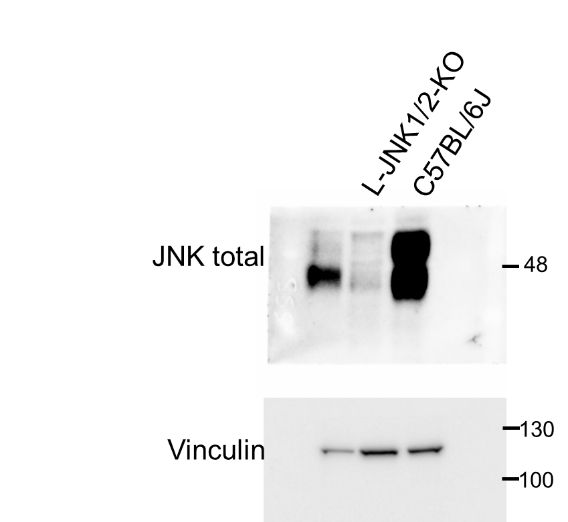

**Figure 4G**

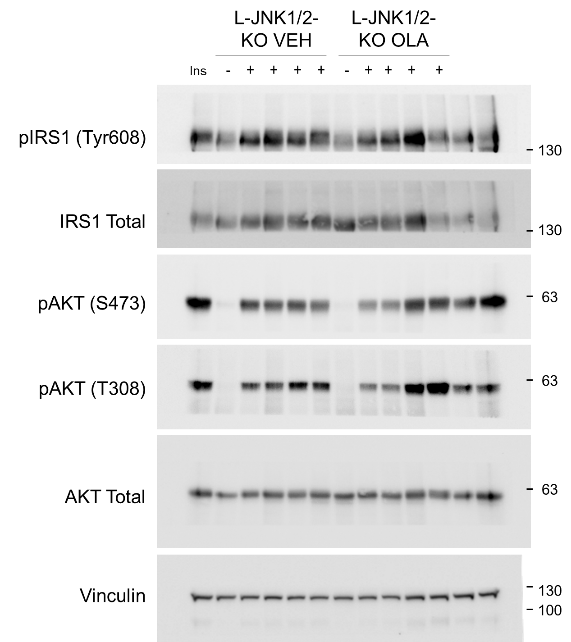

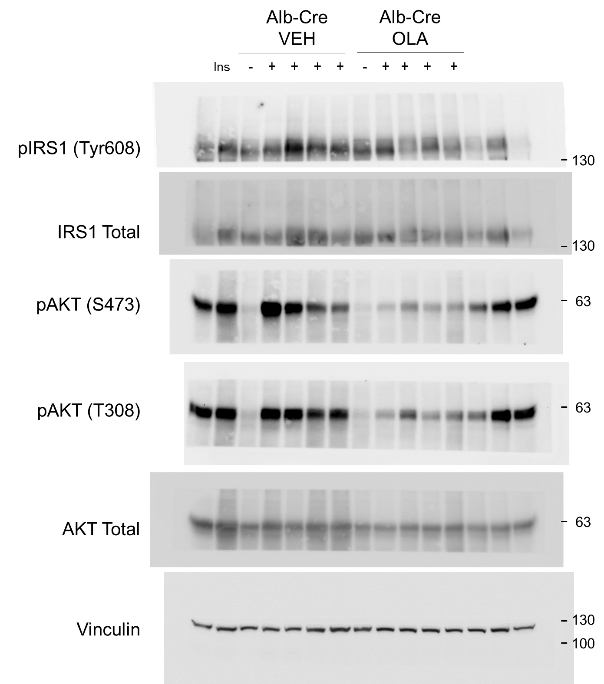

**Figure 5A**

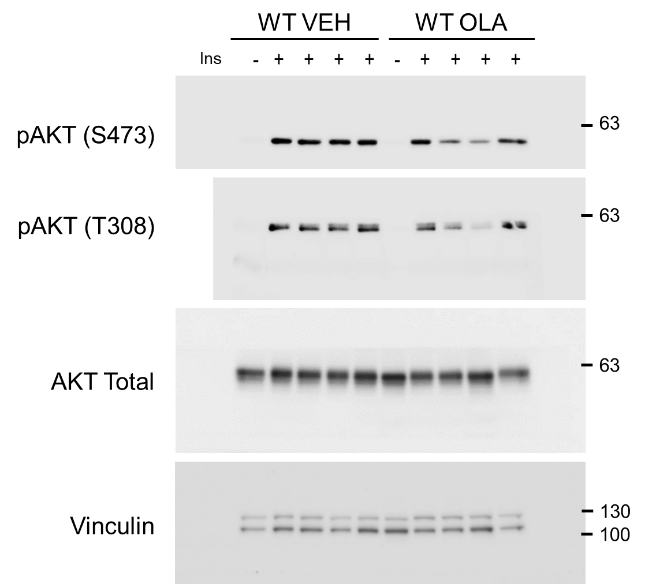

**Figure 5B**

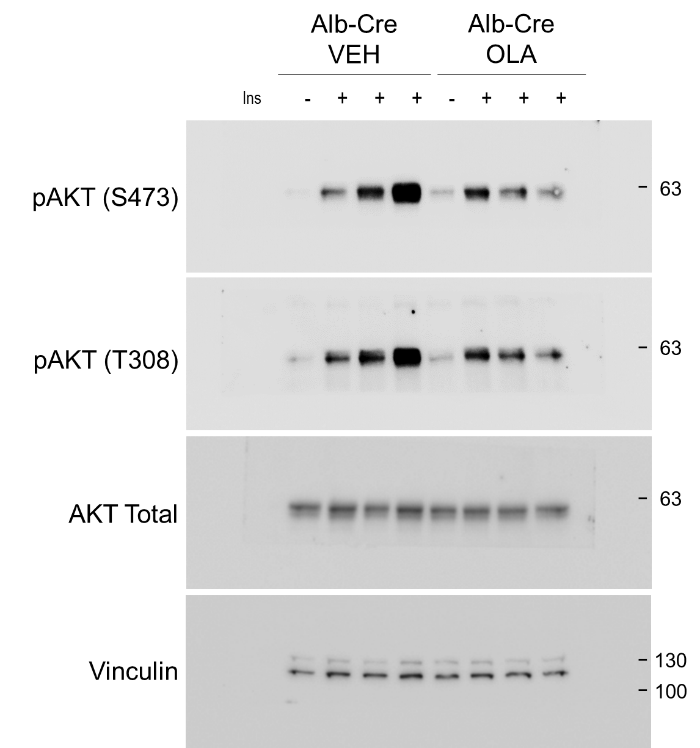

**Figure 5C**

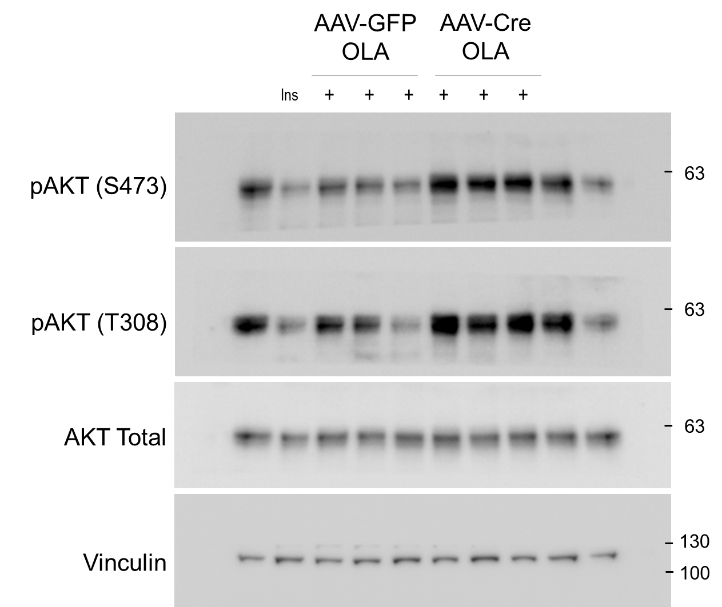

**Figure 5D**

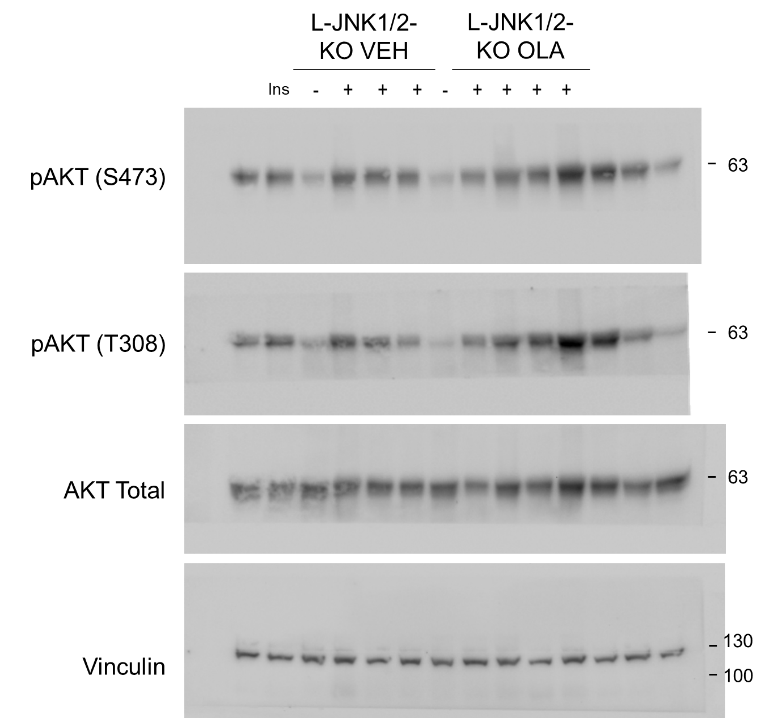

**Figure 5E**

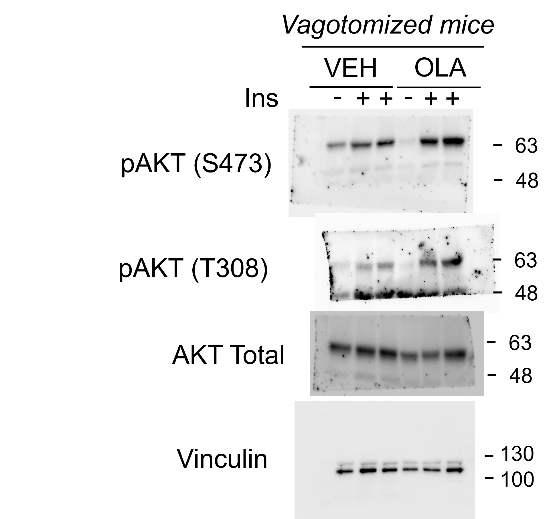

**Figure 6F**

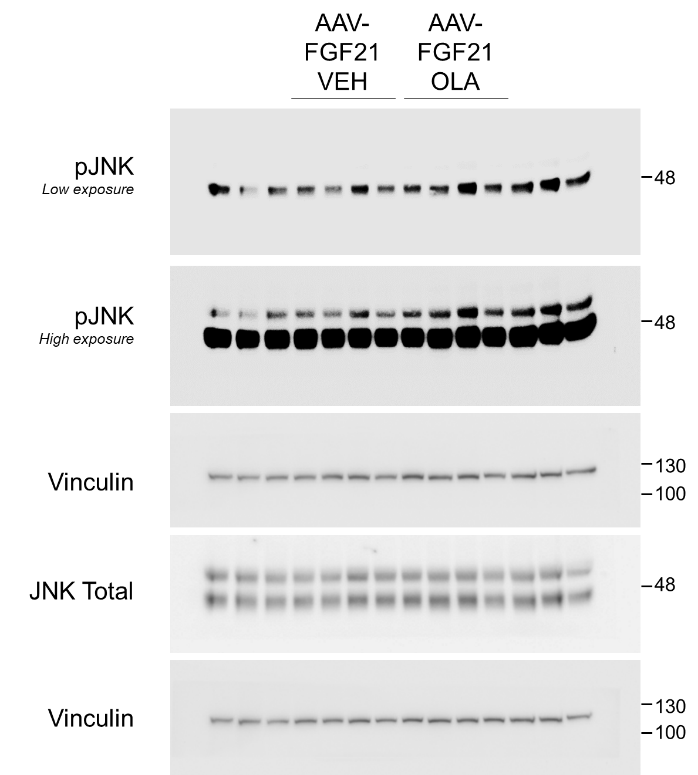

**Figure 6G**

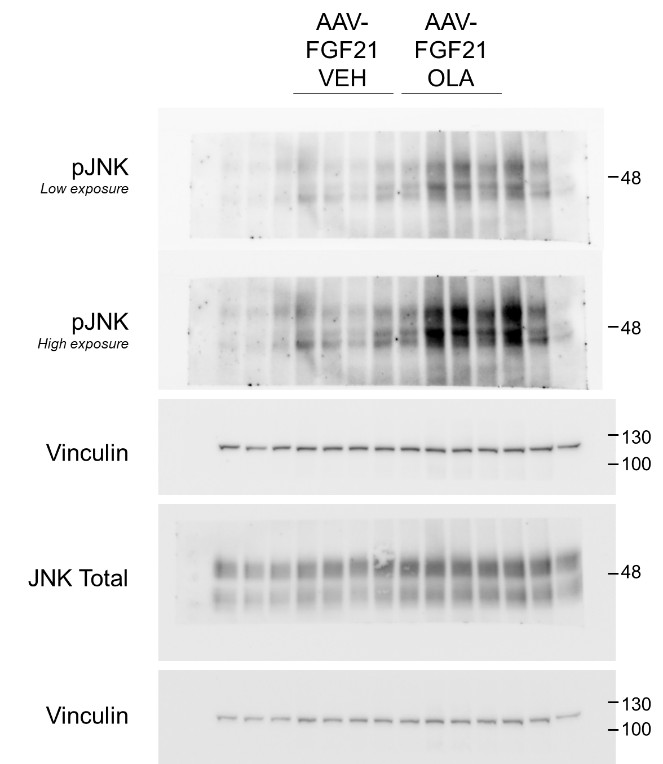

**Figure 6H**

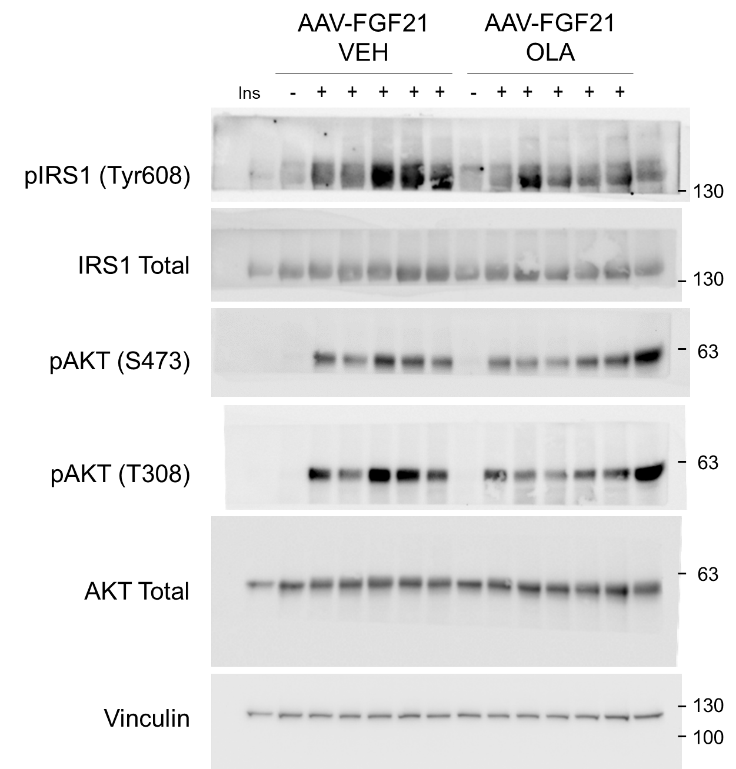

**Figure 6I**

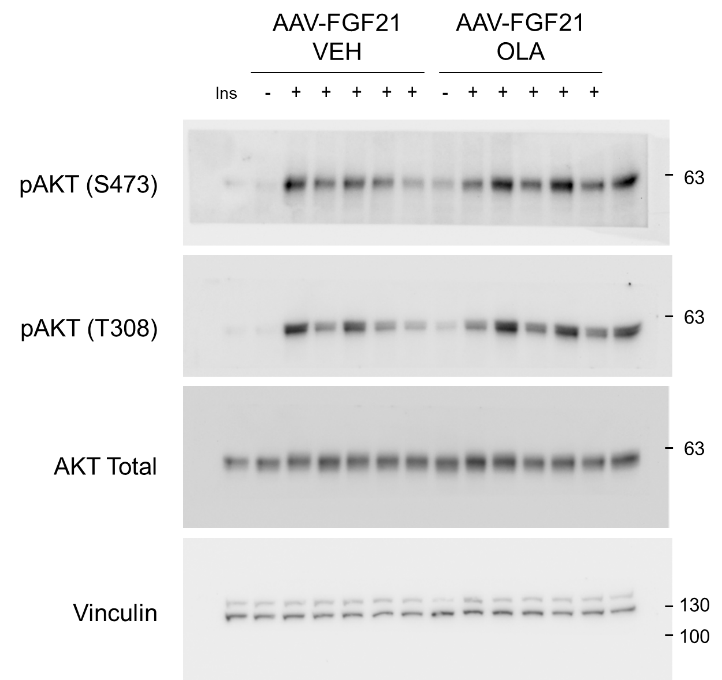

**Figure 7D**

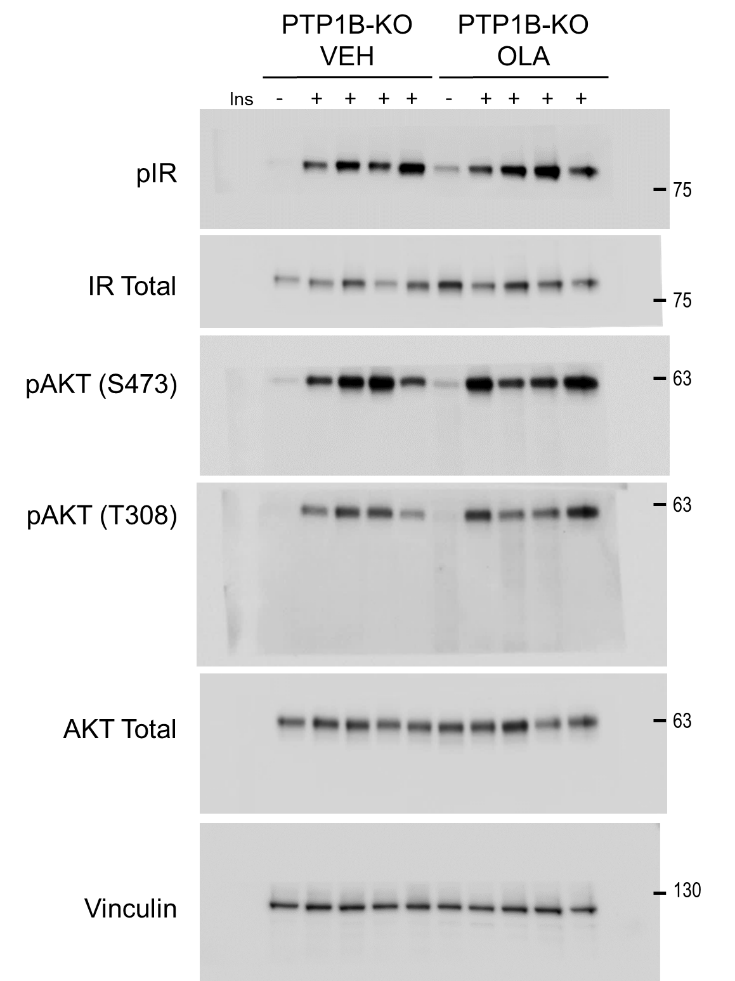

**Figure 7E**

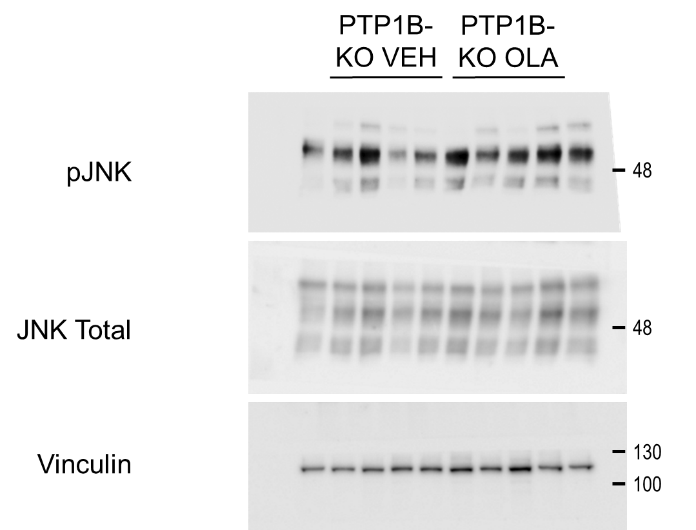

**Figure 7F**

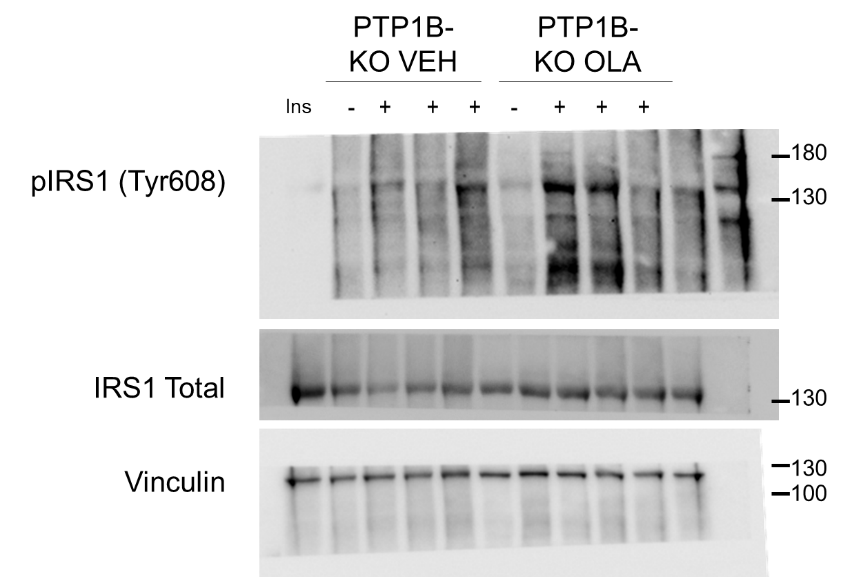

**Figure 7I**

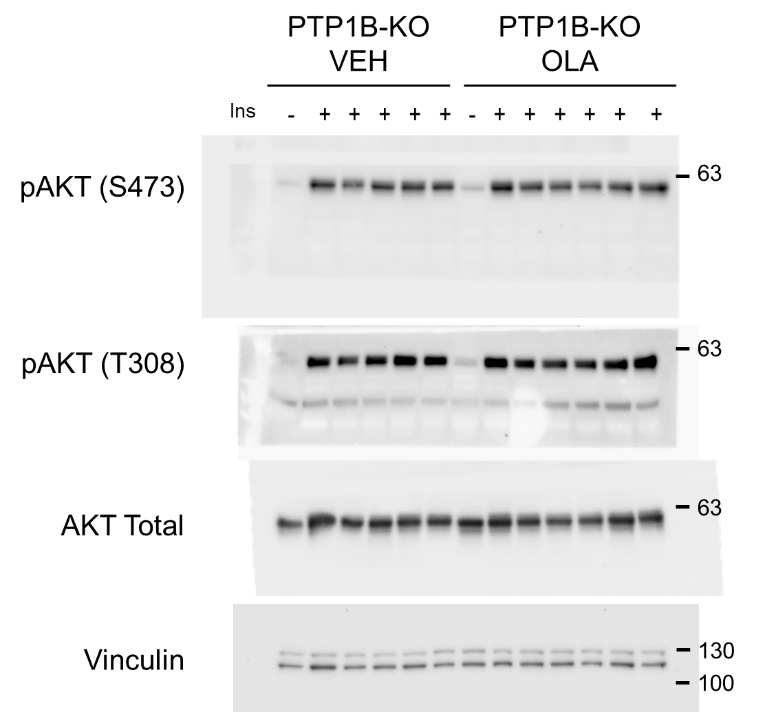

**Sup. Figure 2A**

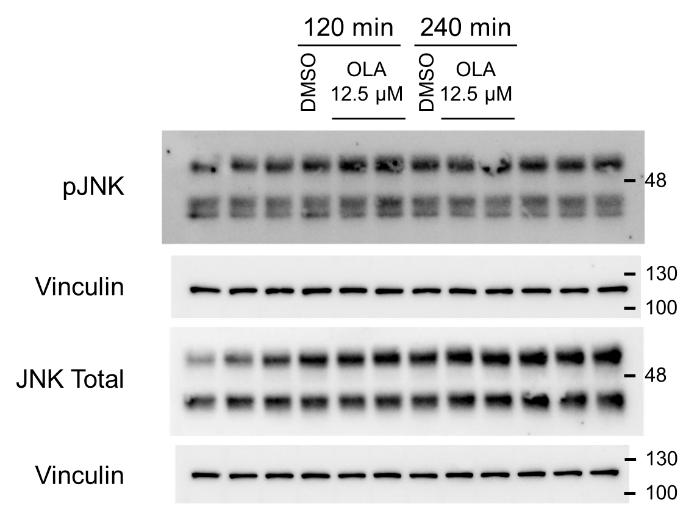

**Sup. Figure 3A**

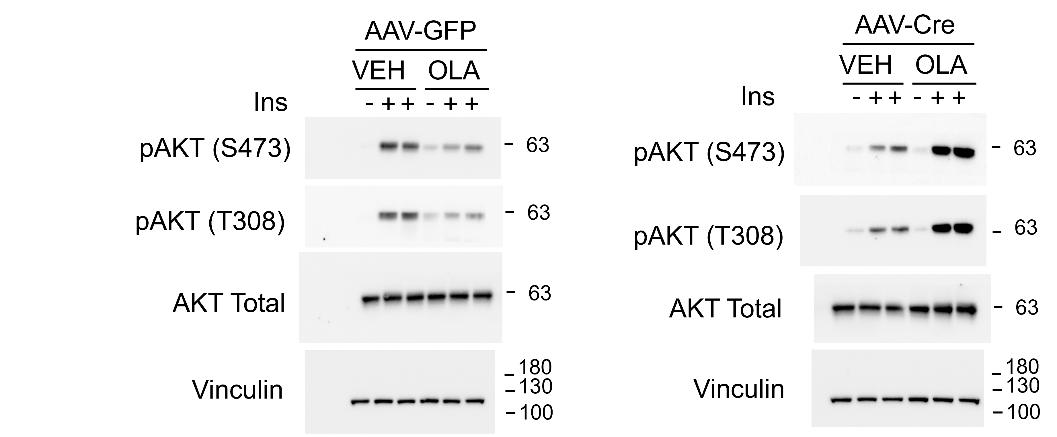

**Sup. Figure 4A**

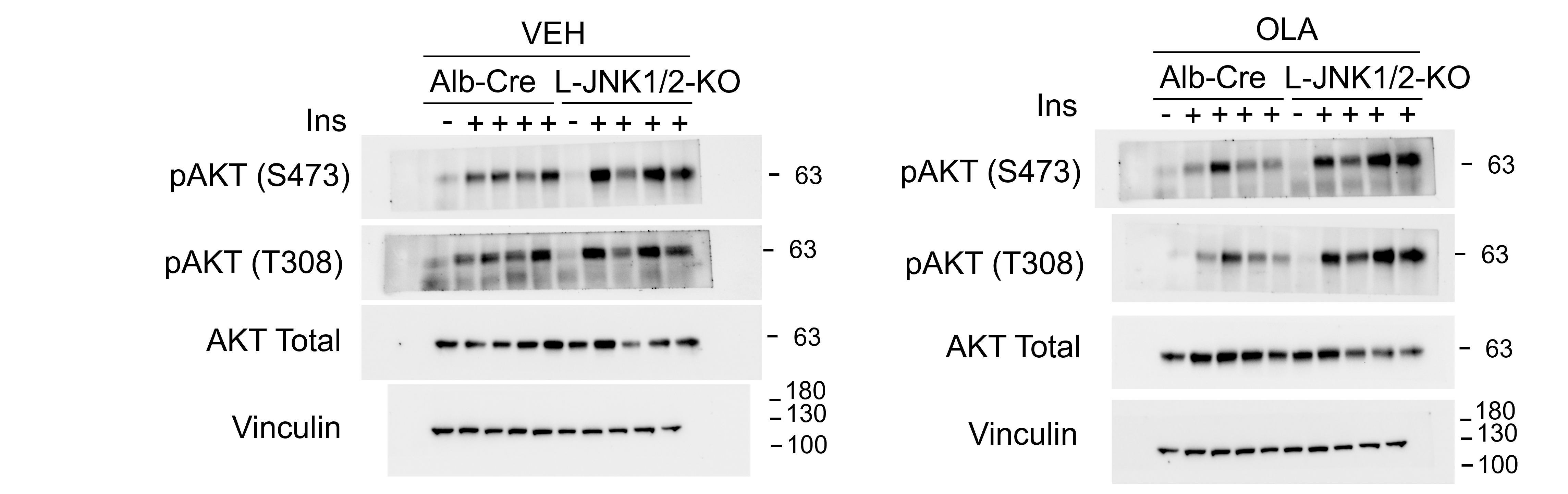

**Sup. Figure 4B**

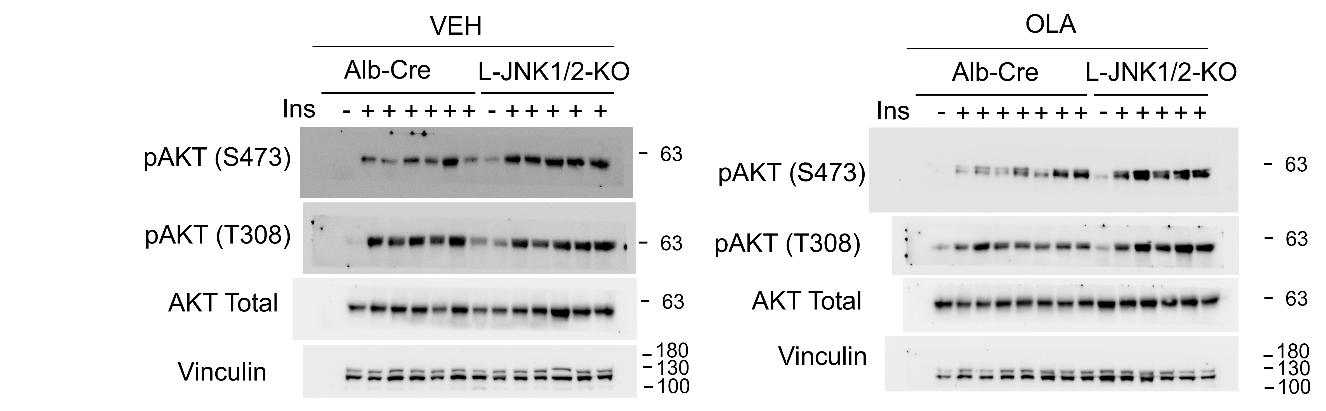

**Sup. Figure 5B**

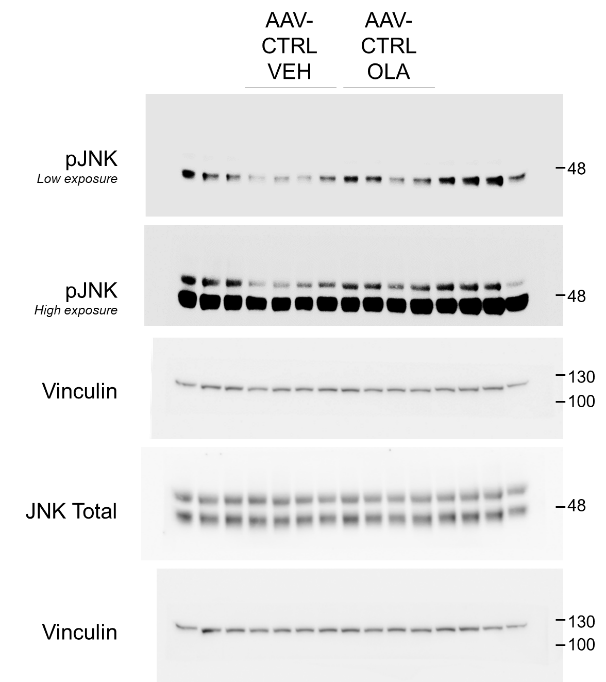

**Sup. Figure 5C**

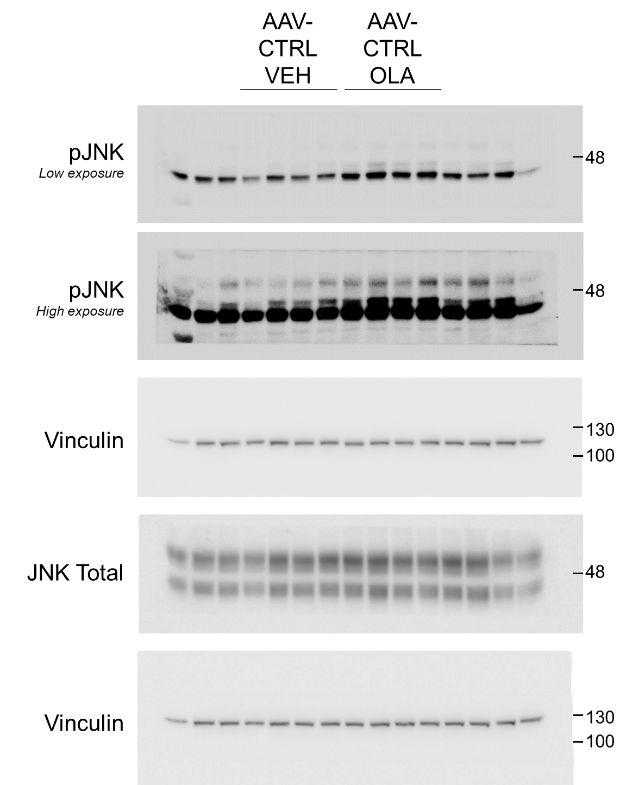

**Sup. Figure 5D**

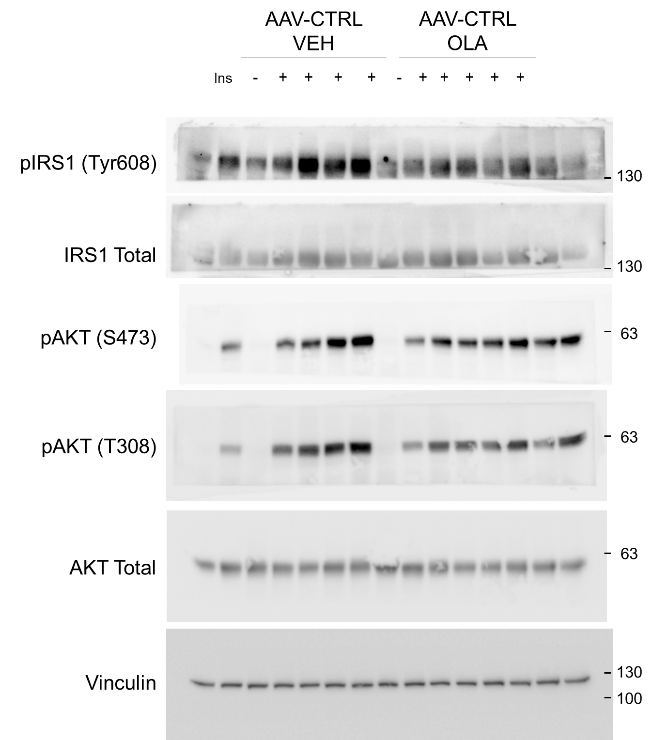

**Sup. Figure 5E**

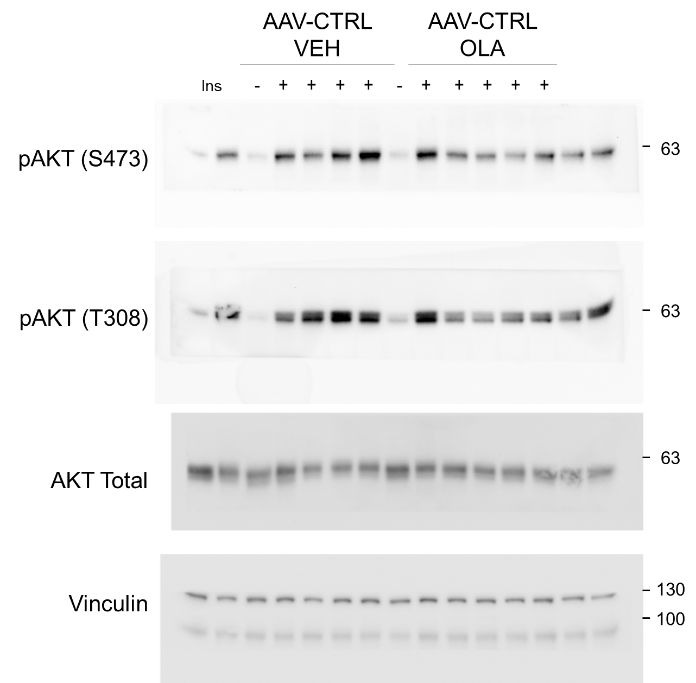
