## Supplementary material for "A hypothalamus-liver-skeletal muscle axis controlled by JNK1 and FGF21 mediates olanzapine-induced insulin resistance in an intraperitoneal treatment in male mice": Sup. Table 1

| **Supplementary Table 1 –** Antibodies and dilutions used for Western blot analysis | | |
| --- | --- | --- |
| **Antibody** | **Dilution** | **Reference, Company** |
| anti-phospho-IRβ/IGFRβ | 1:2000 | 3024S, Cell Signaling Technology |
| anti-IRβ Total | 1:2000 | 3025S, Cell Signaling Technology |
| anti-phospho AKT (S473) | 1:2000 | 4058S, Cell Signaling Technology |
| anti-phospho AKT (S308) | 1:2000 | 4056S, Cell Signaling Technology |
| anti-AKT Total | 1:2000 | 4691T, Cell Signaling Technology |
| anti-phospho IRS1 (Ser612)) | 1:2000 | 3203, Cell Signaling Technology |
| anti-phospho IRS1 (Tyr608) | 1:2000 | 09-432, Merck Millipore |
| anti- IRS1 Total | 1:2000 | 06-248, Merck Millipore |
| anti-Vinculin | 1:5000 | sc-7314, Santa Cruz Biotechnology |
| anti-phospho JNK | 1:2000 | 9251S, Cell Signaling Technology |
| anti-JNK | 1:2000 | sc-571, Santa Cruz Biotechnology |
| IgGκ-HRP Anti Mouse | | sc-516102, Santa Cruz Biotechnology |
| IgG-HRP Anti Rabbit | | A120-108P, Bethyl Laboratories |
