## Supplementary material for "A hypothalamus-liver-skeletal muscle axis controlled by JNK1 and FGF21 mediates olanzapine-induced insulin resistance in an intraperitoneal treatment in male mice": Sup. Table 2

| **Supplementary Table 2 –** Primers used for qRT-PCR. | |
| --- | --- |
| **primers** | **Sequence (5’-3’)** |
| *Hmox mouse - Forward* | *5’-CACAGATGGCGTCACTTCGTC-3’* |
| *Hmox mouse - Reverse* | *5´-GTGAGGACCCACTGGAGGAG-3´* |
| *Col1a1 mouse - Forward* | *5´-AATGGCACGGCTGTGTGCGA-3´* |
| *Col1a1 mouse - Reverse* | *5´-AGCACTCGCCCTCCCGTCTT-3´* |
| *Tgfb mouse - Forward* | *5´-TGCTAATGGTGGACCGCAACAAC-3´* |
| *Tgfb mouse - Reverse* | *5´-AGCTCTGCACGGGACAGCAAT-3´* |
| *Fgf21 mouse - Forward* | *5’-CTGCTGGGGGTCTACCAAG-3’* |
| *Fgf21 mouse - Reverse* | *5’- CTGCGCCTACCACTGTTCC-3’* |
| *Actb mouse - Forward* | *5’-CTCTGGCTCCTAGCACCATGAAGA-3’* |
| *Actb mouse - Reverse* | *5’-GTAAAACGCAGCTCAGTAACAGTCCG-3’* |
