## Supplementary material for "A hypothalamus-liver-skeletal muscle axis controlled by JNK1 and FGF21 mediates olanzapine-induced insulin resistance in an intraperitoneal treatment in male mice": Sup. Table 3

| **Supplementary Table 3 –** Antibodies and dilutions used for immunohistochemistry/immunofluorescence analysis | | |
| --- | --- | --- |
| **Antibody** | **Dilution** | **Reference, Company** |
| F4/80 | 1:100 | Kindly provided by A. Castrillo, CSIC, Spain |
| MRP14 | 1:100 | ab242945, Abcam |
| CD3 | 1:50 | Kindly provided by A. Castrillo, CSIC, Spain |
| Ly6C | 1:50 | Kindly provided by A. Castrillo, CSIC, Spain |
| 4-HNE | 1:200 | ab46545, Abcam |
| DAPI | 1:1000 | D1306, Invitrogen |
| Goat-anti-rabbit IgG (AF546) | | A11035, Invitrogen |
| Donkey-anti-rabbit IgG (AF555) | | A31572, Invitrogen |
| Goat-anti-rat IgG (AF546) | | A11081, Invitrogen |
| Biotinylated anti-Rabbit IgG (H+L) | | BA-1100, Vector Laboratories |
| Biotinylated anti-Rat IgG (H+L) | | BA-4000, Vector Laboratories |
